# Beyond Stickers and Spacers: Residue-level Contact Networks Predict IDR Organization and Phosphorylation Rewiring

**DOI:** 10.64898/2026.09.13.751321

**Authors:** Taseef Rahman, Vladimir N. Uversky

## Abstract

Intrinsically disordered regions (IDRs) act through ensembles of interconverting conformations, and the sticker and spacer framework describes how IDR sequences encode behaviour by sorting residues into two context-dependent classes. Coarse-grained simulations cover whole proteomes, yet force fields have been compared mainly on global dimensions rather than on residue-level contacts where stickers and spacers act. Here we turn simulations of thousands of IDRs with three force fields into contact networks and describe each residue by contact degree, how densely the residue is contacted, and how often the residue bridges distant parts of the chain. We show that residue types dominating these networks follow the parameters of each force field, whereas the positions of densely contacted and bridging residues are shared between force fields, set by sequence arrangement and learned by a model whose frozen predictions track experimental ensembles in blind tests. We then show that phosphorylation rewires the network most where the electrostatic potential of the unmodified ensemble fluctuates, in two force fields and beyond sequence charge descriptors, and that simulated contact changes track NMR shift changes. Contact degree from a simple model extends the sticker and spacer framework from two classes to a predictive, residue-level description of disordered proteins.

## 1 Introduction

Intrinsically disordered proteins and protein regions (IDRs) carry out many functions through ensembles of rapidly interconverting conformations rather than a single folded structure [1]. How the sequence of an IDR shapes the ensemble is largely organised by the sticker and spacer framework, in which a minority of residue types, such as aromatic residues and arginine, act as stickers whose number and spacing set the strength and the pattern of attractive contacts within single chains and in condensates, while the residues between stickers act as spacers [2, 1]. Coarse-grained force fields such as CALVADOS and Mpipi encode these attractions in residue-specific parameters and now simulate tens of thousands of sequences [3, 4, 5, 6, 7], and deep-learning models trained on these simulations predict global properties such as the radius of gyration directly from sequence [8, 6] or generate whole ensembles [9]. These force fields have, however, been compared mainly on global dimensions [8] or on the phase behaviour of a few sequences [10], rather than on residue-level contacts.

The sticker and spacer picture concerns precisely these contacts of individual residues [11, 2], which global dimensions do not capture. One way to resolve these contacts, borrowed from network science [12] and from the analysis of folded proteins [13, 14, 15], represents each snapshot of an ensemble as a graph whose nodes are residues and whose edges are contacts, computes a centrality such as betweenness for every residue and calls the most central residues, those through which the shortest connections between other residues pass, hubs of the contact network. The approach has only recently reached disordered proteins [16, 17, 18, 19] and to condensates [20, 21], where rankings of residue types by hubness are often read as statements about which residues act as stickers, although the sticker and spacer framework treats the division into stickers and spacers as dependent on context [22, 23, 24]. Whether such hubs reflect residue-type interactions, the arrangement of residues along the chain or choices made in the analysis, and whether force fields agree on which residues are hubs, have not been examined.

Here, we address this gap by comparing the contact networks of 10,281 CALVADOS-2, 10,266 Mpipi-Recharged and 908 Mpipi-GG ensembles of a common set of sequences, which is, to our knowledge, the first comparison of residue-level properties between force fields at this scale. We describe every residue by two readouts of the contact network, the mean number of contacts, or degree, and the hub occupancy, the fraction of snapshots in which the residue is a hub, after avoiding three analysis choices that otherwise make the network report chain position, sampling noise or residue size (Section 3.1). We first show that both readouts are reproducible although individual hubs are transient, and introduce a null model, neutral to residue identity and matched to the compaction of each ensemble, that separates chain geometry from residue-specific attraction. Against this null, hub occupancy splits into a residue-type part, which follows the interaction parameters of each force field, and an arrangement part, which the three force fields share and which a pre-registered comparison with shuffled and tyrosine-to-phenylalanine variants identifies as causal.

To identify the sequence features that set the arrangement part, we train a gradient-boosted model to predict the hub occupancy of every residue from residue identities, composition and chain position, which on held-out proteins comes close to the noise ceiling. Fixed before any shuffled sequence was labelled, the model predicts these sequences as accurately as natural ones, so arrangement rather than features peculiar to natural sequences drives the predictions. Trained on one force field, the model also predicts the others about as well as the force fields predict each other, as expected if the model captures the shared arrangement part. Indeed, a model that sees only the sticker and spacer surroundings of each residue recovers most of the full accuracy, and hubs sit preferentially in glycine-rich stretches, in line with the distinct roles of glycine and serine as spacers [25, 26]. Tested against experiment, predictions recorded before any label was computed explain hub occupancy beyond chain position in blind ensembles built from experimental data, and with degree, the more reproducible readout, the frozen predictions meet the pre-registered criteria of both blind tests. The same predictions anticipate how substitutions in designed variants move hubs in two force fields more accurately than a rule based on local stickiness and charge. Finally, the unmodified ensemble anticipates where phosphorylation rewires the contact network in direct simulations with two force fields, and the predicted change in contacts tracks the NMR chemical shifts of ten proteins, including *α*-synuclein. Taken together, these results show that the contact network extends the sticker and spacer framework from discrete residue classes to a continuous, context-dependent and predictive property of every residue.

## 2 Methods

### 2.1 Simulated ensembles

We used the BENDER set of 10,282 IDRs from bacteria, fungi, mammals, plants, protists and viruses [7]. Each sequence was simulated as a single chain at 300 K and an ionic strength of 150 mM with CALVADOS-2 [4] and with Mpipi-Recharged [27] in OpenMM [28], using a time step of 10 fs, a cubic box with a side of 50 nm (larger for chains of more than 300 residues) and run lengths that grow with chain length so that between 10,000 and 20,000 frames are saved. A subset of 908 sequences was also simulated with Mpipi-GG [8], a modified version of Mpipi [5], in LAMMPS [29]. The second half of every trajectory was treated as production. One truncated CALVADOS-2 run and 16 Mpipi-Recharged runs without frames leave 10,281 and 10,266 ensembles.

### 2.2 Contact networks, degree and hub occupancy

For every analysed snapshot we built a graph whose nodes are residues and in which two residues are connected when the two C*_α_* atoms are closer than 0.8 nm and the residues are at least three positions apart in sequence. We computed exact betweenness centrality with the algorithm of Brandes [12] and defined the hubs of a snapshot as the 10% of residues with the highest non-zero betweenness, breaking ties at random with a seed derived from the protein identifier so that no part of the chain is favoured. The hub occupancy of a residue, a residue-level counterpart of the persistent hubs described in folded proteins and condensates [30, 21], is the fraction of snapshots in which the residue is a hub, so that an average residue has an occupancy of 0.10 (Fig. 1d). The degree of a residue is the mean number of contacts per analysed snapshot.

**Figure 1:**
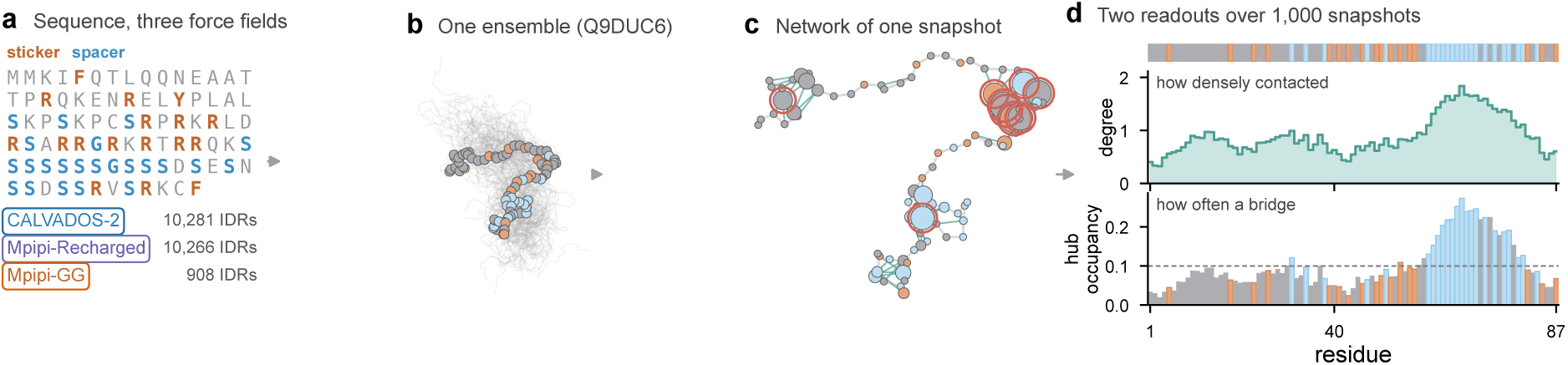
From ensembles to the two readouts of the contact network. (a) The sequence of Q9DUC6, with stickers in orange and spacers in blue, and the number of ensembles simulated with each force field. (b) The CALVADOS-2 ensemble of Q9DUC6 (87 residues), with 60 snapshots in grey aligned to one snapshot whose stickers F, W, Y and R are orange and whose glycine and serine are light blue. (c) The contact network of this snapshot, with contacts in green, node area proportional to betweenness and the 10% most central residues, the hubs, ringed. (d) Degree and hub occupancy of every residue over 1,000 snapshots, with the residue classes above and the mean occupancy dashed.

Labels were computed from 1,000 evenly spaced production snapshots, or from 500 and 250 snapshots for chains longer than 300 and 600 residues. We estimated the reliability of each label by correlating the values obtained from the even and the odd snapshots and applying the Spearman–Brown correction, and the square root of this reliability gives the noise ceiling, the highest correlation that a predictor can be expected to reach. To compare residue types, we also standardised time-averaged betweenness within each protein and removed the quadratic trend of this betweenness along the chain, which gives a position-adjusted *z*-score.

### 2.3 A residue-identity-neutral null model

To separate the contribution of chain geometry from that of residue-specific attraction, we built a residue-neutral null ensemble for every simulated ensemble. The null chains are self-avoiding C*_α_* chains with bonds of 3.8 Å and random dihedral angles whose beads repel as hard spheres of the bead diameters of the force field but never attract, so residues differ only in size. From a pool of 4,000 such chains spanning compact and expanded conformations, we matched every snapshot of the simulated ensemble with the chain of closest radius of gyration, so that the null ensemble has the same compaction but no residue-specific attraction. Null ensembles were analysed like the simulations (Supplementary Section S3).

### 2.4 Learning hub occupancy from sequence

Each residue was described by 487 features that encode the residues within ten positions on either side, the composition of local windows and of the whole sequence, global charge descriptors, the length of the chain and the position of the residue relative to the chain ends (Supplementary Section S5). We predicted hub occupancy with gradient-boosted regression trees and, for comparison, with ridge regression. The same trees trained on the position and length features alone serve as a positional baseline, and a consensus model was trained on the mean occupancy across force fields of every sequence simulated with more than one force field. To compare the contributions of residue type and of sticker and spacer context, we also trained the trees on chain position together with either the type of each residue or the stickiness, aromatic, glycine and serine content and charge of the surrounding residues (Supplementary Section S7).

### 2.5 Evaluation and predictions made in advance

Proteins were assigned to five cross-validation folds by a hash of the protein identifier, so that a protein falls in the same fold in every force field. For each held-out protein we computed the Spearman correlation between predicted and simulated occupancy over residues, together with the fraction of the simulated top 10% of residues that the predicted top 10% recovers, and we report the median correlation and the mean recovery over proteins. Transfer to a force field that was not used for training was evaluated on the same held-out folds. For the blind tests, which comprise 11 PED ensembles not used during development and the designed sequences, predictions were written to disk and the SHA-256 hashes of these files were recorded before any of the corresponding labels were computed, so that neither the models nor the predictions could be adjusted to the results, and we call such predictions *frozen*. We report for every external test both the Spearman correlation and the partial correlation that controls for the distance of a residue from the nearest chain end, always next to the residue-neutral null and to a baseline that uses chain position alone. We added the partial correlation as a co-primary metric after the first eight blind ensembles had shown how strong the positional baseline can be, and before the remaining three were scored.

### 2.6 Separating arrangement from interaction parameters

To separate arrangement from interaction parameters, we used two sets of CALVADOS-2 simulations run with the protocol of Section 2.1. The first contains three composition-preserving global shuffles of each of 597 BENDER sequences of 40 to 300 residues, stratified by kingdom and compaction. The second contains 800 BENDER sequences in which every tyrosine is replaced by phenylalanine, which lowers the CALVADOS-2 stickiness of the substituted residues from 0.98 to 0.87 while leaving residue size almost unchanged. To compare two occupancy profiles of equal length, we computed the per-protein Spearman correlation at matched chain positions, the partial correlation that removes the distance from the nearest chain end as in Section 2.5, and the correlation after additionally subtracting from each residue the mean position-adjusted occupancy of the same residue type in the respective label set, which removes the additive effect of residue identity. The hypotheses, statistics and decision rule of this comparison were recorded in a file that was hashed before any label of the shuffled or substituted sequences was examined (Supplementary Section S4). To relate hubs to the sticker and spacer framework, we called the 10% of residues of each protein with the highest hub occupancy the hubs of that protein and defined the enrichment of a set of residue types among hubs as the fraction of hubs that belong to the set divided by the fraction of all residues that do, pooled over proteins. We took F, W, Y and R as the canonical stickers and G and S as the canonical spacers [2, 1] and examined further sets in Supplementary Section S4.

### 2.7 Phosphorylation

We represented a phosphoserine, phosphothreonine or phosphotyrosine in CALVADOS-2 as the unmodified bead with a charge of −2 and in Mpipi-Recharged as a glutamate bead with doubled electrostatic amplitude, or for phosphotyrosine as the tyrosine bead with that electrostatic term. From an unmodified ensemble alone, we weighted each snapshot by exp(−Δ*U/k*_B_*T*), where Δ*U* is the energy change of phosphorylating the site in that snapshot, which in CALVADOS-2 reduces to the screened Coulomb term. The effective sample fraction of these weights measures how strongly the potential at the site fluctuates, and weighted averages estimate properties of the phosphorylated form. Rewiring was measured separately, in direct simulations of both forms. The rewiring *E*(*s*) of a site is the mean summed absolute difference of per-residue hub occupancy or degree between phosphorylated and unmodified replicates, minus the mean difference between unmodified replicates, and the rewiring *E_i_* of residue *i* is the same difference for the contact frequencies of that residue. Site selection, decision rules and analysis scripts were hashed before the corresponding simulations (Supplementary Section S17).

## 3 Results

### 3.1 Contact networks of simulated ensembles

We turned every snapshot of 10,281 CALVADOS-2, 908 Mpipi-GG and 10,266 Mpipi-Recharged ensembles into a contact network, in which residues are nodes and residues whose C*_α_* atoms lie within 0.8 nm of each other and at least three positions apart in sequence are connected, and described every residue by two readouts of the network (Fig. 1). The degree of a residue is the mean number of contacts per snapshot and reports how densely a stretch of the chain is contacted, whereas the hub occupancy of a residue is the fraction of snapshots in which the residue is among the 10% most central residues by betweenness and reports which residues connect distant parts of the chain. Three analysis choices that are common in network studies of disordered ensembles distort these readouts (Supplementary Section S2), since counting residues two positions apart as contacts lets chain position explain a median 94% of the variance of betweenness, fewer than about 430 snapshots leave per-residue betweenness too noisy to compare residue types, and contact cutoffs that scale with bead size make aromatic residues and arginine appear as hubs even in chains without residue-specific attraction (Supplementary Fig. S3a). We therefore require three positions of separation, analyse 1,000 snapshots and use a fixed cutoff throughout.

### 3.2 Individual hubs flicker, but the contact network is reproducible

Even with these choices fixed, the residues that act as hubs change from one snapshot to the next. In all three force fields, the hub sets of successive snapshots, which are separated by 50 to 100 ps, overlap with a median Jaccard index of 0.061 to 0.064, only modestly above the 0.042 to 0.051 expected for random sets of the same sizes (Fig. 2c). Nevertheless, hub occupancy computed from the even and the odd snapshots agrees with a median reliability, after Spearman–Brown correction, of 0.92 in CALVADOS-2, 0.96 in Mpipi-GG and 0.95 in Mpipi-Recharged (Fig. 2d).

**Figure 2:**
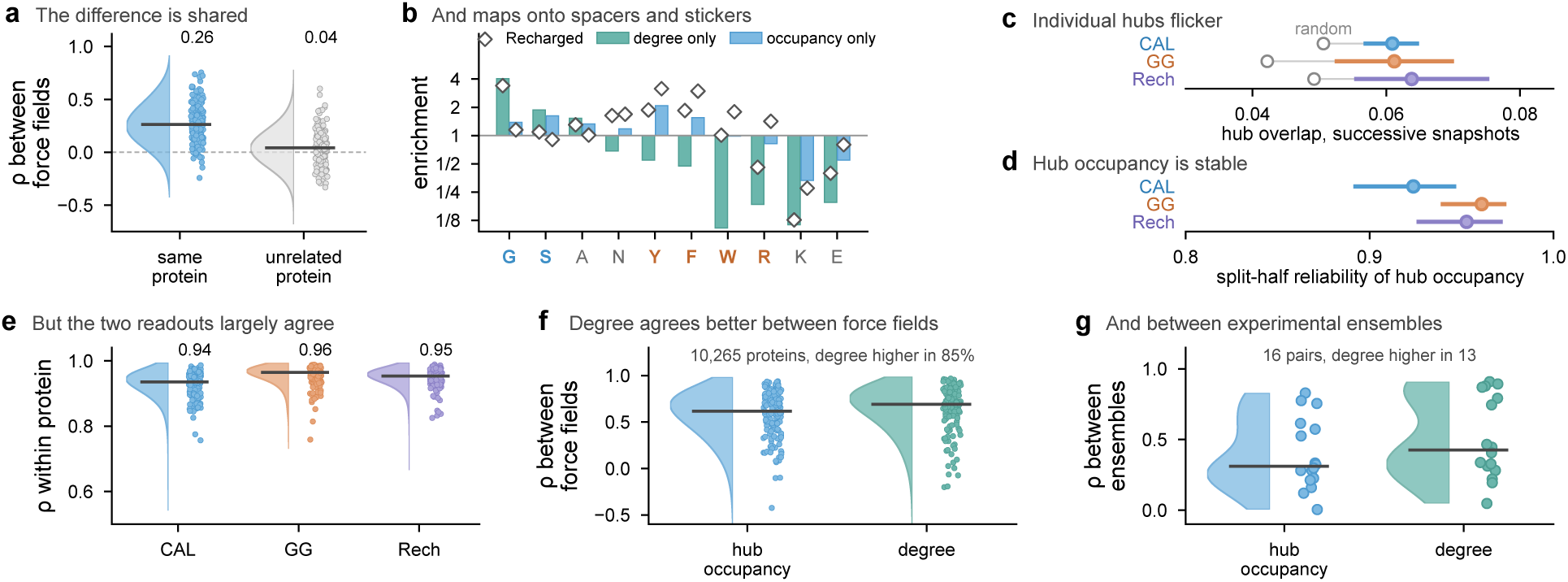
The contact network is reproducible, and degree more so than hub occupancy. (a) Spearman correlation between CALVADOS-2 and Mpipi-Recharged of the part of hub occupancy that degree does not explain, for the same protein and for an unrelated protein of the same length. (b) Enrichment of residue types among residues in the top 10% by only one readout, in CALVADOS-2 (bars) and Mpipi-Recharged (diamonds). (c, d) Median (dot) and interquartile range (line) over all ensembles of each force field of the overlap between the hub sets of successive snapshots, with random sets as open circles (c), and of the split-half reliability of hub occupancy (d). (e) Within-protein Spearman correlation of degree and hub occupancy. (f) Correlation of each readout between CALVADOS-2 and Mpipi-Recharged for 10,265 proteins. (g) Correlation of each readout between 16 pairs of independent experimental ensembles. Chain position is removed in (e) and (f).

The two readouts are closely related, since residues with many contacts tend to lie on many shortest paths. Within proteins, hub occupancy and degree correlate at 0.94 to 0.96 in the three force fields once chain position is removed (Fig. 2e). Nevertheless, the two differ in a reproducible way, as the part of hub occupancy that degree does not explain agrees between CALVADOS-2 and Mpipi-Recharged at a median of 0.26 against 0.04 for unrelated proteins of the same length (Fig. 2a). This part belongs to residues that bridge distant parts of the chain, among which aromatic residues are enriched, whereas residues of high degree but low occupancy are rich in glycine (Fig. 2b and Supplementary Section S16). Degree is the more reproducible readout, as the same protein agrees between CALVADOS-2 and Mpipi-Recharged at a median of 0.69 in degree but 0.62 in hub occupancy, and independent experimental ensembles agree at 0.43 against 0.31 (Fig. 2f,g), and we therefore report both readouts throughout.

### 3.3 Arrangement, not interaction parameters, sets the contact network

To isolate the part of hubness that comes from residue-specific interactions, we next compared each simulation with the matched null on the 907 sequences that were simulated with all three force fields (Supplementary Fig. S3b). In all three force fields, aromatic residues are more central than the null predicts and lysine is less central, but the residue-type profiles agree only moderately between the two model families, as the profile of CALVADOS-2 correlates at 0.52 with that of Mpipi-GG and at 0.36 with that of Mpipi-Recharged, whereas the two Mpipi models, which share one form of attraction, agree at 0.80. Each profile instead follows the interaction parameters of the corresponding force field more closely than those of the other family, correlating at 0.60 with the stickiness parameters *λ* of CALVADOS-2 and at 0.75 and 0.95 with the mean Wang–Frenkel attraction of each residue type in Mpipi-GG and Mpipi-Recharged (Fig. 3b–d, Supplementary Table S4). As a consequence, the canonical stickers F, W, Y and R are depleted among the hubs of CALVADOS-2 (enrichment 0.58) and enriched among those of Mpipi-Recharged (2.22), a reversal that holds for all seven sticker sets we tested (Supplementary Table S5). Whether stickers act as hubs in simulations therefore depends on the force field, in the same way that coarse-grained models disagree on which residue pairs stabilise condensates [10].

**Figure 3:**
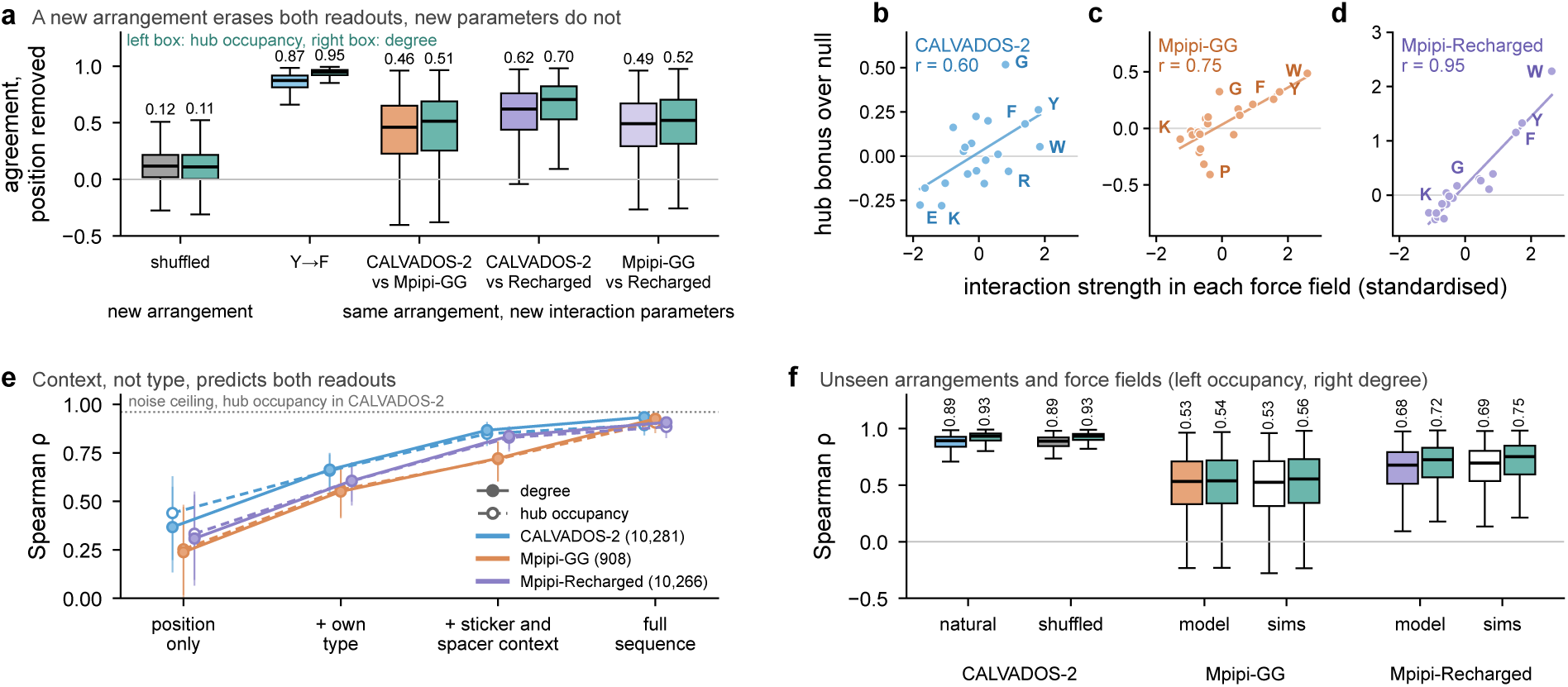
Arrangement, not interaction parameters, sets the contact network, and a sequence model learns this dependence. (a) Agreement between profiles of hub occupancy (left box) and degree (right box, teal) after removing chain position, for three shuffles (averaged) and the tyrosine-to-phenylalanine variant against the natural CALVADOS-2 profile, and between each pair of force fields over all proteins simulated with both. (b–d) Excess of each residue type over the residue-neutral null against the standardised interaction strength of that type, *λ* for CALVADOS-2 (b) and the mean Wang–Frenkel *ε* for Mpipi-GG (c) and Mpipi-Recharged (d). (e) Median and interquartile range of the held-out per-protein Spearman correlation of hub occupancy (dashed) and degree (solid) for gradient-boosted trees given chain position, residue type, sticker and spacer surroundings or all 487 sequence features, with the CALVADOS-2 noise ceiling of hub occupancy (dotted). (f) Models trained on CALVADOS-2 labels on held-out natural proteins, on 1,791 simulated shuffles and on the other two force fields (model), next to the agreement between CALVADOS-2 and each other force field (sims), for hub occupancy (left) and degree (right).

In contrast to the residue types, the positions of the hubs are largely shared. For the same sequences, the hub occupancy of CALVADOS-2 correlates with that of Mpipi-Recharged and Mpipi-GG with medians of 0.74 and 0.52. These agreements fall only to 0.64 and 0.45 when the distance from the chain ends and the average occupancy of each residue type in the respective force field are both removed, although the residue-type effects correlate between these force fields at only 0.12 and 0.42 (Supplementary Table S3). To test whether the shared part reflects arrangement, we compared CALVADOS-2 simulations of natural sequences with three composition-preserving shuffles of each and with variants in which every tyrosine became phenylalanine, under a decision rule fixed before any of these labels was examined, which required agreement with the natural profile below 0.2 for the shuffles and of at least 0.8 for the substitution and 0.4 for the other force fields once chain position is removed. In this comparison, the hub profile of a shuffled sequence agrees with the natural profile at a median of only 0.12. In contrast, keeping the residue order but altering residue interactions preserves much more of the natural profile. Replacing every tyrosine by phenylalanine, which CALVADOS-2 makes less sticky, gives a median agreement of 0.87, and simulating the same sequences with the differently parameterised Mpipi-GG and Mpipi-Recharged gives 0.43 and 0.63 (Fig. 3a), so all three thresholds are met. Degree passes the same rule, fixed again before any degree label of these variants was read, as shuffling erases degree as completely (0.11), whereas the substitution and the other force fields preserve even more of the profile (0.95, 0.44 and 0.72). In summary, the interaction parameters of a residue type set how often residues of that type are hubs, whereas the arrangement of the sequence sets which of these residues become hubs. Consistent with this division, residue identity explains only 22% of the within-protein variance of position-adjusted hub occupancy in natural sequences and 24% in the shuffled sequences.

### 3.4 A sequence model learns the arrangement-dependent map

We next asked whether hub occupancy can be predicted from sequence alone, as expected if arrangement sets where hubs form. Gradient-boosted trees trained on sequence features predict the CALVADOS-2 hub occupancy of held-out proteins with a median per-protein Spearman correlation of 0.89, close to the median noise ceiling of 0.96 (Fig. 3e). In addition, ridge regression on the same features reaches 0.84, which indicates that most of the signal lies in the features rather than the model (Supplementary Section S5). Degree is easier still to learn, as the same trees reach 0.93 on held-out proteins and transfer to Mpipi-Recharged at 0.72 (Fig. 3e,f), close to the agreement of 0.75 between the simulations, while chain position alone explains less of degree than of hub occupancy (0.37 against 0.44, Supplementary Section S16).

To test whether the model has learned how arrangement sets hubs rather than features peculiar to natural sequences, we turned to the shuffles, which share little of the natural hub profile. The model trained on CALVADOS-2 labels, fixed before any shuffle was labelled, predicts the 1,791 simulated shuffles with a median correlation of 0.89, as accurately as natural proteins, and the frozen degree model reaches 0.93 on the same shuffles (Fig. 3f), although the natural counterparts of these sequences were part of the training data. Moreover, although trained only on CALVADOS-2, the model predicts the hub occupancy of the same held-out proteins simulated with Mpipi-GG and Mpipi-Recharged with median correlations of 0.53 and 0.68, about as well as the CALVADOS-2 simulations predict the other two force fields (0.53 and 0.69).

We then asked whether this knowledge can be written in the language of stickers and spacers. Residue type alone falls short, as adding the type of each residue to chain position raises the median correlation only from 0.44 to 0.66. In contrast, a description of the sticker and spacer surroundings of each residue, by stickiness, aromatic, glycine and serine content and charge, reaches 0.85 (Fig. 3e) and thus recovers most of the accuracy of the full model. The same description reaches 0.83 in Mpipi-Recharged and, once proline is added, 0.84 in Mpipi-GG, which enlarges proline by a third [8] (Fig. 3e and Supplementary Table S7). For degree, the same description recovers a similar share of the full model but draws about twice as much on the glycine and serine content of the surroundings (Supplementary Section S16). Taken together, stickers and spacers predict hubs well once applied to the context of a residue rather than to the residue type, as the framework has long proposed [22, 23, 24].

To see which surroundings matter, we compared the neighbours of hubs with the neighbours of residues of the same type. Hubs have 15 to 30% more glycine among neighbouring residues in the three force fields and 63% more in the ensembles built from experimental data, an excess that residue-neutral nulls with the same bead sizes do not produce and that supports the view that glycine and serine are not equivalent spacers [25, 26]. Given this dependence on context, the model also predicts that the same substitution can raise or lower hub occupancy depending on the surrounding sequence, which designed variants test (Section 3.6).

Applied to designed sticker and spacer sequences and to all 28,058 IDRs of the human IDRome [6], the frozen consensus model places hubs on glycine-rich spacers next to aromatic stickers (Supplementary Section S14).

### 3.5 Predictions made in advance track experimental ensembles

To test the predictions against experiment, we turned to ensembles built from experimental data and deposited in the Protein Ensemble Database [31] (Fig. 4a). On 18 ensembles that we had used during development, predicted and ensemble occupancy correlate with a median of 0.46, compared with 0.31 for the residue-neutral null and 0.30 for chain position. We then assembled 11 further ensembles that had not been used before and recorded the model’s predictions before computing any of the labels. In this blind set the median correlation is 0.38, compared with 0.44 for the null and 0.46 for chain position, a strong baseline here because four of the 11 chains contain only 32 to 59 residues. We therefore controlled for the distance of each residue from the chain ends, which isolates the contribution of each predictor beyond chain position. In this comparison, the model reaches a median partial correlation of 0.38 on the development set and 0.22 on the blind set, where the partial correlation is positive in all 11 ensembles and above the null in nine (one-sided sign tests, *P* = 0.0005 and *P* = 0.03), whereas the null adds little (0.09 and 0.05, Supplementary Section S8). In a second blind set of 14 proteins, the model reaches a median of 0.31 against 0.03 for the null but beats the null in only 10 of the 14, short of the pre-registered criterion (*P* = 0.09, Supplementary Section S9). With degree in place of hub occupancy, in a reanalysis fixed before any degree label of these ensembles was computed, the frozen degree model meets both criteria, with median partial correlations of 0.31 on the blind set, positive and above the null in 10 of 11 ensembles, and 0.34 on the second blind set, above the null in 11 of 14 proteins (*P* = 0.03, Fig. 4a and Supplementary Section S16).

**Figure 4:**
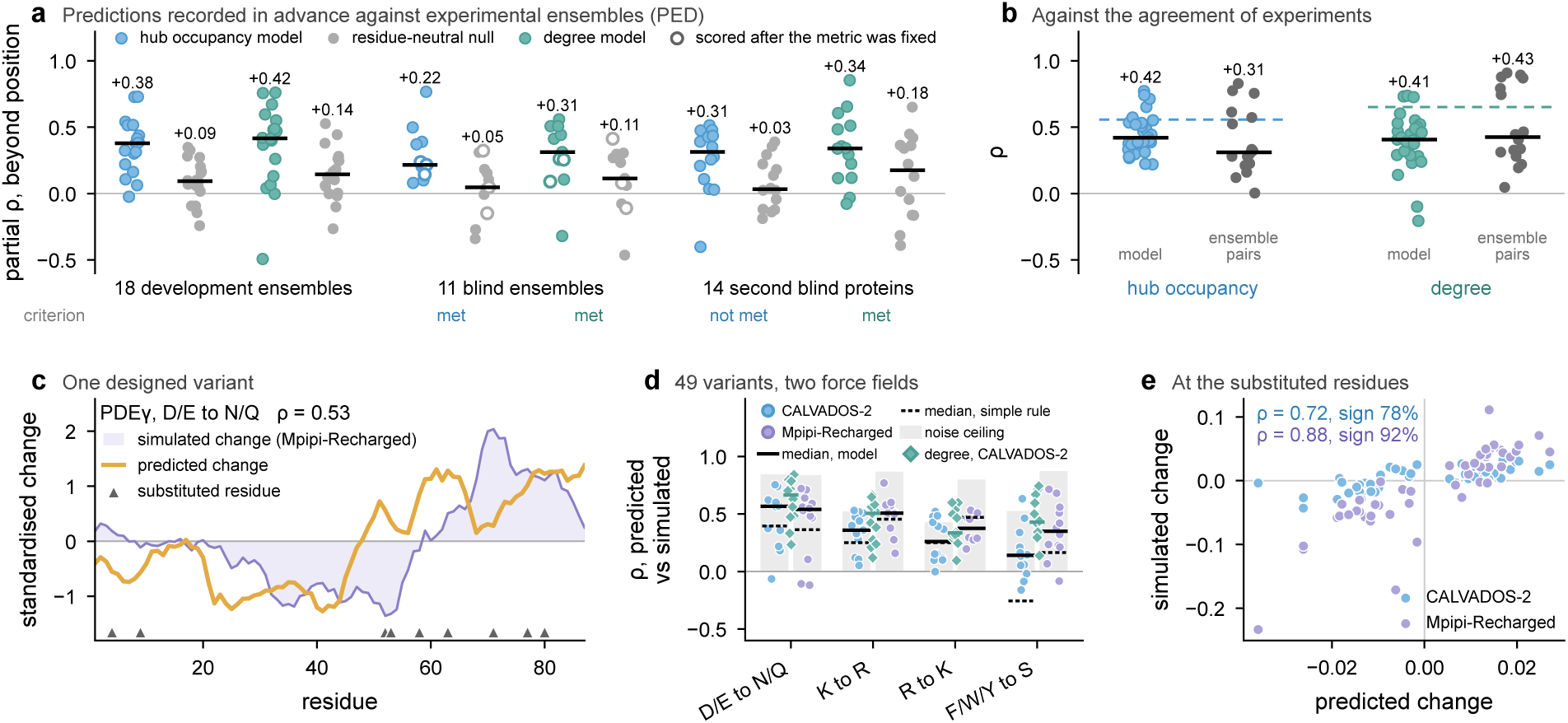
Predictions made in advance track experimental ensembles and designed substitutions. (a) Partial correlation, controlling for distance from the chain ends, between predictions recorded in advance and the ensemble labels for the hub occupancy model and the degree model, each next to its residue-neutral null, on 18 development ensembles, 11 blind ensembles (open symbols, scored after the partial correlation was fixed) and 14 proteins of a second blind set, with the verdict of each pre-registered criterion. Bars mark medians. (b) Spearman correlation of each model over the 29 development and blind ensembles next to the agreement between 16 pairs of independent ensembles, whose square root (dashed) approximates the reach of a perfect predictor. (c) Simulated (Mpipi-Recharged, shaded) and predicted change of hub occupancy for a PDE*γ* variant with neutralised acidic residues (triangles), standardised five-residue means. (d) Correlation of predicted and simulated change along the chain for the 49 substitution variants, with medians of the model (solid) and of a rule based on local stickiness and charge (dotted), the median noise ceiling (grey) and the frozen CALVADOS-2 degree model (teal diamonds, post hoc). (e) Mean predicted and simulated change at the substituted residues.

These correlations should be judged against the agreement between independent ensembles of drkN and of Sic1, which agree in hub occupancy at a median of only 0.31 over 16 pairs (Supplementary Table S11), so a perfect predictor would correlate with a single ensemble at only about 0.56 if every ensemble combined the same profile with independent errors, or about 0.65 for degree (Fig. 4b).

In the FUS N-terminal domain, which Supplementary Section S13 follows residue by residue, predicted degree agrees with the experimental profile at 0.74, against 0.54 for hub occupancy.

### 3.6 Predictions made in advance anticipate how substitutions move hubs

After recording the frozen predictions, we simulated the 22 experimental constructs and 75 variants of these constructs with CALVADOS-2 and Mpipi-Recharged (Fig. 4c–e, Supplementary Section S12). The predictions track the simulated hub occupancy of the constructs at a median Spearman correlation of 0.80 and 0.72 in the two force fields. For the 49 variants with substitutions, the predicted change of hub occupancy along the chain correlates with the simulated change at a median of 0.34 and 0.49, positive in 44 and 46 of the 49 variants (*P <* 10*^−^*^8^, pre-registered). Across variants, the predicted and simulated changes at the substituted residues correlate at 0.72 and 0.88. By contrast, a rule based only on local stickiness and charge reaches 0.24 and 0.43, below the model in both force fields (*P* = 0.0005 in CALVADOS-2), and the frozen degree model tracks the substitutions in CALVADOS-2 at 0.48, positive in all 49 variants (post hoc, Fig. 4d).

### 3.7 The contact network predicts how phosphorylation rewires disordered chains

Given that arrangement sets the contact network, we reasoned that the network should also register how a change of charge at one residue reorganises the chain, and that the unmodified ensemble should anticipate where. Phosphorylation adds two negative charges to a serine, threonine or tyrosine, which dominate the effect on chain dimensions [32, 33], and to first order the response of a network property is the covariance of that property with the electrostatic potential at the site, so sites where this potential fluctuates strongly can rewire the network most. We therefore simulated 20 proteins unmodified and with one of three sites phosphorylated, namely an annotated phosphosite and unannotated sites of matched and of low fluctuation, in three replicates each, and measured rewiring as the change in hub occupancy beyond the difference between unmodified replicates, under rules fixed before any simulation. Matched and annotated sites rewire the network more than low-fluctuation sites in 19 and 18 of 20 proteins (one-sided Wilcoxon *P* = 3 × 10*^−^*^6^ and 5×10*^−^*^5^), annotated sites rewire the network no more than matched ones (*P* = 0.43), and direct rewiring grows with the fluctuation of the unmodified runs across all 60 sites (*ρ* = −0.64 with the effective sample fraction, Fig. 5a,b). In Mpipi-Recharged, the same design replicates both contrasts (15 of 20 proteins, *P* = 0.006 and 0.002), and rewiring follows the fluctuation computed with that force field (*ρ* = −0.48, Supplementary Section S17). In both force fields, degree gives the same pattern with stronger contrasts.

**Figure 5:**
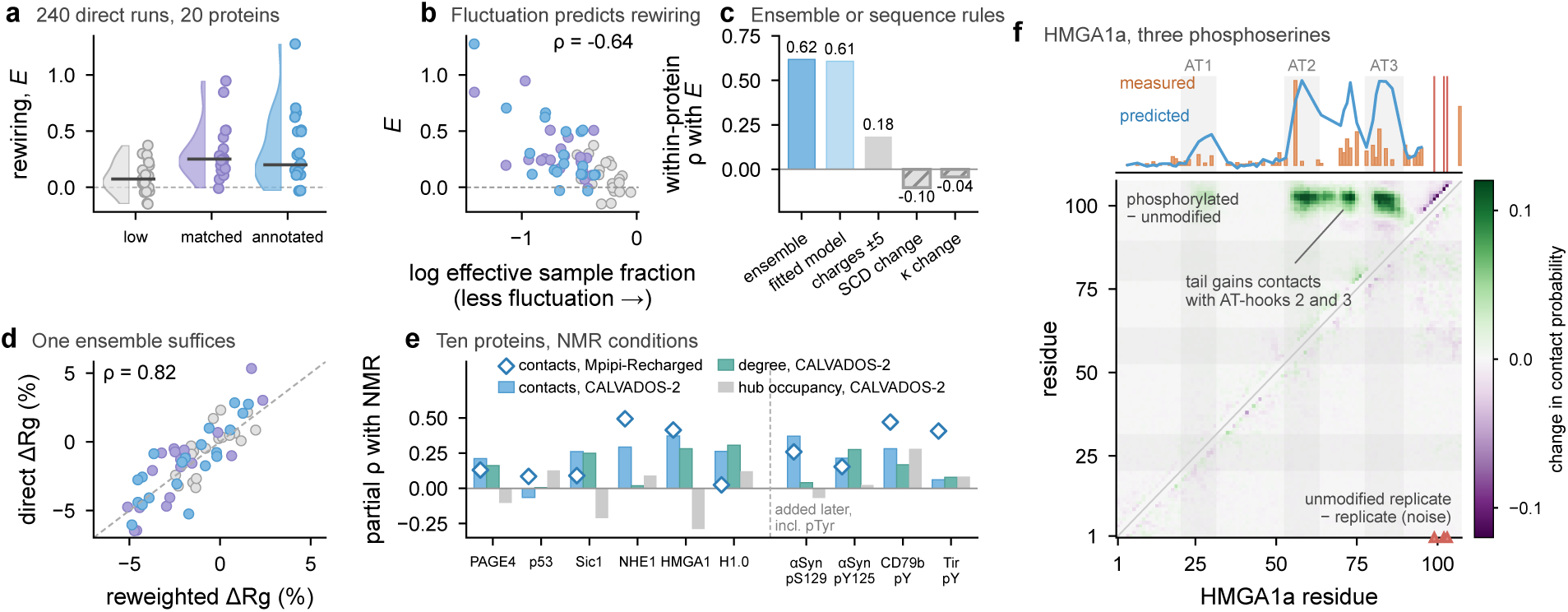
The contact network predicts how phosphorylation rewires disordered chains. (a) Rewiring *E* of hub occupancy beyond the difference between unmodified replicates for the low-fluctuation, matched and annotated site of 20 proteins in 240 CALVADOS-2 simulations, with sites as points and medians as lines. (b) *E* against the effective sample fraction of charge-model weights on the unmodified runs, which falls as the potential at the site fluctuates more. (c) Within-protein Spearman correlation of *E*, scaled by the replicate noise, with the ensemble fluctuation, a fitted sequence charge model, charged residues within five positions and the change in sequence charge decoration and in *κ* (hatched, post hoc). (d) Change in radius of gyration reweighted from the unmodified runs against the direct one. (e) Partial Spearman correlation of the simulated change of each residue with the measured amide shift change given the distance to the nearest phosphosite, for contacts (pre-registered), degree and hub occupancy in ten proteins in CALVADOS-2 (bars), with contacts in Mpipi-Recharged (diamonds) and the four pairs of a later plan right of the dashed line. (f) Change in contact probability between residues of HMGA1a on phosphorylation of S99, S102 and S103 (red) above the diagonal and between two unmodified replicates below it (gain green, loss purple), with measured amide shift changes (orange) and predicted rewiring (blue) above.

The environment of a site is only partly readable from sequence. A linear model of charge counts at increasing sequence separations, fitted to the fluctuation of 1,735 sites in other proteins, ranks the sites of a protein almost as well as the ensemble (*ρ* = 0.61 against 0.62 within proteins), yet the ensemble predicts rewiring beyond this model (*P* = 0.0007), whereas counts of charged residues within five positions reach 0.18 and the change in sequence charge decoration [34] or in *κ* [11] does not rank local rewiring at all (Fig. 5c). Reweighting the unmodified ensemble also reproduces the direct change in radius of gyration (*ρ* = 0.82 in CALVADOS-2 and 0.75 in Mpipi-Recharged, Fig. 5d), so one unmodified simulation gives the fluctuation that ranks single sites and the change in chain dimensions.

We then simulated disordered proteins with NMR shifts of both forms under each sample’s conditions and compared the predicted change in contacts of every residue with the change of the amide shifts of that residue, controlling for distance to the nearest site. In six proteins with phosphoserines or phosphothreonines, the predicted change tracks the measured shifts beyond distance in five (pre-registered, *P* = 0.031) and, with Mpipi-Recharged, in all six (*P* = 0.016). Four further pairs, fixed in a later plan, add *α*-synuclein phosphorylated at S129 or Y125 and two proteins with phosphotyrosines, and the prediction tracks the shifts in all four in both force fields (mean partial *ρ* = 0.23 and 0.32, *P* = 10*^−^*^4^). The change in degree does so in CALVADOS-2 but less in Mpipi-Recharged, and the change in hub occupancy does not (Fig. 5e). In HMGA1a, three phosphoserines in the acidic tail shift amides 20 to 45 residues away near the basic AT-hooks, where the network predicts that the tail gains contacts and rewires the chain most (Fig. 5f). The agreement is modest, but the contact network of a model with one bead per residue anticipates where phosphorylation reorganises disordered chains.

## 4 Discussion

In summary, once three analysis choices are avoided, the contact network separates into a residue-type part, which reads out the interaction parameters of each force field, and an arrangement part, which is shared between force fields, erased by shuffling and learnable from sequence, which suggests a division of labour between the sticker and spacer framework and the network view. Stickers and spacers describe which residue types attract and how the number and spacing of stickers control interactions [2, 1], whereas the contact network describes where contacts concentrate in a sequence and which residues bridge distant parts of the chain, a residue-level property set by arrangement, agrees between force fields and, unlike sequence-level patterning parameters [11, 34, 35] and window-based attraction maps [36], accounts for chain connectivity. Beyond where contacts sit, the network registers how a sequence responds to change, as phosphorylation rewires the network most where the potential at the site fluctuates in the unmodified ensemble, which sequence-based charge descriptors capture only in part. No parameter was fitted to phosphorylation, although sequence models trained on such simulations could extend these predictions to whole proteomes. Degree agrees better with experimental ensembles, whereas hub occupancy adds the bridging role of aromatic residues. With the larger weight of glycine and serine in models of degree, this suggests that spacers set how densely a stretch is contacted and stickers set which residues connect distant parts of the chain.

The main limitation is that the three force fields share one bead per residue, an implicit solvent and isotropic pair interactions, so the shared arrangement part alone cannot show that real proteins behave alike, although all-atom force fields agree on hub positions similarly (Supplementary Section S9). Experimental ensembles and NMR shifts provide independent tests but are few and agree only moderately, especially where residual helices form (Supplementary Sections S8 and S12), and the phosphate is represented coarsely. Finally, all ensembles are single chains in dilute solution, and network features failed three pre-registered tests of phase behaviour (Supplementary Section S15).

## Acknowledgements

We thank Kresten Lindorff-Larsen for helpful discussions, and we acknowledge the GAIVI research computing cluster of the University of South Florida for computing resources.

## Supplementary Material

### S1 Simulation data, trajectory readers and reproducibility of the labels

Table S1 summarises the simulated ensembles, and Fig. 1 shows how a hub occupancy label is obtained from the snapshots of one ensemble. CALVADOS-2 and Mpipi-Recharged trajectories were written by OpenMM in DCD format and Mpipi-GG trajectories by LAMMPS in compressed XTC format. We read both formats with dependency-free readers, and the radii of gyration that our XTC reader obtains agree with those reported by LAMMPS to within 0.001 Å. In addition, these readers reject files that hold fewer frames than the file header declares, so one CALVADOS-2 trajectory (Q3UHK0) has no labels. Labels for ensembles that remain on a computing cluster are computed there with the same code. To confirm that both routes agree, we recomputed on the cluster the labels of 523 Mpipi-Recharged residues and 416 Mpipi-GG residues, which were identical in every column to those computed on a workstation.

**Table S1:** Simulated ensembles used for training and evaluation.

| Force field | Engine and protocol | Sequences | Labels |
| --- | --- | --- | --- |
| CALVADOS-2 [4] | OpenMM, 300 K, 150 mM, 10 fs, 10,000 to 20,000 frames | 10,282 | 10,281 |
| Mpipi-Recharged [27] | OpenMM, same protocol as CALVADOS-2 | 10,282 | 10,266 |
| Mpipi-GG [8] | LAMMPS, 300 K, 150 mM, 20 fs | 908 | 908 |

### S2 Analysis choices that manufacture hubs

Network analyses of disordered ensembles differ in which residue pairs count as contacts, in how many snapshots are analysed and in whether the contact cutoff depends on residue size, and these rarely examined choices decide which residues appear as hubs. For example, applied to the same CALVADOS-2 simulations, a rule that counts residues two positions apart ranks glutamate, aspartate and arginine as hubs and the aromatic residues as anti-hubs (Supplementary Fig. S1b), whereas a cutoff scaled to bead size ranks tryptophan, arginine, tyrosine and phenylalanine as hubs and glycine, alanine and serine as anti-hubs even in residue-neutral chains without any attraction (Supplementary Fig. S3a). Depending on the analysis, the same simulations thus suggest either that stickers are hubs or that stickers are not hubs. However, the first ranking follows from chain position and the second from bead size. In addition, the few tens to few hundreds of snapshots that typical analyses use leave enough sampling noise to blur the differences between residue types under any rule. Below, we derive each of these effects in closed form (Supplementary Fig. S2). Throughout this work, we avoid these effects by requiring contacts to span at least three positions under a fixed cutoff, by analysing 1,000 snapshots per ensemble and by comparing each ensemble with a matched residue-neutral null, as described in Section 2.

#### The three choices in the simulations

Section 3.1 of the main text summarises the following analysis. We first examined three common analysis choices that decide which residues appear as hubs. The first is whether residues two positions apart count as a contact. If such pairs count, residues *i* and *i* + 2, whose C*_α_* atoms can never be more than 0.76 nm apart, are connected in every snapshot, and the shortest paths of the contact graph run along this ladder. If these contacts were the only edges, the graph would split into two paths through the even and the odd residues, on which the residue at index *j* of a path of *m* residues lies on exactly *j*(*m* − 1 − *j*) shortest paths (Supplementary Section S2). Betweenness then rises steeply from the chain ends towards the middle of the chain (Supplementary Fig. S1a). Indeed, in 1,198 CALVADOS-2 ensembles analysed with this rule and 200 snapshots, the mean *z*-scores of glutamate, aspartate and arginine are positive (0.07, 0.05 and 0.06) while those of phenylalanine, tryptophan and tyrosine are negative (−0.14, −0.13 and −0.06), which gives a ranking in which charged residues act as hubs and aromatic residues as anti-hubs (Supplementary Fig. S1b). This closed form explains a median 94% of the variance of time-averaged betweenness under this rule, compared with 41% when contacts require three positions of separation, and once averaged over the positions that each residue type occupies, predicts the ranking with a correlation of 0.99 across the 20 residue types. Once the positional trend is removed, however, every residue type falls within 0.07 of zero. The ranking therefore reflects where residue types sit along the chain.

The second choice is the number of snapshots, which matters because the betweenness of individual residues fluctuates strongly from one snapshot to the next. In 100 ensembles simulated with CALVADOS-2, with a median length of 75 residues, residue values averaged over two disjoint random sets of snapshots correlate with medians of 0.22 for 25 snapshots, 0.32 for 50, 0.47 for 100, 0.65 for 200 and 0.81 for 500 snapshots (Supplementary Fig. S1c), which projects to a reliability of 0.90 for 1,000 snapshots. These values follow the Spearman–Brown relation for independent snapshots with a single-snapshot reliability of only 0.009, which implies that about 430 snapshots are needed for a reliability of 0.8 and about 970 for 0.9 (Supplementary Section S2). Noise of this size blurs the differences between residue types, and we therefore compute all labels from 1,000 snapshots.

The third choice concerns the contact cutoff, which contact maps of coarse-grained simulations of disordered proteins and of condensates often scale with the bead diameters of the two residues [37, 38, 10], because residues in these models differ in size. In residue-neutral null ensembles matched to 908 CALVADOS-2 ensembles, a size-scaled rule calibrated to give the same mean number of contacts as the fixed cutoff makes position-adjusted hubness an almost linear function of bead diameter (*r* = 0.98), with tryptophan, tyrosine, phenylalanine and arginine appearing as hubs and glycine, alanine and serine as anti-hubs (Supplementary Fig. S3a). This is the familiar division into aromatic and arginine stickers and glycine and serine spacers, produced here by residue size alone in chains without any residue-specific attraction, and under this rule all 20 residue types depart significantly from the average (Supplementary Section S2). In contrast, with the fixed cutoff of 0.8 nm the same null ensembles give residue-type means only between −0.10 and 0.19. The restraint-free starting pools deposited in the Protein Ensemble Database for *α*-synuclein, Sic1 and drkN show the same size-dependent pattern under the scaled rule (Supplementary Section S3), which confirms that the bias arises from chain geometry and bead size alone. Taken together, residue rankings obtained with size-scaled rules largely report residue size, and we therefore use the fixed cutoff throughout.

**Figure S1:**
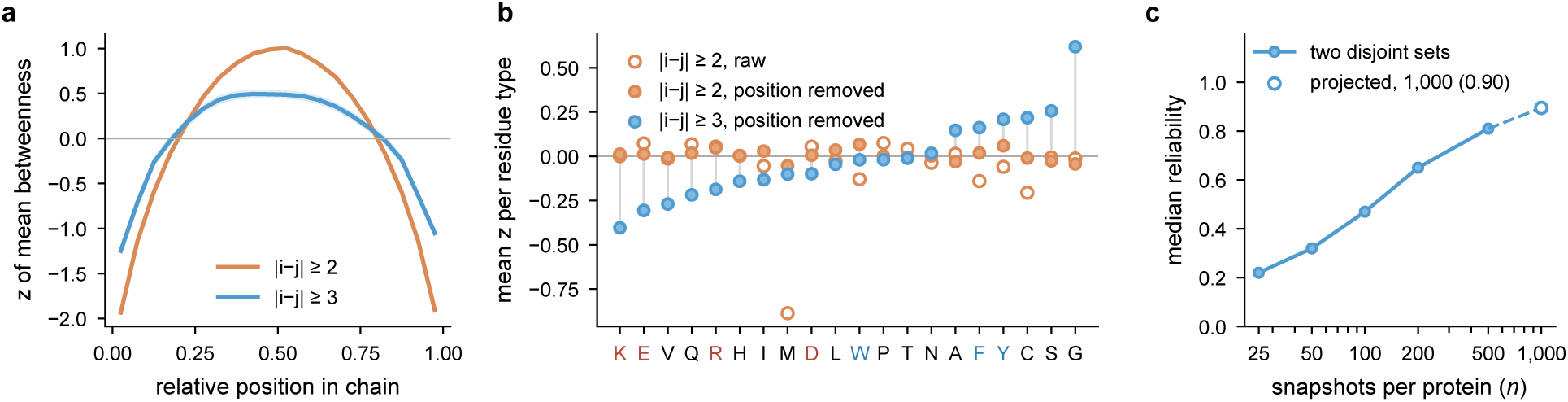
Analysis choices that manufacture hubs. (a) Within-protein *z*-score of time-averaged betweenness along the chain in 1,198 CALVADOS-2 ensembles, with contacts requiring at least two (orange) or three (blue) positions of separation (bands, 95% confidence intervals). (b) Mean *z*-score per residue type under the two-position rule before (open) and after (filled orange) removal of the positional trend, and under the three-position rule after removal (blue). (c) Median split-half correlation of per-residue betweenness from *n* snapshots in 100 ensembles, with the Spearman–Brown projection to 1,000 snapshots.

#### Contacts between near neighbours

Consecutive C*_α_* atoms are 0.38 nm apart, so residues *i* and *i* + 2 are never more than 0.76 nm apart and are in contact in every snapshot whenever the contact rule admits residues two positions apart. If these were the only contacts, the contact graph would be the union of two paths, one through the residues with even and one through those with odd index. On a path every pair of nodes is joined by exactly one shortest path, and the node at index *j* of a path of *m* nodes, counted from zero, lies on the shortest paths of the *j*(*m* − 1 − *j*) pairs that this node separates, so the betweenness of the node is proportional to *j*(*m* − 1 − *j*), a parabola that vanishes at the ends of the path and peaks at the centre. Residue *i* of a chain of *N* residues has *j* = ⌊*i/*2⌋ on a path of about *N/*2 nodes, so in this limit betweenness depends on chain position alone. Although additional contacts add shortcuts, in the 1,198 CALVADOS-2 ensembles of Supplementary Fig. S1a the closed form still explains a median 94% of the variance of time-averaged betweenness under this rule, compared with 41% when contacts require three positions of separation, a rule under which no contact is guaranteed because residues *i* and *i* + 3 can be up to 1.14 nm apart. Averaging the standardised closed form over the positions that each residue type occupies predicts the mean *z*-score of the 20 residue types under the near-neighbour rule with a correlation of 0.99 (Spearman 0.94), whereas the same average correlates with the ranking obtained under the stricter rule at only 0.33. The ranking in which charged residues appear as hubs and aromatic residues as anti-hubs therefore follows from where these residues sit along the chain. We note that several published network analyses of disordered ensembles connect residues that are one or two positions apart or do not report excluding such pairs [16, 18, 19], and widely used network tools keep such contacts by default [39, 40, 41], although other studies and tools exclude such pairs [14, 42]. More generally, that the backbone alone shapes the statistics of protein contact networks was noted earlier for folded proteins [43].

#### Sampling

If snapshots are independent, the reliability of a residue value averaged over *n* snapshots follows the Spearman–Brown relation *R*(*n*) = *nr*_1_*/*(1+(*n*−1)*r*_1_), where *r*_1_ is the reliability of a single snapshot. The five median split-half reliabilities of Supplementary Fig. S1c each imply a single-snapshot reliability between 0.0085 and 0.011, and the single value *r*_1_ = 0.0092 reproduces these reliabilities with predicted values of 0.19, 0.32, 0.48, 0.65 and 0.82 against observed values of 0.22, 0.32, 0.47, 0.65 and 0.81, which indicates that the analysed snapshots behave as independent samples. Solving for *n* gives *n^∗^* = *R^∗^*(1−*r*_1_)*/*(*r*_1_(1−*R^∗^*)), so about 430 snapshots are needed for a reliability of 0.8, about 970 for 0.9 and about 2,050 for 0.95. In a related approach, resampling tests of the convergence of network properties, as implemented in PyInKnife2 [15], address the same question for ensembles of folded proteins.

#### Contact cutoffs scaled to residue size

To quantify this bias, we averaged the position-adjusted *z*-score of each residue type within every protein of the residue-neutral null ensembles matched to 908 CALVADOS-2 ensembles and tested these protein means against zero with Wilcoxon signed-rank tests, correcting for the 20 tests. Under the size-scaled rule all 20 residue types depart significantly from zero, 17 of the 20 by at least 0.2 standard deviations, with tryptophan (1.42), arginine (1.04), tyrosine (0.92) and phenylalanine (0.77) as the most central and glycine (−1.19), alanine (−0.79) and serine (−0.68) as the least central types. In contrast, under the fixed cutoff the protein means of all types lie between −0.12 and 0.29, and only methionine, which mostly occupies the first position, exceeds 0.2. Eleven of these small departures are nevertheless statistically detectable with 908 ensembles. We therefore compare every simulation with the matched null of the same ensemble rather than with zero.

### S3 Validation of the residue-neutral null model

The chains of the null model have bonds of 3.8 Å, uniformly random dihedral angles and virtual bond angles drawn from one of three ranges (85 to 150, 100 to 150 or 115 to 155 degrees, each used for a third of the pool, so that the pool spans compact and expanded chains). Non-bonded beads repel each other as hard spheres at a distance of 0.8 *σ_ij_*, where *σ_ij_* is the mean bead diameter of the two residues in the force field. For Mpipi-GG, whose pair sizes are not additive for proline, *σ_ij_* is the pair size defined by the force field. In every case, growth backtracks locally whenever a new bead clashes. For the experimental ensembles, the null occupancy is the average over five independent pools, since a single pool gives noisy values for individual residues.

As an independent check of the null model, we used the restraint-free conformer pools that IDPConformerGenerator [44] produced for *α*-synuclein, Sic1 and drkN before experimental restraints were applied, which the Protein Ensemble Database deposits together with the restrained ensembles. These all-atom pools are a residue-neutral reference with realistic local structure. Under the size-scaled contact rule, the pools and our null model agree on the direction and approximate size of the residue-class biases (Table S2), for example placing glycine, serine and alanine between −0.6 and −1.0 in all three proteins. In contrast, under the fixed cutoff the class biases of both the pools and our null model remain small, within ±0.5. The null values also differ by 0.1 to 0.3 between independent pools, so the two do not agree in detail. We note that the values in Table S2 come from a single null pool.

**Figure S2:**
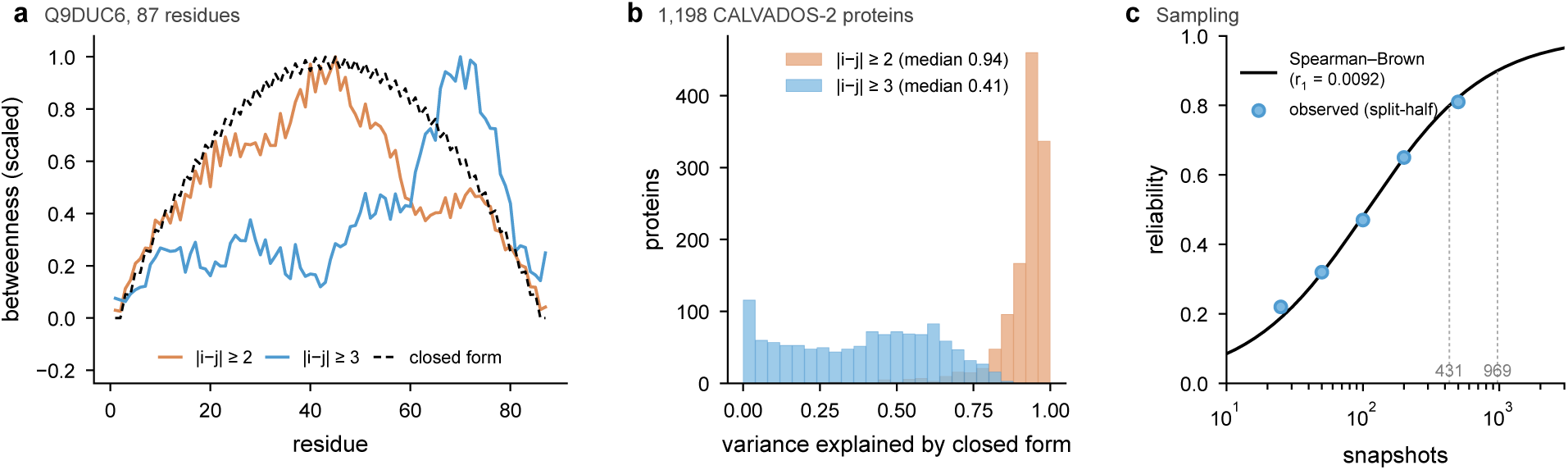
Closed forms for the analysis choices of Section 3.1. (a) Time-averaged betweenness along the chain of the protein of Fig. 1 under the two-position (orange) and three-position (blue) rules, each scaled to a maximum of one, with the closed form *j*(*m* − 1 − *j*) of the two-path limit (dashed). (b) Variance of time-averaged betweenness explained by the closed form in 1,198 CALVADOS-2 proteins under the two rules. (c) Reliability of per-residue betweenness against the number of snapshots, with the Spearman–Brown curve for a single-snapshot reliability of 0.009, the observed split-half medians and the numbers of snapshots needed for reliabilities of 0.8 and 0.9 (dotted).

**Figure S3:**
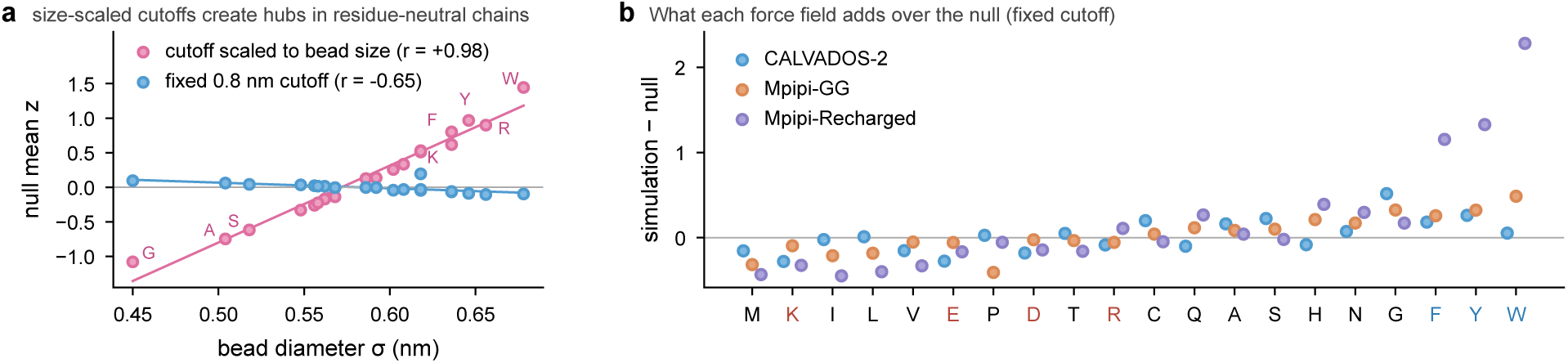
Residue-neutral nulls. (a) Mean position-adjusted *z*-score per residue type in residue-neutral nulls against bead diameter, for a size-scaled and a fixed 0.8 nm cutoff. (b) Excess of each force field over the matched null per residue type under the fixed cutoff, on 907 sequences simulated with all three.

### S4 Arrangement and interaction parameters

The shuffled sequences are three composition-preserving global permutations of each of 597 BENDER sequences. These sequences were chosen in proportion to the kingdoms of the dataset, in equal thirds across three ranges of the scaling exponent and with lengths between 40 and 300 residues, and were simulated with CALVADOS-2 under the protocol used for all other ensembles. All 1,791 trajectories are complete, and 590 of the natural sequences have labels, the remaining seven having left the dataset in a later revision. Every shuffle has exactly the composition of the natural sequence and shares a median of 11% of residues with the natural sequence at matched positions. We read the shuffled sequences from the topology file of each trajectory, because a sequence file stored separately from the shuffles describes different permutations. Similarly, the substituted sequences replace every tyrosine of 800 BENDER sequences by phenylalanine, which lowers the CALVADOS-2 stickiness of the substituted residues from 0.98 to 0.87 and the bead diameter from 0.646 to 0.636 nm, and 772 of the corresponding natural sequences have labels.

The hypotheses, statistics and decision rule of the comparison were written down in a file that was hashed before any label of these simulations was examined and after predictions for these simulations had been recorded. However, the predictions for the shuffles had been computed from that separate sequence file, and therefore from permutations that were not simulated. We thus recomputed the predictions from the simulated sequences with the same hash-verified models, which are fixed and so cannot be tuned to the labels. The decision rule required, after removing chain position, a median agreement with the natural profile below 0.2 for the shuffles, of at least 0.8 for the tyrosine-to-phenylalanine variants and of at least 0.4 for the other force fields, and all three were met (Table S3).

**Table S2:** Mean position-adjusted *z*-score per residue class in the restraint-free starting pools (pool) and in our null model built for the restrained ensemble of the same protein (null).

| Rule | Protein | Aromatic (FWY)<br>pool / null | Acidic (DE)<br>pool / null | Basic (KR)<br>pool / null | GSA<br>pool / null |
| --- | --- | --- | --- | --- | --- |
| fixed | α-synuclein | -0.27 / -0.08 | -0.27 / -0.00 | +0.34 / +0.03 | -0.04 / -0.03 |
| fixed | Sic1 | +0.27 / -0.24 | - | -0.14 / -0.17 | +0.18 / +0.11 |
| fixed | drkN | +0.29 / -0.17 | -0.13 / -0.21 | +0.07 / -0.48 | -0.11 / -0.04 |
| size | α-synuclein | +0.43 / +0.94 | -0.11 / +0.14 | +1.13 / +1.09 | -0.70 / -0.87 |
| size | Sic1 | +1.29 / +0.95 | - | +0.81 / +1.14 | -0.61 / -0.85 |
| size | drkN | +0.84 / +0.68 | -0.19 / -0.39 | +0.58 / +0.30 | -0.74 / -0.99 |

Table S3 lists all agreements between occupancy profiles, and on the subsets of the shuffled proteins that were also simulated with Mpipi-Recharged (589) and Mpipi-GG (48), the position-removed agreements of CALVADOS-2 with these force fields are 0.63 and 0.43, close to those on all proteins (0.62 and 0.46, Fig. 3a). Shuffling leaves the residue-type part almost unchanged, as residue identity explains 22.0% of the variance of position-adjusted occupancy in the natural sequences and 23.9% in the shuffled sequences, a difference whose 95% bootstrap interval over proteins ranges from 0.9 to 2.7 percentage points. Likewise, the enrichment of aromatic residues among hubs is 0.75 in the natural sequences and 0.78 in the shuffled sequences (95% bootstrap interval of the difference −0.17 to 0.11), and that of glycine and serine is 2.80 and 2.84 (−0.11 to 0.03). Natural arrangements are nevertheless more structured than random ones. Compared with the mean of the three shuffles of each sequence, natural sequences have a larger standard deviation of position-adjusted occupancy, a higher lag-one autocorrelation of this occupancy along the chain and a higher maximum occupancy (two-sided Wilcoxon tests, *P <* 10*^−^*^7^ in each case), and natural sequences exceed all three shuffles in 36%, 35% and 34% of proteins, compared with 25% if natural and shuffled arrangements were exchangeable. In contrast, the changes of occupancy produced by the substitution of tyrosine by phenylalanine are too small to be resolved for individual residues, with a mean change of −0.003 at the substituted positions, and the model, fixed in advance, does not predict these changes (Spearman correlation 0.04).

**Table S3:** Agreement between hub occupancy profiles at matched positions (median per-protein Spearman correlation), raw, after removing the distance from the chain ends and after additionally removing the mean position-adjusted occupancy of each residue type. The upper block compares the CALVADOS-2 profiles of natural sequences with changes of arrangement or of interaction parameters, and the lower block compares the force fields on the 907 sequences simulated with all three, with the correlation between the 20 residue-type means of each pair in the last column.

| Comparison | $n$ | Raw | Position removed | Identity also removed | Type means |
| --- | --- | --- | --- | --- | --- |
| Natural versus shuffled, CALVADOS-2 | 590 | 0.28 | 0.12 | 0.11 |  |
| Natural versus tyrosine to phenylalanine | 772 | 0.91 | 0.87 | 0.80 |  |
| CALVADOS-2 versus Mpipi-GG | 908 | 0.53 | 0.46 | 0.44 |  |
| CALVADOS-2 versus Mpipi-Recharged | 10,265 | 0.69 | 0.62 | 0.59 |  |
| CALVADOS-2 versus Mpipi-GG | 907 | 0.52 | 0.46 | 0.45 | 0.42 |
| CALVADOS-2 versus Mpipi-Recharged | 907 | 0.74 | 0.67 | 0.64 | 0.12 |
| Mpipi-GG versus Mpipi-Recharged | 907 | 0.53 | 0.49 | 0.47 | 0.35 |

To test whether the reversal depends on which residues count as stickers, we repeated the enrichment of stickers and spacers among hubs for further definitions (Table S5 and Supplementary Fig. S4). Besides the canonical stickers F, W, Y and R, we used the aromatic residues with histidine instead of arginine, the aromatic residues alone, tyrosine with phenylalanine as in prion-like domains [2], tyrosine with arginine as in the FUS family [25], and the canonical set with histidine or lysine added, and besides glycine and serine we used the polar spacers G, S, N and Q. The own sets of each force field take the residue types that the force field makes most attractive, ranked by interaction parameter as W Y F R G L C I M H S N Q T P A V K D E for CALVADOS-2, W Y F H Q N R G P S C A M L V I T E D K for Mpipi-GG and W Y F H R Q N G P C S E A D M T L V I K for Mpipi-Recharged. Each of these seven sticker sets is depleted among the hubs of CALVADOS-2 and enriched among those of Mpipi-Recharged, and so are the four residue types that each force field makes most attractive, which are W, Y, F and R in CALVADOS-2 and W, Y, F and H in both Mpipi models. In contrast, sets that contain glycine, which CALVADOS-2 makes nearly as attractive as arginine, are enriched among the hubs of CALVADOS-2, so that the five, six and ten residue types that CALVADOS-2 makes most attractive are enriched by 1.94, 1.74 and 1.48 times. Measured against the residue-neutral null, the four most attractive residue types of each force field are more central than geometry predicts in both Mpipi models but not in CALVADOS-2, where arginine falls slightly below the null, whereas the aromatic residues alone exceed the null in all three force fields. Finally, in the ensembles built from experimental data, the sticker sets are enriched among hubs by 1.27 to 1.60 times, with intervals that include 1 for three of the seven sets, and glycine and serine are not enriched.

**Table S4:** Pearson correlation across the 20 residue types between the excess of each force field over the matched residue-neutral null on the 907 sequences simulated with all three force fields and three interaction scales, the stickiness parameters *λ* of CALVADOS-2 and the mean Wang–Frenkel attraction *ε* of each residue type over all partner types in Mpipi-GG, read from the Mpipi-GG simulation input, and in Mpipi-Recharged, read from the parameter file distributed with OpenMpipi [27]. The scale of CALVADOS-2 correlates with those of Mpipi-GG and Mpipi-Recharged at 0.78 and 0.73, and the two Mpipi scales correlate with each other at 0.99.

| Excess of | CALVADOS-2 $\lambda$ | Mpipi-GG $\varepsilon$ | Mpipi-Recharged $\varepsilon$ |
| --- | --- | --- | --- |
| CALVADOS-2 | 0.60 | 0.36 | 0.28 |
| Mpipi-GG | 0.50 | 0.75 | 0.74 |
| Mpipi-Recharged | 0.69 | 0.94 | 0.95 |

### S5 Predictor details

Each residue is described by 487 features, which comprise a one-hot encoding of the residues within ten positions on either side, the amino-acid composition of windows of ±5 and ±15 residues, the composition of the whole sequence together with the fraction of charged residues, the net charge per residue, the sequence charge decoration and the logarithm of the sequence length, and three features that describe the position of the residue relative to the chain ends. The gradient-boosted trees are HistGradientBoostingRegressor models of scikit-learn with 300 iterations, 63 leaves, a learning rate of 0.08 and an L2 penalty of 1.0. For comparison, the same trees reach a median of only 0.71 when trained and scored on the position-adjusted *z*-score of time-averaged betweenness instead of hub occupancy. Supplementary Fig. S5 shows the prediction for a held-out protein of median accuracy and the predictions recorded in advance for single residues placed at the centre of four hosts, which the design batch tests.

We also trained the predictor on several force fields at once, without telling the model which force field a label comes from (Table S6). Trained on two force fields and scored on held-out proteins of the third, with five folds by protein, the model reaches medians of 0.61, 0.54 and 0.64 for CALVADOS-2, Mpipi-GG and Mpipi-Recharged. Trained on all three with each force field weighted equally, the pooled model predicts held-out proteins of each force field at 0.75, 0.77 and 0.80. The consensus model, trained on the mean occupancy over force fields for the 10,266 sequences labelled in at least two, predicts the held-out consensus label at 0.90 and the labels of CALVADOS-2 and Mpipi-Recharged at 0.79 and 0.85, values that exceed the agreement between these two simulations largely because the consensus label averages the two. The agreement between CALVADOS-2 and Mpipi-Recharged, which bounds any predictor that ignores the force field, has a median of 0.69 over 10,265 sequences, or 0.62 after removing chain position, varies little between organism groups (0.66 to 0.72) and is lower for chains of at least 400 residues (0.55) and for compact chains with *ν* below 0.40 (0.57, Supplementary Fig. S6).

**Figure S4:**
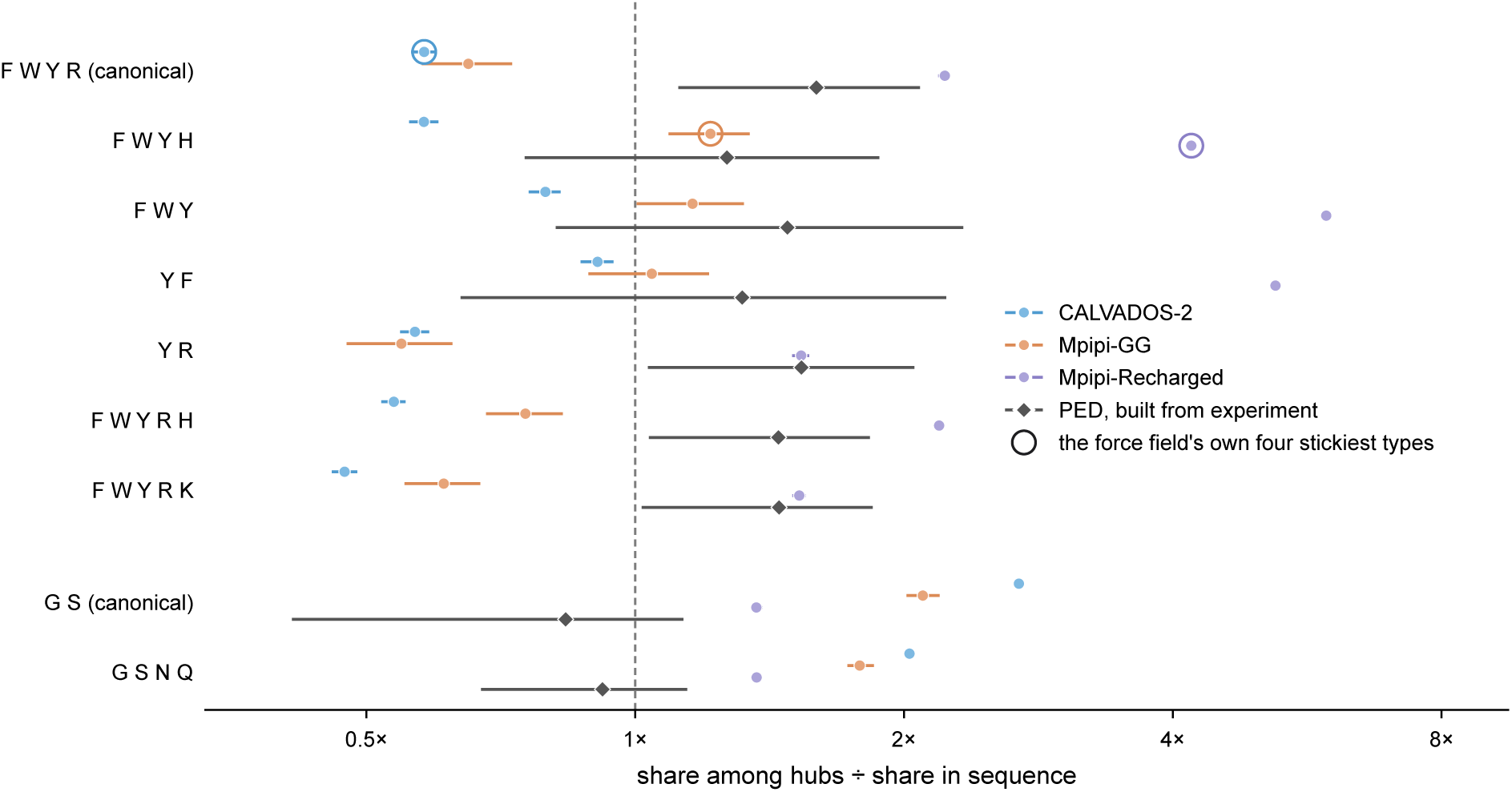
Enrichment of sticker and spacer sets among hubs under different definitions. Hubs are the 10% of residues of each protein with the highest hub occupancy, and the enrichment of a set is the share of hubs that belong to the set divided by the share of all residues that do, with 95% bootstrap intervals over proteins. Rings mark the four residue types that each force field makes most attractive, W, Y, F and R in CALVADOS-2 and W, Y, F and H in both Mpipi models. The values are those of Supplementary Table S5.

### S6 Hub occupancy across taxa

The BENDER set spans bacteria, fungi, mammals, plants, protists and viruses [7], so we asked whether the learned map depends on the group of origin, in a descriptive analysis specified before the models below were trained (Supplementary Fig. S7). In the five-fold cross-validation of Section 2.5, the full model reaches median per-protein correlations of 0.88 to 0.91 in every group, close to noise ceilings of 0.96 to 0.97, whereas chain position alone reaches 0.38 to 0.48. To test transfer between groups directly, we trained the same model on five groups and scored the sixth, which lowers the median correlation by at most 0.006, to between 0.87 and 0.90, so the map learned from one set of organisms applies to sequences from another. Transfer of the CALVADOS-2 model to Mpipi-Recharged ranges from 0.63 in fungi to 0.71 in bacteria for both training schemes and follows the agreement between the two simulations of each group, which ranges from 0.66 to 0.72. Finally, the reversal of sticker enrichment between force fields holds in every group, as the canonical stickers F, W, Y and R are depleted among the hubs of CALVADOS-2, with enrichments of 0.44 to 0.90, and enriched among those of Mpipi-Recharged, with enrichments of 1.95 to 2.75, whereas glycine and serine are enriched among hubs in both force fields and every group.

### S7 Sticker and spacer context

The context models use the folds and settings of the full model and replace the 487 sequence features by the chain position and length features together with a short description of each residue and the surrounding residues. This description comprises the stickiness and charge of the central residue, the mean stickiness, the fractions of aromatic residues (F, W and Y) and of glycine and serine and the net charge in windows of ±2, ±5 and ±10 residues, and the mean stickiness, aromatic fraction and net charge per residue of the whole sequence. Stickiness is the *λ* of CALVADOS-2 or the mean Wang–Frenkel attraction of each residue type in Mpipi-GG or Mpipi-Recharged (Supplementary Table S4), and charges are −1 for D and E, +1 for K and R and 0 otherwise. Each context model uses either the scale of the force field whose labels the model learns or the scale of the other model family, and the proline extension adds whether a residue is a proline and the proline fraction of the same windows. Table S7 and Supplementary Fig. S8 give the medians of the held-out per-protein Spearman correlations.

**Figure S5:**
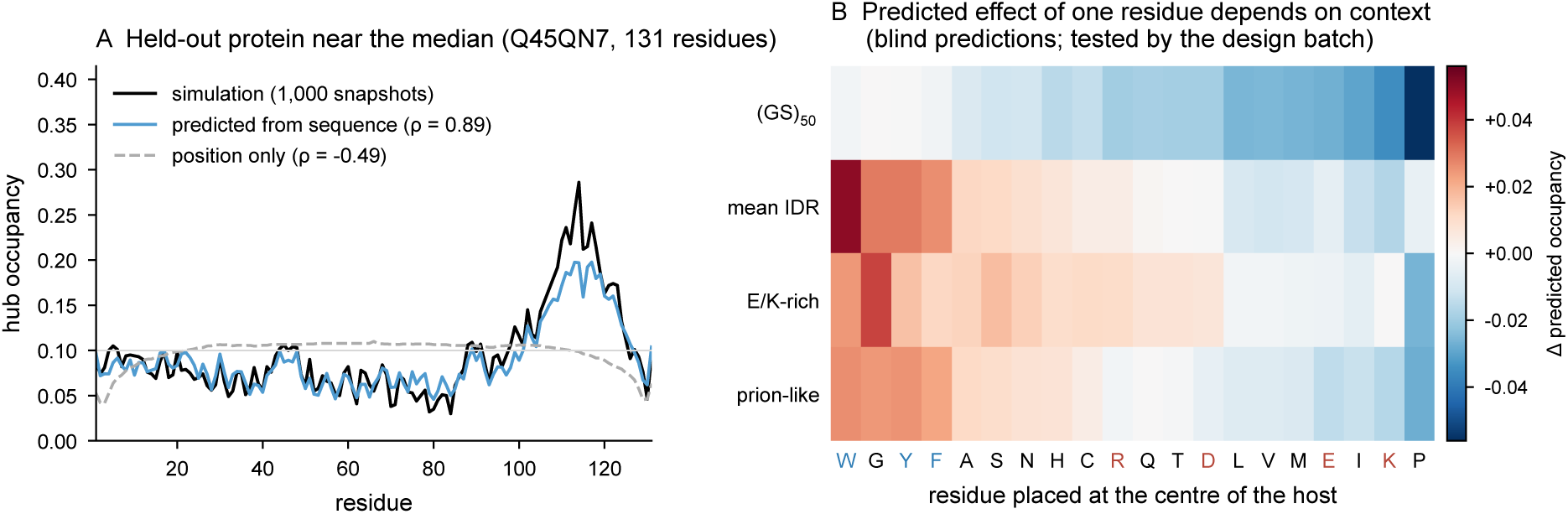
Details of the sequence-based predictor. (A) Simulated CALVADOS-2 and predicted occupancy for a held-out protein with a median score (Q45QN7, 131 residues), together with the prediction from chain position alone. (B) Predicted change in occupancy when the central residue of four 100-residue hosts is replaced, from consensus-model predictions that were recorded before the corresponding designed sequences were simulated.

**Figure S6:**
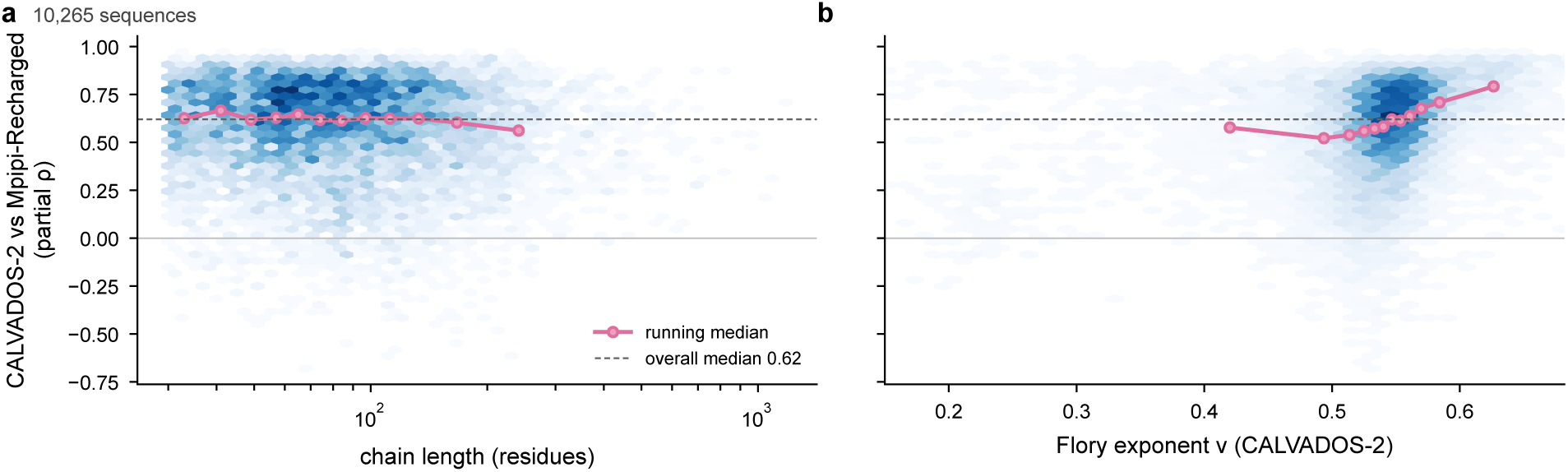
Agreement between the hub occupancy of CALVADOS-2 and Mpipi-Recharged for the same sequences, after removing chain position, against chain length (a) and the Flory exponent of the CALVADOS-2 ensemble (b), shown as density with running medians over twelve equal-count bins. This agreement bounds any predictor that does not see the force field.

The choice of scale matters little, since the scale of the other model family costs at most 0.04. Proline matters only in Mpipi-GG, whose parameters enlarge the excluded volume of proline by a third, strengthen the attraction between glycine residues and between glycine and serine 2.19-fold and weaken the attraction between charged and aromatic residues by 60% [8]. We note that the gap between the context model and the full model in Mpipi-GG is not a consequence of the smaller set of proteins simulated with Mpipi-GG. On the same 908 proteins, the CALVADOS-2 labels give 0.84 for the context model and 0.88 for the full model, and the CALVADOS-2 labels of 908 randomly drawn proteins give 0.84 and 0.87. In addition, context models trained on CALVADOS-2 predict the same held-out proteins in Mpipi-GG and Mpipi-Recharged with medians of 0.50 and 0.64, against 0.53 and 0.69 for the full model and 0.53 and 0.70 for the agreement between the simulations. The context model trained on CALVADOS-2 labels also predicts the 1,791 simulated shuffles with a median of 0.84, against 0.89 for the full model fixed in advance, although this analysis was made after the shuffle labels were known. Finally, when groups of residue-level features are permuted on the held-out folds, the net charge and the stickiness of the windows and the stickiness of the central residue cost the most accuracy, whereas the aromatic fraction of the windows adds at most 0.04 beyond the window stickiness.

To characterise this context, we called the 10% of residues of each protein with the highest position-adjusted occupancy the hubs of that protein and compared the composition of the ten residues around each hub, five on either side, with the composition around residues of the same type, so that a glycine hub is compared with other glycines (Table S8 and Supplementary Fig. S9). The comparison with residues of the same type matters because many hubs are glycines, 40% in CALVADOS-2 against 10.5% of all residues, and glycines cluster, so a comparison with all residues gives a larger glycine enrichment of 31 to 58%. Relative to the same type, glycine is enriched around hubs by 15 to 30% in the three force fields and by 63% in the ensembles built from experimental data, proline is depleted to 0.59 in Mpipi-GG and to 0.64 in the experimental ensembles but only to 0.93 and 0.95 in Mpipi-Recharged and CALVADOS-2, lysine is depleted in CALVADOS-2 and Mpipi-Recharged, and aromatic neighbours are enriched only in Mpipi-Recharged.

**Figure S7:**
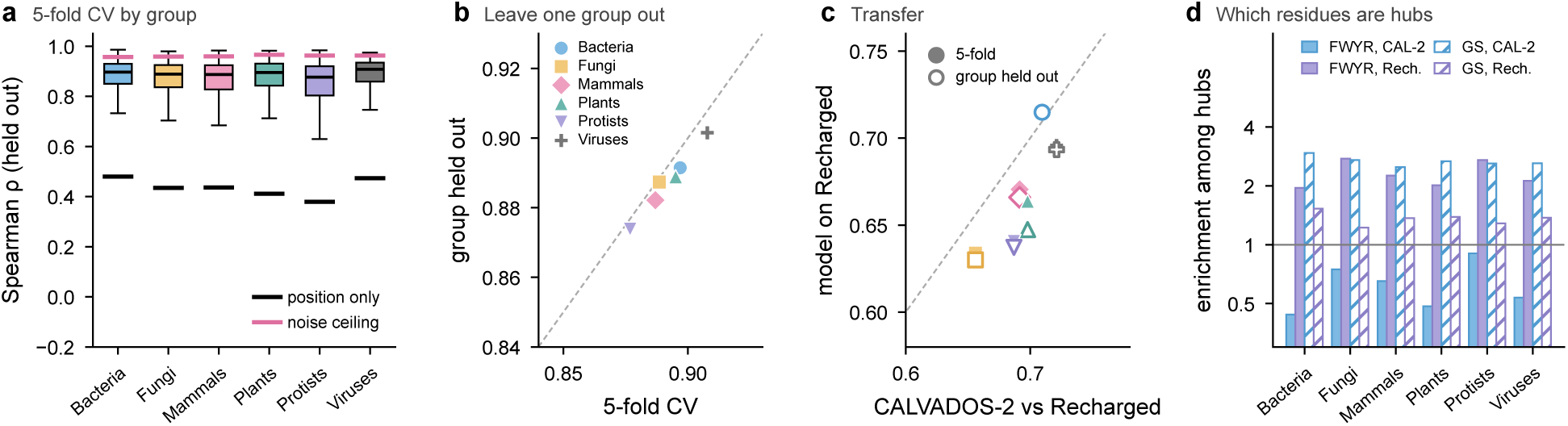
Hub occupancy across the six groups of the BENDER set. (a) Held-out per-protein Spearman correlation of the full model in five-fold cross-validation for each group, with the median of chain position alone (black) and the median noise ceiling (magenta). (b) Median correlation when each group is held out of training, against the five-fold value for the same proteins. (c) Median correlation of the CALVADOS-2 model with the Mpipi-Recharged labels, for the five-fold (filled) and the held-out-group (open) models, against the median agreement between the two simulations of the group. (d) Enrichment of the canonical stickers F, W, Y and R (filled) and of glycine and serine (hatched) among hubs, the 10% of residues of each protein with the highest hub occupancy, in CALVADOS-2 and Mpipi-Recharged.

To test whether the glycine enrichment arises from the small bead of glycine, we repeated the comparison on the 907 sequences simulated with all three force fields, with hubs defined by the position-adjusted *z*-score of time-averaged betweenness in the simulations, in the matched residue-neutral nulls and in the excess of the former over the latter (Table S9). The nulls, which contain the bead sizes of each force field but no attraction, enrich glycine around null hubs by at most 3%, whereas the excess retains a glycine enrichment of 7 to 16% and, in Mpipi-GG, a proline depletion to 0.67. These patterns therefore arise from attraction acting on the arrangement of residues rather than from geometry alone.

### S8 Ensembles built from experimental data

We used ensembles from the Protein Ensemble Database [31] that describe disordered proteins, which we required to have a mean relative solvent accessibility of at least 0.55 and a mean DSSP entropy of at least 0.15 in the database’s own analysis, and we excluded full-length proteins that contain folded domains. Table S10 lists the 18 ensembles used during development and the 11 blind ensembles, whose predictions were recorded before the labels were computed. Supplementary Fig. S10 shows how such ensembles are built and how hub occupancy is read from the deposited models, Supplementary Fig. S11 shows the label of one blind ensemble in detail, and Supplementary Fig. S12 shows the predictions for all 11 blind ensembles. The three blind ensembles marked with *a* were scored after the partial correlation had been fixed as a co-primary metric. Several entries describe the same protein with different restraints or methods, so the development set contains 14 distinct sequences from 13 proteins. We note that none of the deposited files contains per-model weights, so every model contributes equally to the hub occupancy of the ensemble. For the blind ensembles of A*β*40 and ACTR, which are molecular dynamics trajectories with the a99SB-disp force field, this means that the consecutive and non-redundant frames represent the unweighted simulation, because the maximum-entropy weights of these trajectories were not deposited. Moreover, the blind ensemble of CBP ID4 was generated with flexible-meccano from backbone propensities derived from chemical shifts, without selection against experimental data [45]. To test whether these ensembles affect the result, we left out the two unweighted simulations, after which the nine remaining blind ensembles give a median partial correlation of 0.24 for the model and 0.10 for the null. Excluding CBP ID4 as well leaves eight blind ensembles that were selected or refined against experimental data, for which the median partial correlation is 0.30 for the model and 0.13 for the null, with the model above the null in six of the eight.

**Figure S8:**
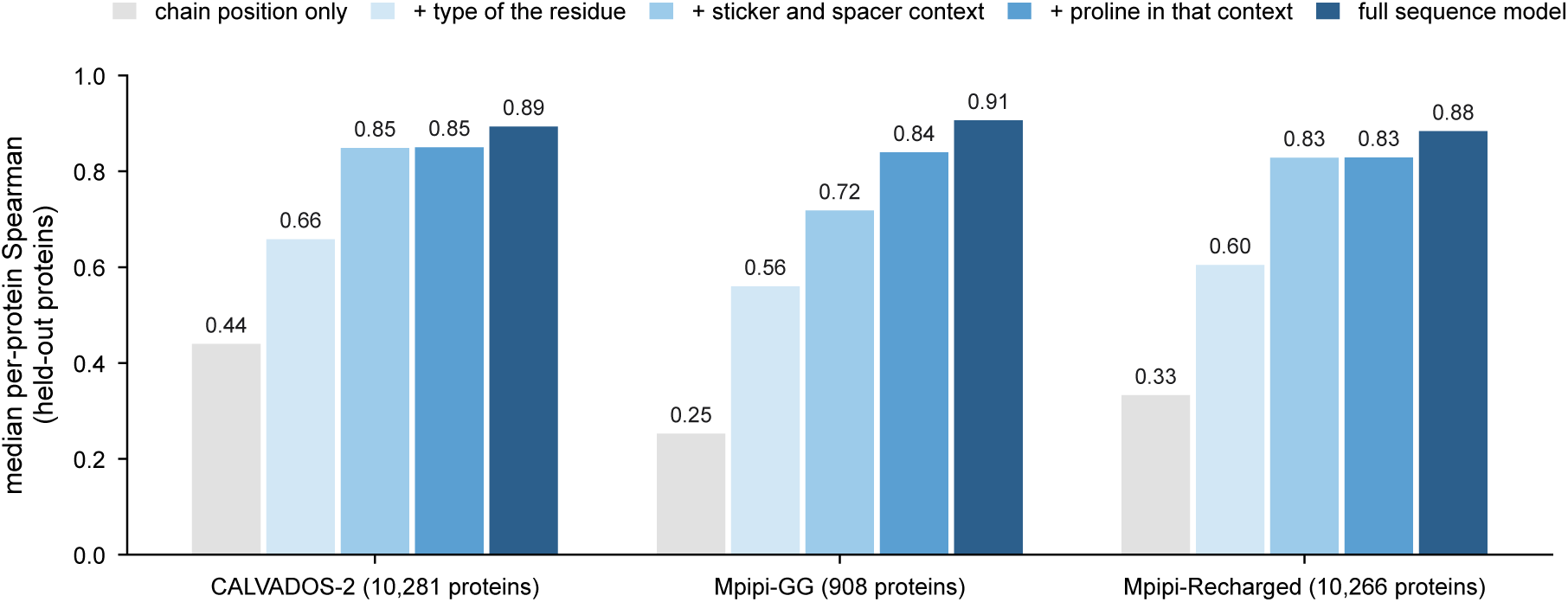
Median held-out per-protein Spearman correlation between predicted and simulated hub occupancy in each force field, for gradient-boosted trees that see chain position only, together with the type of each residue, together with the residue’s sticker and spacer surroundings, with the proline content of the surroundings added, and for the full model with 487 sequence features (Supplementary Table S7).

Individual ensembles also differ in how informative the labels are. For example, the blind ensemble of 4E-BP2 phosphorylated at five sites passes our disorder criteria, but six residues between positions 34 and 56 are hubs in 78 to 96% of the models, which is consistent with the four-stranded *β*-domain into which phosphorylation folds residues 18 to 62 of this protein [46], so the hubs of this ensemble change little from model to model. Similarly, PED00001 contains only 11 models and PED00023 only 16, so the labels of these two ensembles are noisier than the rest.

Table S11 lists the agreement in hub occupancy between independent ensembles of drkN and of Sic1, which sets the scale for the correlations in Table S10. The ensembles of each protein were built with different methods and combinations of restraints, and the median agreement between ensembles of the same protein is 0.54 for drkN and 0.26 for Sic1. Part of the disagreement between the Sic1 ensembles may be real, because two of these ensembles describe the phosphorylated protein. Chain position alone reaches 0.78 to 0.83 in three of the four ensembles of drkN, whereas the hubs of the five Sic1 ensembles are not set by chain position. Against the Sic1 ensembles, the model reaches a median of 0.39, compared with 0.25 for the null and 0.16 for chain position.

Finally, how much a restrained ensemble tests experiment, rather than the generator of the starting conformers, can be measured for the three proteins whose restraint-free pools are deposited. Hub occupancy in the restrained ensemble correlates with that of the corresponding pool with Spearman coefficients of 0.29 for *α*-synuclein, 0.51 for Sic1 and 0.69 for drkN, and the occupancy of an individual residue changes on average by 0.047, 0.032 and 0.037 between pool and restrained ensemble, compared with an average occupancy of 0.10.

**Figure S9:**
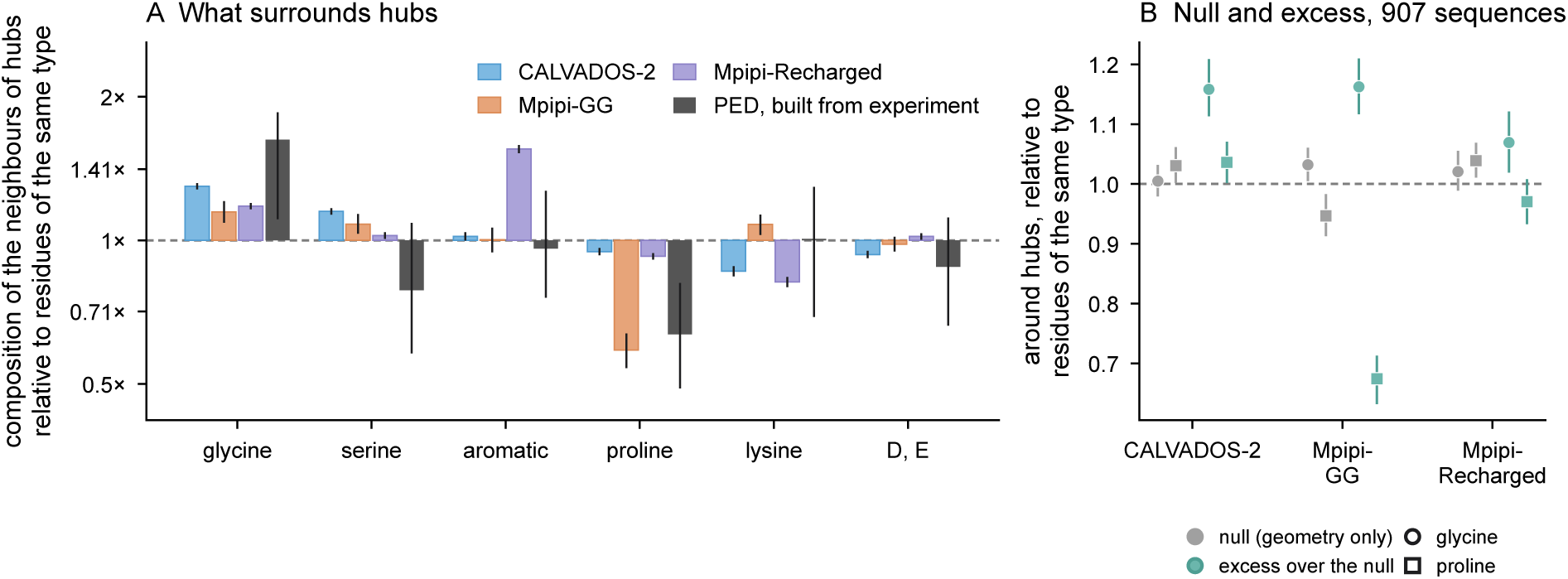
Composition of the ten residues around hubs relative to residues of the same type. (A) The three force fields and the 27 ensembles built from experimental data, with 95% bootstrap intervals over proteins (Supplementary Table S8). (B) Glycine (circles) and proline (squares) around hubs on the 907 sequences simulated with all three force fields, for the hubs of the residue-neutral null, which contains the bead sizes of each force field but no attraction, and for the excess of the simulations over the null (Supplementary Table S9).

### S9 A second blind set and all-atom force fields

After the analysis of Supplementary Section S8 was complete, we listed all 465 experimental entries of the Protein Ensemble Database and selected unused single-chain ensembles of disordered regions with at least about 200 models. All candidates pass the disorder criteria of Supplementary Section S8. In addition, we required a mean AIUPred disorder score [47] of at least 0.5 over the construct, which removes the ensemble of IAPP (0.43). No UniProt accession of the new set occurs among the simulated sequences, but in seven entries from five proteins a simulated ortholog shares exact 12-residue windows covering 52 to 67% of the construct, so we also report the nine proteins without such an ortholog. The same check finds no ortholog for any of the 29 ensembles of Table S10, where the largest coverage is 29%. All ensembles deposited for an entry were pooled and thinned to at most 1,000 models, and labels and the five-pool null were computed as before. Before any label was computed, the plan of this test, including these exclusions, and the predictions of the frozen consensus model were hashed with SHA-256.

Over the 14 proteins of the primary set (Table S12 and Supplementary Fig. S13), the median Spearman correlation is 0.39 for the model, 0.33 for the null and 0.38 for chain position, and the median partial correlation is 0.31 for the model and 0.03 for the null. The partial correlation of the model is positive in 13 of the 14 proteins (*P* = 0.0009, one-sided sign test) and above that of the null in 10 (*P* = 0.09), so the pre-registered criterion, which required *P <* 0.05 for the second comparison, is not met. In the sensitivity analysis, the nine proteins without an ortholog give 0.35 for the model and 0.04 for the null, with the model positive in eight and above the null in six. The weakest case is the linker of the SARS-CoV-2 nucleocapsid protein, whose hubs follow chain position closely (0.79). We note that pooling both blind sets, which was not planned in advance, places the model above the null in 19 of 25 proteins (*P* = 0.007). Finally, five further ensembles cover proteins of Table S10 in new conditions or as overlapping constructs, such as FUS NTD in the condensed state and phosphorylated Sic1, and are listed without a test.

The Protein Ensemble Database also holds ensembles of A*β*40, ACTR, drkN, PaaA2 and *α*-synuclein from explicit-solvent simulations with a99SB-disp in the a99SB-disp water model and with CHARMM22* and CHARMM36m in TIP3P water [48], of which the a99SB-disp ensembles of A*β*40 and ACTR belong to the first blind set. Each deposited ensemble holds 3,000 frames of the unbiased simulation without the maximum-entropy weights, so these ensembles compare three all-atom force fields rather than experiment. Hub occupancy agrees between these force fields with a median Spearman coefficient of 0.56 over 15 pairs and of 0.51 once the distance from the chain ends is removed (Table S13), close to the agreement between the coarse-grained force fields. Agreement is lowest for *α*-synuclein, at 0.18 to 0.24. In contrast, the all-atom ensembles agree less well with the expert-built ensembles of the same protein, with a median partial correlation of 0.22 over 15 pairs. In addition, the consensus model, trained on coarse-grained simulations, predicts the CHARMM22* and CHARMM36m ensembles of ACTR and drkN poorly, with partial correlations between −0.30 and −0.12.

**Figure S10:**
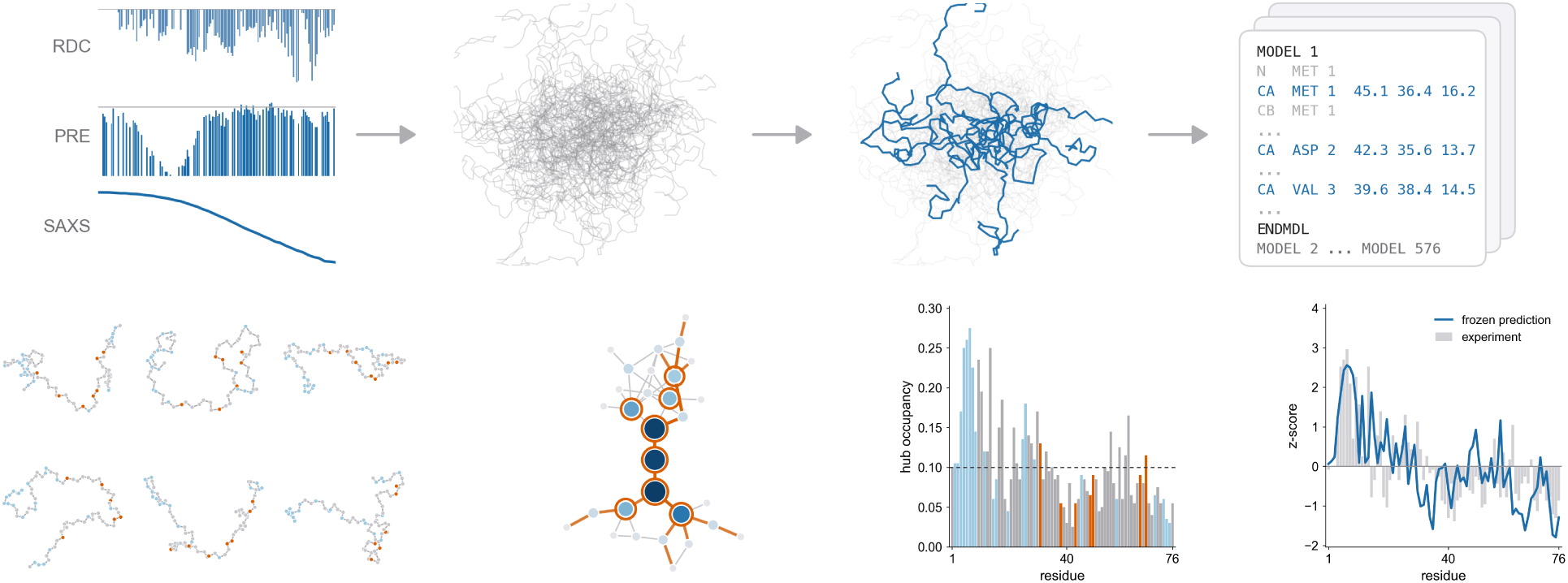
How ensembles are built from experimental data and how hub occupancy is read from such ensembles. Top, from left to right, measurements for *α*-synuclein (backbone residual dipolar couplings, PRE intensity ratios for the spin label at residue 42 and the SAXS profile), a pool of conformers drawn for illustration, a subset selected from the pool, and an excerpt of the deposited file of PED00024 with the C*_α_* lines highlighted. Bottom, from left to right, six of the 200 deposited models of the N-terminal region of MKK4 (a blind ensemble), the largest contact cluster of one model drawn as a network with the eight hubs of that model ringed, the hub occupancy of every residue across the 200 models, highest in the N-terminal stretch rich in glycine and serine, and the standardised occupancy (grey bars) with the prediction recorded in advance (blue line), which correlate at 0.38.

### S10 Reweighting simulations against paramagnetic relaxation enhancement data

We also tested whether simulations reweighted against paramagnetic relaxation enhancement (PRE) data could serve as additional experimental references. Specifically, PRE rates were predicted from C*_α_* distances with a single correlation time per protein, intensity ratios were converted to rates with standard relaxation and delay parameters, and frame weights were optimised by Bayesian maximum-entropy reweighting with the strength of the restraint chosen by leave-one-label-out cross-validation, keeping the unweighted ensemble whenever reweighting did not improve the held-out labels. We applied this procedure to the restraint-free pools of *α*-synuclein and drkN with PRE data from the PeptoneDB-Integrative set [49]. The values provided for Sic1 include the diamagnetic relaxation rate, which stays near 30 s*^−^*^1^ even far from every spin label, so we did not use the Sic1 values. With the pre-declared exclusion of residues within three positions of each label, close approaches between nearby C*_α_* atoms dominate the predicted rates, and reweighting concentrates the weight on 2% of the frames without moving hub occupancy towards that of the expert-built ensembles (Table S14). When residues within nine positions of each label are excluded instead, the forward model reproduces the PRE data of the expert-built *α*-synuclein ensemble well and those of the unweighted pool almost as well, so that reweighting is no longer needed. However, the hub occupancy of these two ensembles, which are both consistent with the same PRE data, agrees only with a Spearman coefficient of 0.29. Taken together, a handful of spin labels does not determine the residue-level contact network, and we therefore do not use reweighted simulations as references.

As a model-free check, we correlated the model’s predictions with the excess of each PRE profile above the trend of that profile with sequence separation. This check gives correlations between −0.43 and 0.43 with a median of 0.03 over seven proteins (Supplementary Fig. S14), so raw PRE profiles do not resolve the positions of hubs either.

**Figure S11:**
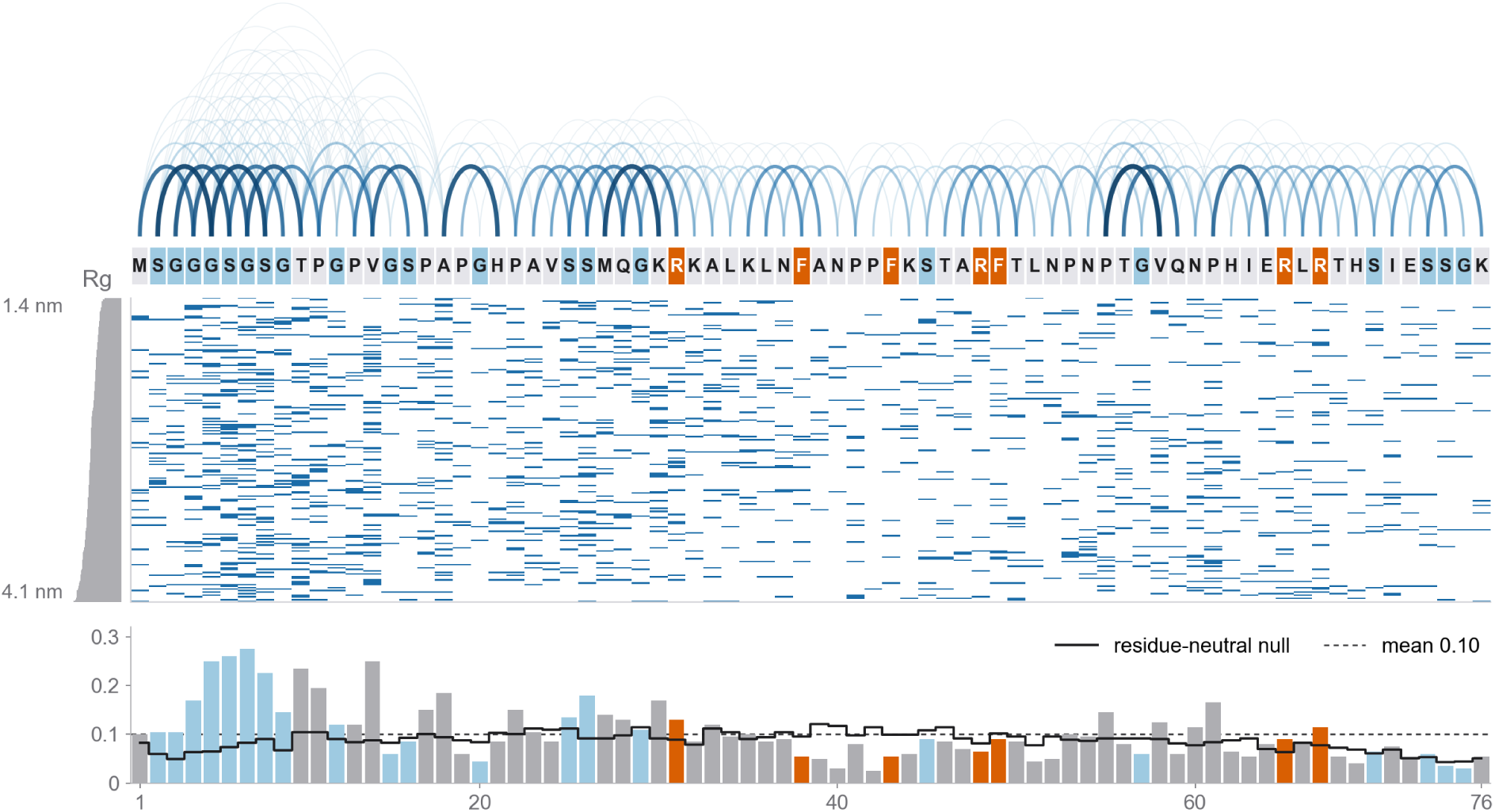
The hub label of the MKK4 ensemble in detail. From top to bottom, the contacts of the 200 models drawn as arcs over the 76-residue sequence, densest in the N-terminal stretch rich in glycine and serine, the sequence with the canonical stickers in orange and glycine and serine in light blue, the eight hubs of every model with one row per model sorted by radius of gyration from 1.4 to 4.1 nm, which differ from model to model, and the hub occupancy as the mean over the models, highest at G5 to S8 near 0.25 and well above the residue-neutral null of 0.06 to 0.09.

### S11 Missense variants in human disordered regions

To ask whether predicted hubs relate to disease variants, we followed the ClinVar analysis of IDRome [6]. We used the 18,445 ClinVar [50] missense variants that IDRome mapped to the human IDRs of the database, of which 17,214 have a wild-type residue that matches the IDR sequence (2,706 pathogenic or likely pathogenic and 14,508 benign or likely benign). As a replication set, we used the 54,542 missense variants of the ClinVar release of 5 October 2026 that map to the same IDRs and are not part of the first set (7,725 pathogenic or likely pathogenic). The frozen consensus model predicted the hub occupancy of every IDR and of every variant, and we compared pathogenic with benign variants by two-sided Brunner–Munzel tests as in IDRome, reporting Cohen’s *d* as pathogenic minus benign. Before these comparisons, we had pre-registered a test of whether pathogenic variants are enriched at predicted hubs, which instead found a depletion (odds ratio per quarter of the within-IDR percentile of predicted occupancy 0.85 and 0.83 in the two sets).

Collagens dominate these comparisons, since a quarter of the pathogenic variants of the first set (717 of 2,706) and almost half of those of the replication set (3,396 of 7,725) lie in collagens, and 94 and 97% of these replace a glycine of the Gly-X-Y repeats, which form folded triple helices in the cell but count as disordered in simulations of isolated chains. The model places hubs on these glycines, so collagen variants remove hubs and reverse the direction of both comparisons when included (Table S15 and Supplementary Fig. S15). Outside collagens, pathogenic variants sit at residues of lower predicted hub occupancy than benign variants, with *d* = −0.27 and −0.35 in the two sets, and raise the occupancy of the substituted residue more often, with *d* = +0.21 and +0.16, at *P <* 10*^−^*^14^ in every case. Removing the NOTCH3 cysteine variants, as in IDRome, changes neither result. These effect sizes are similar to the difference in conformational entropy per residue that IDRome reported between pathogenic and benign variants (*d* = −0.27).

Like those of IDRome, these unadjusted differences partly reflect sequence composition and conservation.

**Figure S12:**
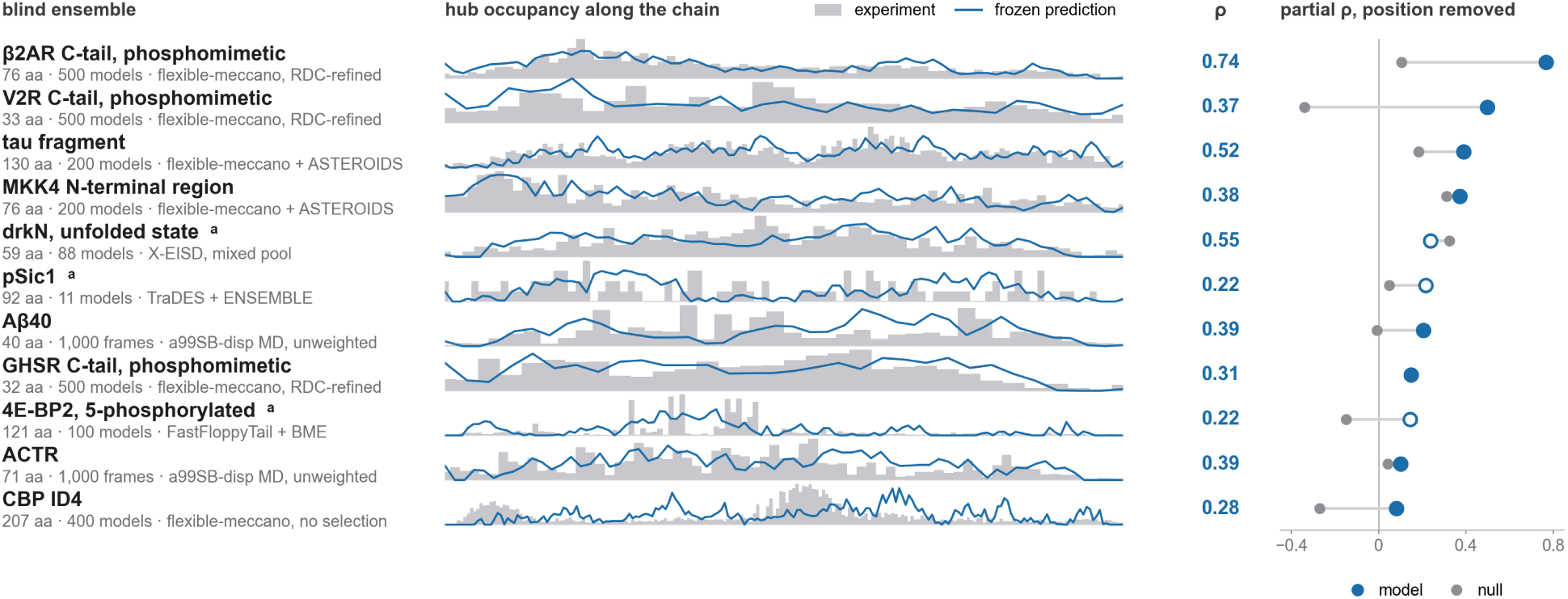
Predictions recorded in advance for the 11 blind ensembles, sorted by partial correlation. Each row gives the ensemble with length, number of models and method, the hub occupancy along the chain (grey) with the prediction (blue), the Spearman correlation, and the partial correlation after removing chain position for the model (blue) and the residue-neutral null (grey). The three ensembles marked *a*, drawn with open symbols, were scored after the partial correlation had been fixed as a co-primary metric.

In logistic regressions that also account for the wild-type and mutant residues, the distance from the IDR ends, the IDR length, the glycine and serine content around the variant and the severity of the substitution, with standard errors clustered by gene, the lower occupancy at pathogenic sites outside collagens is no longer significant. The gain of occupancy, in contrast, remains associated with pathogenicity, with odds ratios per standard deviation of 1.20 and 1.15 (*P* = 0.01 and 0.001). Adding the conservation of the residue among IDR orthologues, on which ClinVar classifications partly draw, reduces these odds ratios to 1.11 and 1.08 (*P* = 0.19 and 0.09). All comparisons other than the pre-registered test were made after the result of that test was known and are exploratory. Taken together, predicted hubs mark glycine- and serine-rich stretches in which variants are mostly tolerated, and hub occupancy describes the single-chain contact network rather than functional constraint.

### S12 Design batch

The prospective batch contains 292 sequences, to be simulated with all three force fields under the protocol of Table S1, and the 113 sequences of the experimental constructs, the PRE proteins, the replicas and the variants have so far been simulated with CALVADOS-2 and Mpipi-Recharged. The batch comprises the 22 distinct experimental constructs behind the ensembles of Table S10, with phosphorylated Sic1 represented by a phosphomimetic sequence, four further disordered proteins with PRE data and 75 variants of the experimental constructs, in which aromatic residues were replaced by serine, arginine and lysine were exchanged for each other, acidic residues were neutralised or the sequence was shuffled. The remaining sequences are 20 homopolymers of 100 residues, every amino acid placed at the centre of four 100-residue hosts together with substitutions near the ends of a glycine–serine host, valence and spacing series of aromatic and charged residues, a series of eight glutamate and lysine sequences that span the range of charge patterning [11], and 12 BENDER sequences that are simulated a second time to measure the noise between independent runs. Each host and guest sequence was simulated three times in Mpipi-Recharged, because the change in occupancy that a single substitution produces at one residue is comparable to the noise of a single run.

The predictions of the frozen consensus model were recorded and hashed on 2 October 2026, before any of these sequences was simulated, and the analysis of the responses to substitutions was pre-registered before any label existed. The 12 replicas agree with the runs they repeat at a median Spearman correlation of 0.93 in CALVADOS-2 and 0.95 in Mpipi-Recharged, and the between-run noise of a residue, which grows with occupancy, sets the reliability of each simulated response. After removing chain position, the predictions correlate with the simulated occupancy at a median of 0.79 and 0.74 for the 22 constructs, 0.68 and 0.67 for the four PRE proteins and 0.74 and 0.72 for the 75 variants, in CALVADOS-2 and Mpipi-Recharged respectively, and positive for every sequence except one construct in CALVADOS-2. For the 49 substitution variants, the response passes the pre-registered criterion in both force fields (Fig. 4d and Supplementary Fig. S16a,b), with medians of 0.34 and 0.49, positive in 44 and 46 variants (*P* = 4 × 10*^−^*^9^ and 4 × 10*^−^*^11^) and at 63% and 56% of the noise ceiling. The response is predicted best for neutralised acidic residues, with 0.57 and 0.54, and least well for aromatic residues replaced by serine, with 0.14 and 0.35. The rule based on local stickiness and charge reaches 0.24 and 0.43, so the model is better in 36 of the 49 variants in CALVADOS-2 (*P* = 0.0005, paired Wilcoxon test) and in 29 in Mpipi-Recharged (*P* = 0.08). The predicted and simulated shift of the occupancy-weighted centre of the chain correlate at 0.68 and 0.76 over variants. At the substituted residues, predicted changes are 1.3 and 3.6 times smaller than simulated changes, and beyond ten residues 3.3 and 5 times smaller, so the model orders the responses correctly but underestimates the size of distant responses, which exceed the change expected from between-run noise by factors of 1.3 and 2.0. Averaged over the substituted residues, the predicted and simulated changes correlate at 0.72 and 0.88 across variants and agree in sign for 78% and 92% of variants (Fig. 4e). Against the mean of the two force fields, the response is predicted at a median of 0.54 and the change at the substituted residues at 0.90, higher than against either force field alone. The 26 shuffled variants are predicted at medians of 0.76 and 0.73.

**Figure S13:**
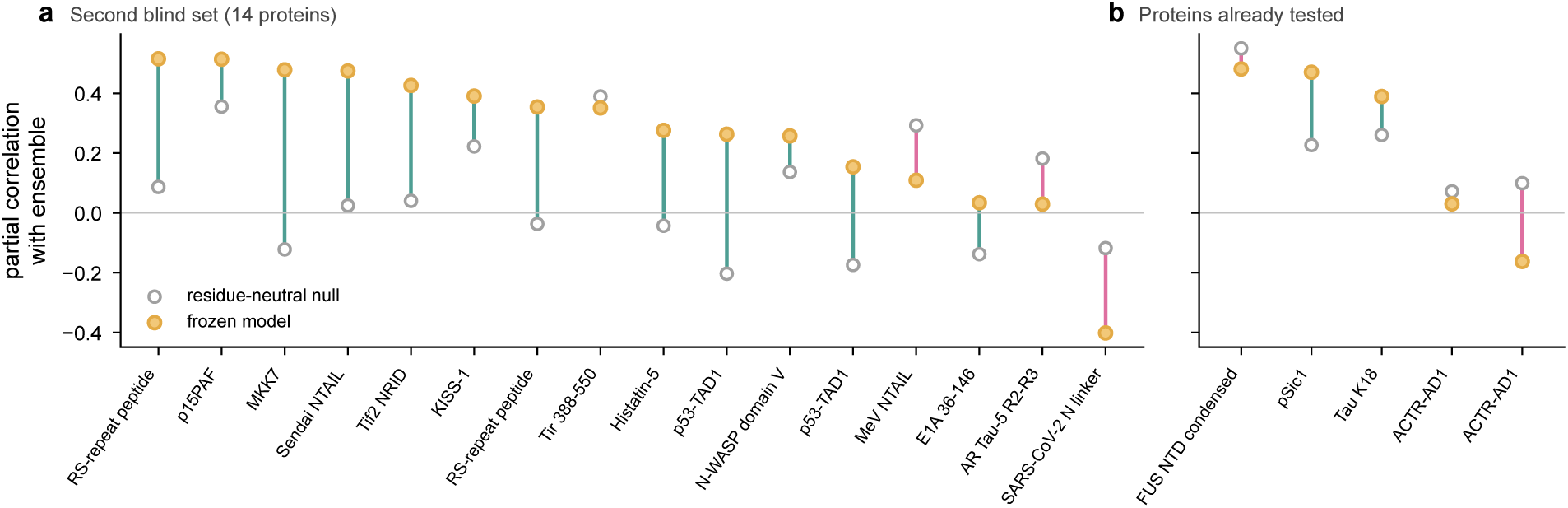
The second blind set. Partial correlation between the frozen predictions and the hub occupancy of each ensemble after removing chain position, for the model (orange) and the residue-neutral null (open), with teal lines where the model exceeds the null and magenta lines where the null exceeds the model, for the 14 proteins of the primary set (a) and the five ensembles of proteins already tested, under new conditions or as overlapping constructs (b).

**Figure S14:**
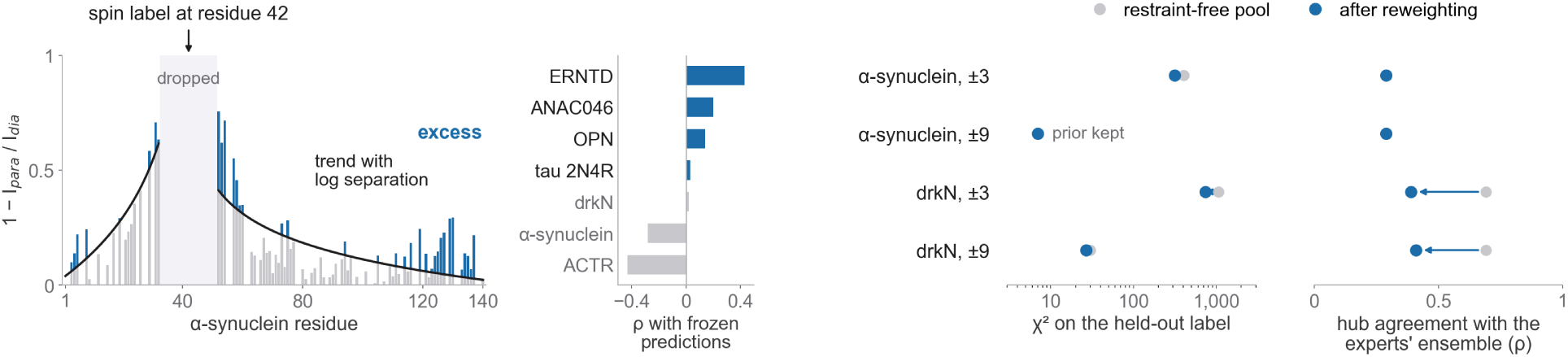
Raw PRE profiles and reweighting against PRE data. Left, the PRE profile of *α*-synuclein for the spin label at residue 42 as one minus the intensity ratio, with residues within ten positions of the label excluded, the trend with sequence separation on either side (black) and the excess above the trend (blue), together with the Spearman correlation between such excess profiles and the model’s predictions for seven proteins, the four not used during development in blue. Right, reweighting of the restraint-free pools of *α*-synuclein and drkN against PRE data with residues within three or nine positions of each label excluded, with the held-out *χ*^2^ and the agreement of hub occupancy with the expert-built ensembles before and after reweighting (Table S14).

**Figure S15:**
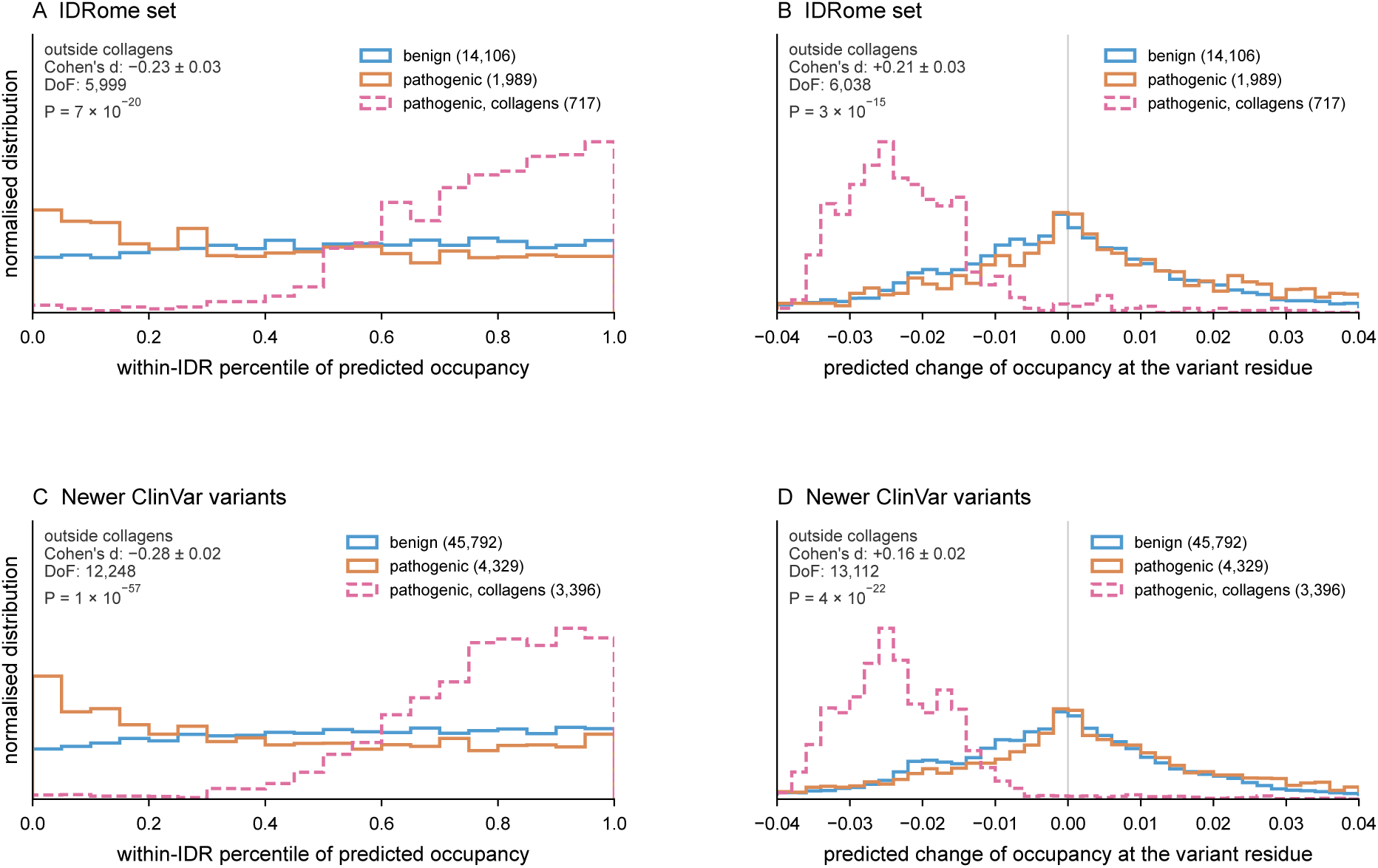
Predicted hub properties of pathogenic and benign ClinVar missense variants in human IDRs, shown as in IDRome. Normalised distributions of the within-IDR percentile of the predicted occupancy of the wild-type residue (A, C) and of the predicted change of occupancy at the variant residue (B, D), for the variants mapped by IDRome (A, B) and for newer variants of the ClinVar release of 5 October 2026 (C, D). Solid lines show benign and pathogenic variants outside collagens, with Cohen’s *d*, Brunner–Munzel degrees of freedom and two-sided *P* of the comparison between the two, and dashed lines show pathogenic variants in collagens, almost all glycine substitutions in Gly-X-Y repeats. The window of B and D, from −0.04 to 0.04, contains 96% of all variants.

The constructs of the batch also allow a comparison of simulation, prediction and experiment for the same sequences (Table S16 and Supplementary Fig. S16c). For the 20 constructs whose PED ensembles were built from experimental data, the hub occupancy simulated with CALVADOS-2 agrees with that of the experimental ensembles at a median Spearman correlation of 0.47, or 0.38 after removing chain position, and the hub occupancy simulated with Mpipi-Recharged at 0.34, or 0.33. Both simulations exceed the residue-neutral null after removing chain position in 18 of the 20 constructs. The predictions of the frozen consensus model agree with the experimental ensembles at 0.43, or 0.37, as closely as the better force field, and exceed the CALVADOS-2 simulation after removing chain position in 12 of the 20 constructs and the Mpipi-Recharged simulation in 14. Such agreement is expected if the model, trained on the occupancy averaged over force fields, captures the arrangement part that the force fields share, whereas a single simulation also carries the residue-type part of the force field used. This comparison was not declared in advance, 14 of the constructs belong to the development set, and independent experimental ensembles of the same protein agree at only 0.31 (Supplementary Table S11), so differences of a few hundredths between the columns of Table S16 are not meaningful.

An exploratory analysis, carried out after these values were known, relates the agreement to residual secondary structure in the experimental ensembles. We assigned secondary structure to every deposited model with DSSP [51], computed from the backbone atoms, and found helix in at least 30% of the models for at least a tenth of the residues in five constructs, namely CBP ID4, PaaA2, drkN, the eIF4G1 fragment and p27 KID. For these five constructs, the predictions agree with the experimental ensembles at a median of 0.21 after removing chain position, against 0.39 for the other 15, and both simulations show a drop, from 0.46 to −0.09 with CALVADOS-2 and from 0.34 to 0.18 with Mpipi-Recharged. Across the 20 constructs, the agreement of the CALVADOS-2 simulation decreases with the mean helix fraction (Spearman correlation −0.48, *P* = 0.03), that of the predictions decreases less and not significantly (−0.37, *P* = 0.11), and that of the Mpipi-Recharged simulation shows no trend (−0.02), so whether the model or a simulation agrees better in helical proteins depends on the force field. Helical residues also rank higher in hub occupancy in the experimental ensembles than in the predictions, with a median Spearman correlation of 0.17 between the rank error and the helix fraction. Such a difference is expected from a contact definition that counts the contacts between residues *i* and *i* + 3 or *i* + 4 within a helix, which coarse-grained models without backbone hydrogen bonds do not form. Hub positions in disordered proteins with residual helices therefore reflect local structure in addition to residue arrangement, and the model is expected to apply mainly to chains without stable secondary structure.

**Figure S16:**
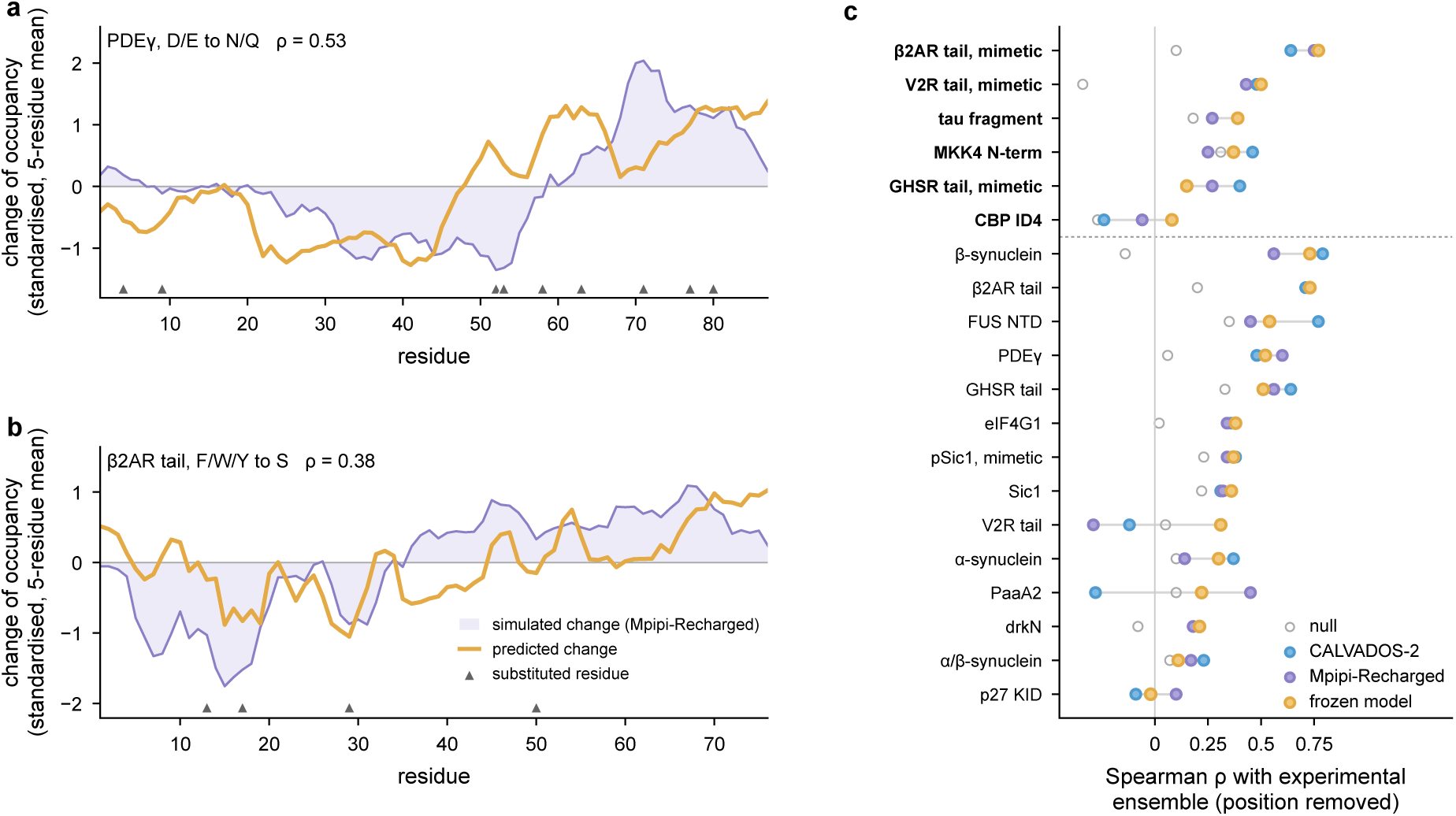
Prospective test of the frozen consensus model on the design batch simulated with CALVADOS-2 and Mpipi-Recharged, complementing Fig. 4c–e. (a, b) Change of hub occupancy along the chain simulated with Mpipi-Recharged and predicted by the model for a PDE*γ* variant with neutralised acidic residues (a) and a *β*_2_AR C-tail variant with aromatic residues replaced by serine (b), each profile standardised and shown as a running mean over five residues for display, with triangles marking the substituted residues. The two variants lie near the median correlation of the variant type. (c) Spearman correlation, after removing chain position, between the hub occupancy of the experimental PED ensembles and that of the two simulations, the frozen model and the residue-neutral null for the 20 constructs with ensembles built from experimental data, with the constructs not used during development in bold above the dashed line.

### S13 A case study of the FUS N-terminal domain

Supplementary Fig. S17 follows hub occupancy through the DEER-restrained ensemble of the 270-residue N-terminal low-complexity domain of FUS (PED00494), an ensemble of the development set with 551 models, and Supplementary Fig. S18 adds the view across the ensemble. In the model of median radius of gyration, the 27 residues with the highest betweenness hold together a network of 274 contacts. Between the tyrosine residues Y211 and Y235, the hubs G220 and G223 lie in an RGG segment and G226 and G231 on a run of ten glycine residues, where hubs sit at junctions of contact triangles. Neighbouring glycine residues such as G227, G228 and G230 are not hubs in this model but rank among the ten residues with the highest hub occupancy over all models, so hubs move from model to model while returning to the same glycine-rich stretches (Supplementary Fig. S18c). Over the whole chain, the CALVADOS-2 simulation of the same construct reproduces the experimental hub occupancy with a Spearman correlation of 0.76, compared with 0.45 for Mpipi-Recharged and 0.54 for the frozen consensus model (Table S16). The model places the main peaks on the glycine-rich stretches near residues 175 and 225, but underestimates the height of the peak near residue 225 and, like both simulations, misses part of the occupancy near residue 250. This arrangement of hubs on glycine runs between tyrosine residues is the pattern described for prion-like domains in Supplementary Section S15.

**Figure S17:**
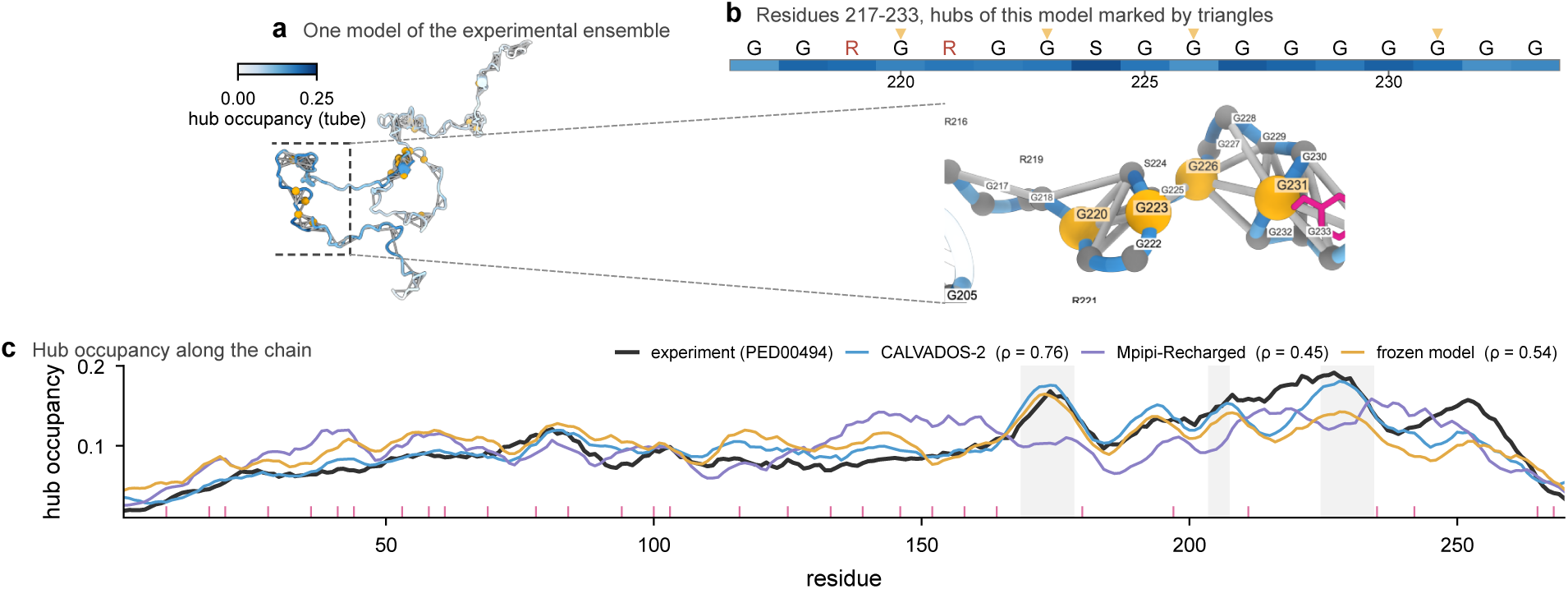
Hubs move between models of an experimental ensemble but return to glycine runs, shown for the FUS N-terminal domain (PED00494, 270 residues, 551 models). (a) The model of median radius of gyration coloured by hub occupancy, with contacts as light grey lines and the 27 hubs of this model as orange spheres. (b) A labelled close-up of residues 217 to 233, with the sequence above over a track of hub occupancy and triangles marking the hubs of this model. (c) Hub occupancy along the chain in experiment, in simulations of the same construct and in the frozen model (seven-residue means), with aromatic residues as ticks and glycine runs shaded.

### S14 Predicted hubs in designed sequences and human IDRs

To map these rules beyond the simulated sequences, we applied the frozen consensus model to designed sticker and spacer sequences and to all 28,058 IDRs of the human IDRome [6] in an exploratory analysis fixed in advance (Supplementary Fig. S19). In designed sequences, spacer composition matters more than sticker spacing, as glycine spacers carry the most hub occupancy beyond chain position and spacers with proline or charged residues the least, and aromatic stickers are hubs themselves, whereas arginine and lysine stickers pass hub occupancy to the neighbouring spacer residues. Across the human IDRome the same rules hold, as tryptophan, tyrosine, glycine and phenylalanine are enriched among predicted hubs and predicted occupancy rises with nearby glycine and aromatic residues, most visibly in prion-like domains.

The analysis of Supplementary Fig. S19 was specified before any prediction was computed and is exploratory and descriptive. Designed sequences of about 120 residues repeat a spacer of length *L* followed by one sticker, and residues within 15 positions of either end are left out of every summary. Excess occupancy is the prediction of the frozen consensus model minus a positional baseline, gradient-boosted trees on chain length and position trained on the CALVADOS-2 labels, so that chain-end effects do not dominate. For the human IDRome, predicted hubs are the 10% of residues of each IDR with the highest prediction. Across the 3.95 million residues of the 28,058 IDRs, predicted hub occupancy beyond the positional baseline rises from −0.010 for residues without glycine or aromatic residues within four positions to +0.083 for residues with seven glycine residues nearby, and each additional aromatic residue adds about 0.01. Supplementary Fig. S20 shows the predicted profiles of six prion-like domains, in which predicted hubs concentrate on glycine-rich stretches beside aromatic residues.

**Figure S18:**
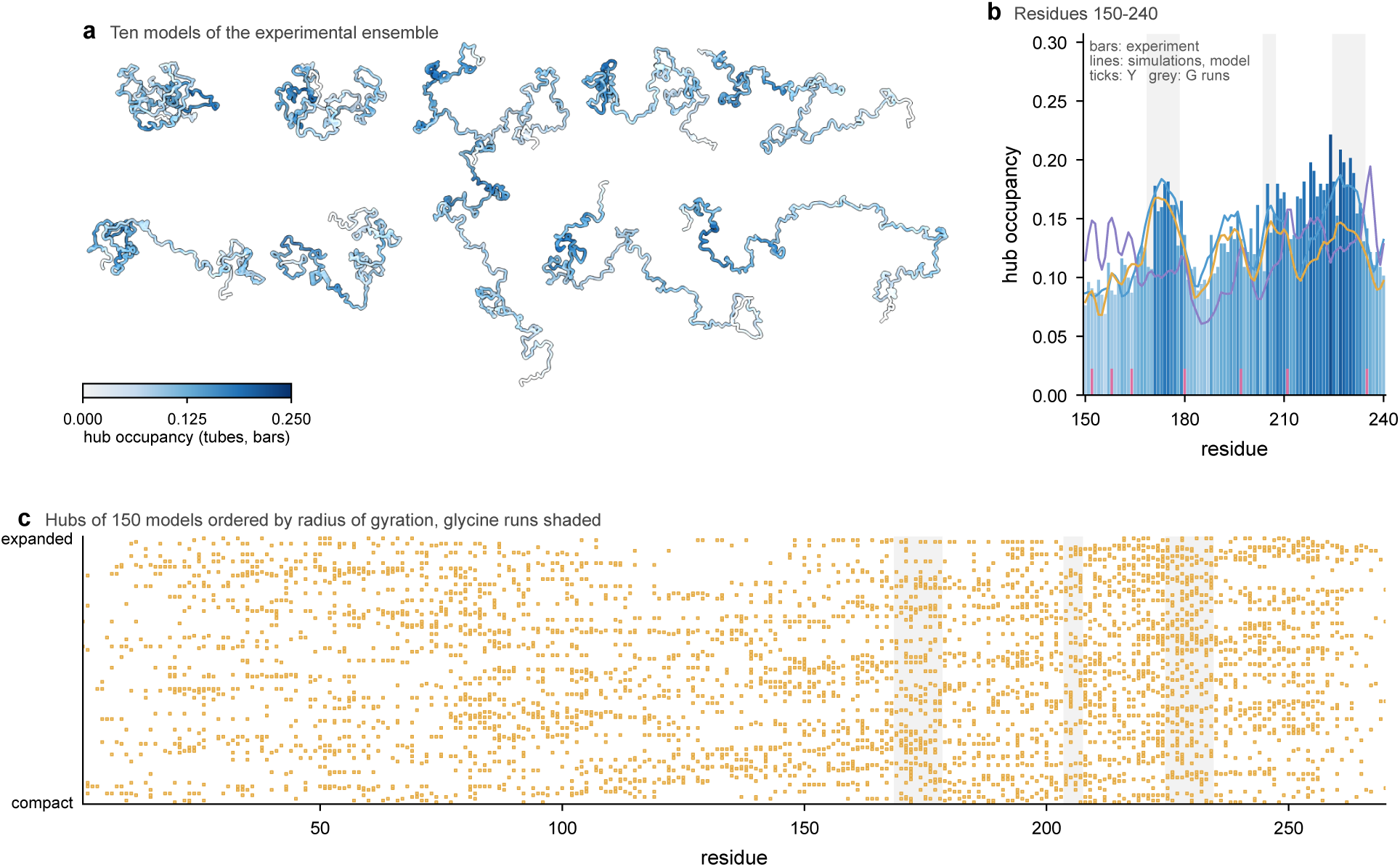
Hub occupancy across the experimental ensemble of the FUS N-terminal domain (PED00494, 270 residues, 551 models), complementing Supplementary Fig. S17. (a) Ten models spanning the range of radius of gyration, coloured by hub occupancy. (b) Hub occupancy of residues 150 to 240 in the experimental ensemble (bars), in the CALVADOS-2 and Mpipi-Recharged simulations of the same construct and in the frozen consensus model (lines), with tyrosine residues as ticks and runs of at least four glycine residues shaded. (c) Hubs of 150 of the 551 models ordered by radius of gyration from 2.7 to 8.7 nm, with glycine runs shaded.

**Figure S19:**
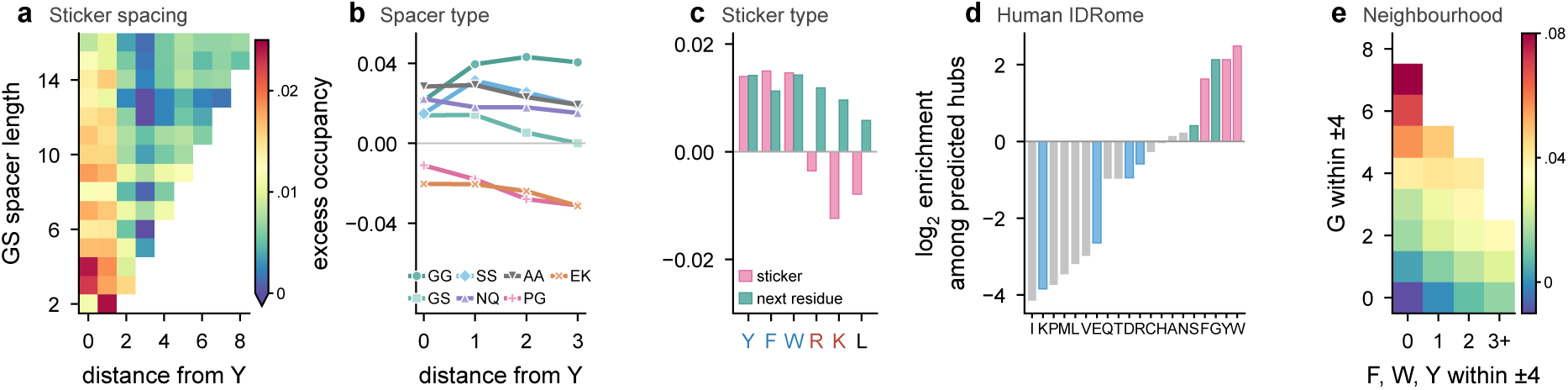
The learned map places hubs on glycine-rich spacers and aromatic residues, in designed sequences and across the human IDRome. All panels show predictions of the frozen consensus model as hub occupancy beyond the positional baseline (excess). (a) Excess against the distance from the nearest tyrosine for glycine and serine spacers of 2 to 16 residues. (b) The same for six-residue spacers of different composition. (c) Excess at the sticker and the next spacer residue for six-residue spacers between different stickers. (d) Log_2_ enrichment of each residue type among the predicted hubs of 28,058 human IDRs. (e) Mean excess of all residues of these IDRs by the number of glycine and aromatic residues within four positions.

**Figure S20:**
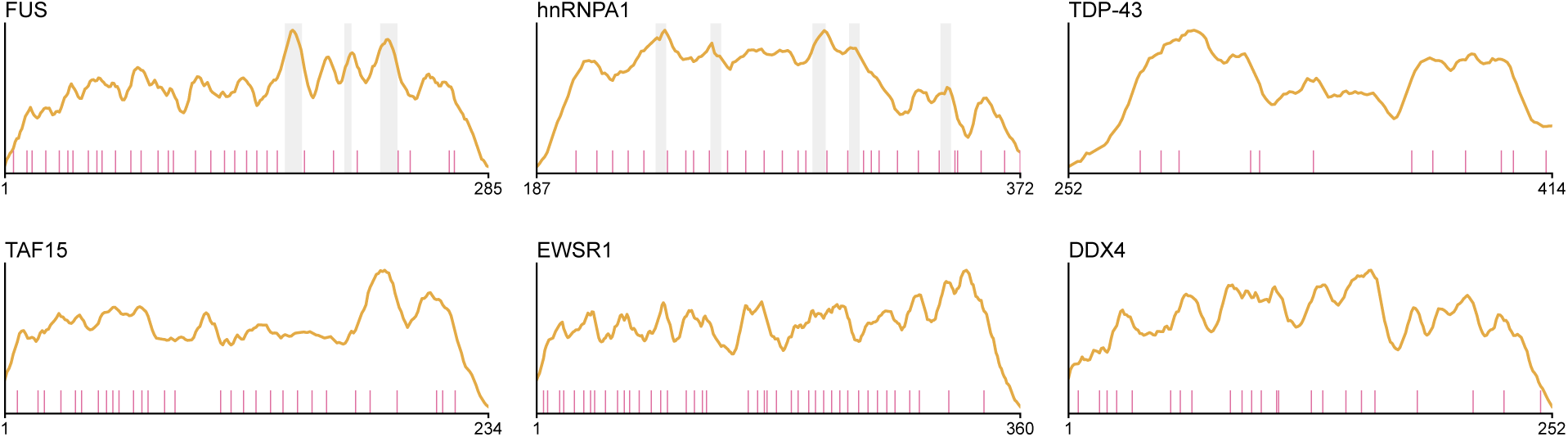
Predicted hub occupancy (seven-residue means) of the prion-like domains of FUS, hnRNPA1, TDP-43, TAF15, EWSR1 and DDX4 as defined in the IDRome, with aromatic residues as ticks and runs of at least four glycine residues shaded. Residue numbers refer to the full-length protein.

### S15 Predicted hubs and phase separation

We asked in four analyses whether predicted hub occupancy relates to phase separation, using only the frozen consensus model and no simulation of condensates (Table S17 and Supplementary Fig. S21). In an exploratory scan of 22 well-studied disordered regions that drive phase separation, the predicted hub profile of the wild type was more uneven than the profiles of sequences with residues permuted within windows of 25 residues, at a median percentile of 98 against 68 for 300 random regions of the human IDRome [6]. In the prion-like domains of hnRNPA1, hnRNPA2, FUS, DDX4, TAF15 and EWSR1, aromatic residues are more evenly spaced than in shuffled sequences, as reported before [2], whereas glycine residues are more clustered. The predicted hubs of these domains fall on glycine-rich stretches next to the aromatic residues rather than on the aromatic residues themselves, and across shuffled sequences the clustering of glycine is the composition feature most closely associated with uneven hub profiles, with Spearman correlations of 0.19 to 0.51.

Three pre-registered tests did not support a link between predicted hubs and phase behaviour. For 22 variants of the hnRNPA1 low-complexity domain with saturation concentrations measured at 4 °C [26], the share of predicted hub occupancy carried by aromatic residues and arginine does not explain the saturation concentration beyond a baseline of sticker number and net charge. For 30 human drivers of phase separation from PhaSePro [52], excluding the proteins of the exploratory scan, the unevenness of the wild type relative to local shuffles does not exceed that of five disordered regions matched in length and composition. For 23 disease mutations listed in PhaSePro as altering condensates, the predicted local change is not larger than that of the same substitution at other positions with the same wild-type residue. Repeated with predicted degree in place of predicted hub occupancy, under a plan hashed before any degree prediction and with Holm correction across the three, all three tests remain null (*P* = 0.99, 0.59 and 0.87). A comparison of the drivers of the exploratory scan with matched regions, made after these tests, still favours the drivers, which suggests that concentrated hubs may characterise prion-like and RGG-rich domains specifically, a hypothesis that requires an independent test.

### S16 Contact degree beside hub occupancy

The degree of a residue is the mean number of contacts per snapshot under the contact rule of the main text. In a comparison of the two readouts whose hypotheses and decision rules were hashed before the comparison was run, hub occupancy and degree correlate within proteins at 0.935, 0.953 and 0.964 in CALVADOS-2, Mpipi-Recharged and Mpipi-GG once chain position is removed. The part of hub occupancy that a monotone function of degree does not explain amounts to 7.3% of the variance, replicates between force fields (linear residual agreement 0.264 against 0.042 for a control) and belongs to bridging residues, which are enriched in tyrosine and phenylalanine, and in Mpipi-Recharged also in histidine and tryptophan, and depleted in lysine and in the aliphatic residues V, M, I and L. Residues of high degree but low occupancy are rich in glycine, which is 4.0 and 3.4 times more frequent among these residues in CALVADOS-2 and Mpipi-Recharged. Across 16 pairs of independent experimental ensembles of drkN and Sic1, degree agrees at a median of 0.426 and hub occupancy at 0.310, and degree is the more reproducible readout in 13 of the 16 pairs. For the 49 designed substitutions of Section 3.6, in a comparison that was not blind, the change in degree predicted by a CALVADOS-2 degree model correlates with the simulated change at a median of 0.48 and is positive in all 49 variants, against 0.35 and 44 of 49 for the CALVADOS-2 hub occupancy model.

**Figure S21:**
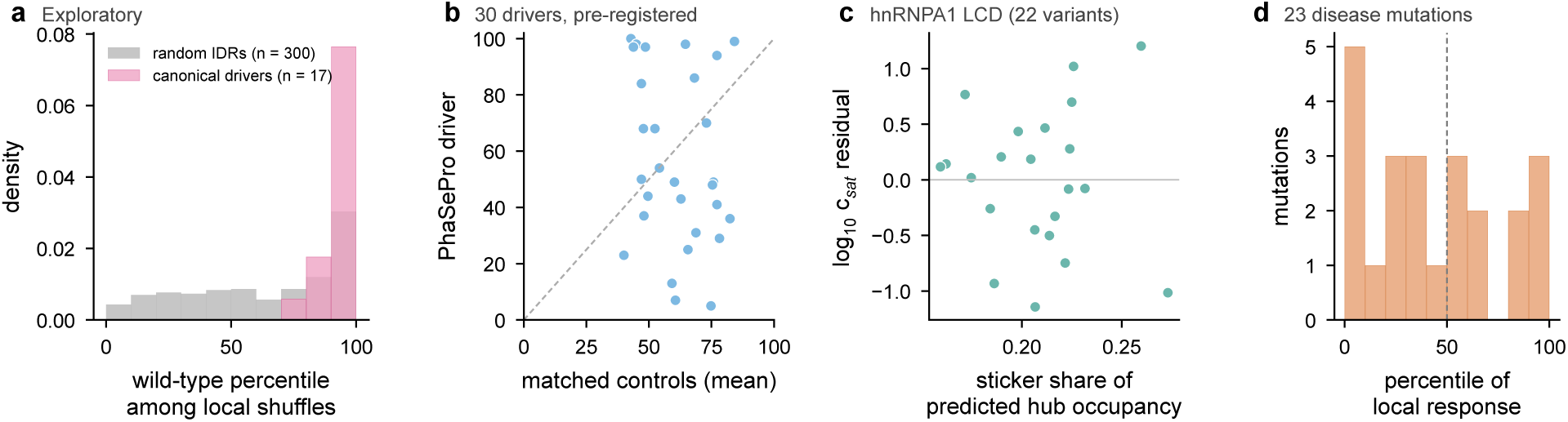
Predicted hubs and phase separation. (a) Percentile of the unevenness of the wild-type hub profile among sequences shuffled within windows of 25 residues, for 17 canonical drivers of phase separation and 300 random IDRs of the IDRome (exploratory). (b) The same percentile for 30 PhaSePro drivers against the mean of five matched control IDRs, with the diagonal dashed (pre-registered). (c) Share of predicted hub occupancy carried by aromatic residues and arginine against the residual of log_10_ saturation concentration after a baseline of sticker number and net charge, for variants of the hnRNPA1 low-complexity domain (pre-registered). (d) Percentile of the predicted local response of 23 disease mutations among the same substitution at positions with the same wild-type residue (pre-registered).

#### Sequence models of degree

The analyses of Supplementary Section S5 were repeated with degree as the target, with the same data, folds, features and booster (Table S18). Degree is predicted more accurately than hub occupancy for 9,446 of the 10,281 CALVADOS-2 proteins, by a median of 0.036, and depends less on chain position. No split-half reliability of degree is stored with the labels, so no noise ceiling is given for degree. A description of the sticker and spacer surroundings of each residue recovers as large a share of the full model for degree as for hub occupancy (0.867 of 0.934 in CALVADOS-2, 0.837 of 0.907 in Mpipi-Recharged and 0.722 of 0.925 in Mpipi-GG), and predicts the 1,791 simulated shuffles at 0.858. The surroundings that matter shift with the readout. In grouped permutation importance, the glycine and serine fraction of the surrounding window matters about twice as much for degree as for hub occupancy in all three force fields (a drop of 0.205 against 0.109 in CALVADOS-2, 0.200 against 0.131 in Mpipi-GG and 0.148 against 0.082 in Mpipi-Recharged), whereas the stickiness of the window matters about equally (0.129 against 0.154, 0.064 against 0.053 and 0.254 against 0.244). Residues of high position-adjusted degree have 34 to 58% more glycine among neighbouring residues in the three force fields, about as much as residues of high hub occupancy (31 to 58%), so the shift in importance concerns how the model uses spacer content rather than which residues surround the top residues (Supplementary Fig. S22).

#### Degree in ensembles built from experimental data

The PED tests of Supplementary Sections S8 and S9 were repeated with degree under a plan hashed before any degree prediction was made for these ensembles. The ensembles, the subsampling of models and the residue-neutral null pools were exactly those of the occupancy tests, which we confirmed by reproducing the stored occupancy labels to the last digit for one ensemble of each set, and the predictions came from the frozen CALVADOS-2 degree model. Since these ensembles had been analysed for hub occupancy before, this is a pre-registered reanalysis rather than a blind test (Table S19). Both decision rules are met with degree, including that of the second blind set, which hub occupancy missed. The paired differences between the two readouts are not significant, with degree higher in 11 of 18 development ensembles and 7 of 11 blind ensembles.

In the FUS N-terminal domain the frozen degree model agrees with the experimental degree at 0.74 (partial 0.76), against 0.54 for hub occupancy. The gain lies among the more connected residues, where degree reaches 0.66 and hub occupancy 0.42 in the upper half of the experimental profile, whereas the lower half favours hub occupancy slightly (0.47 against 0.38). With 551 models and a 10% threshold, 67% of the experimental occupancy values of this ensemble are tied, so hub occupancy resolves the most central residues poorly, while degree grades these residues.

**Figure S22:**
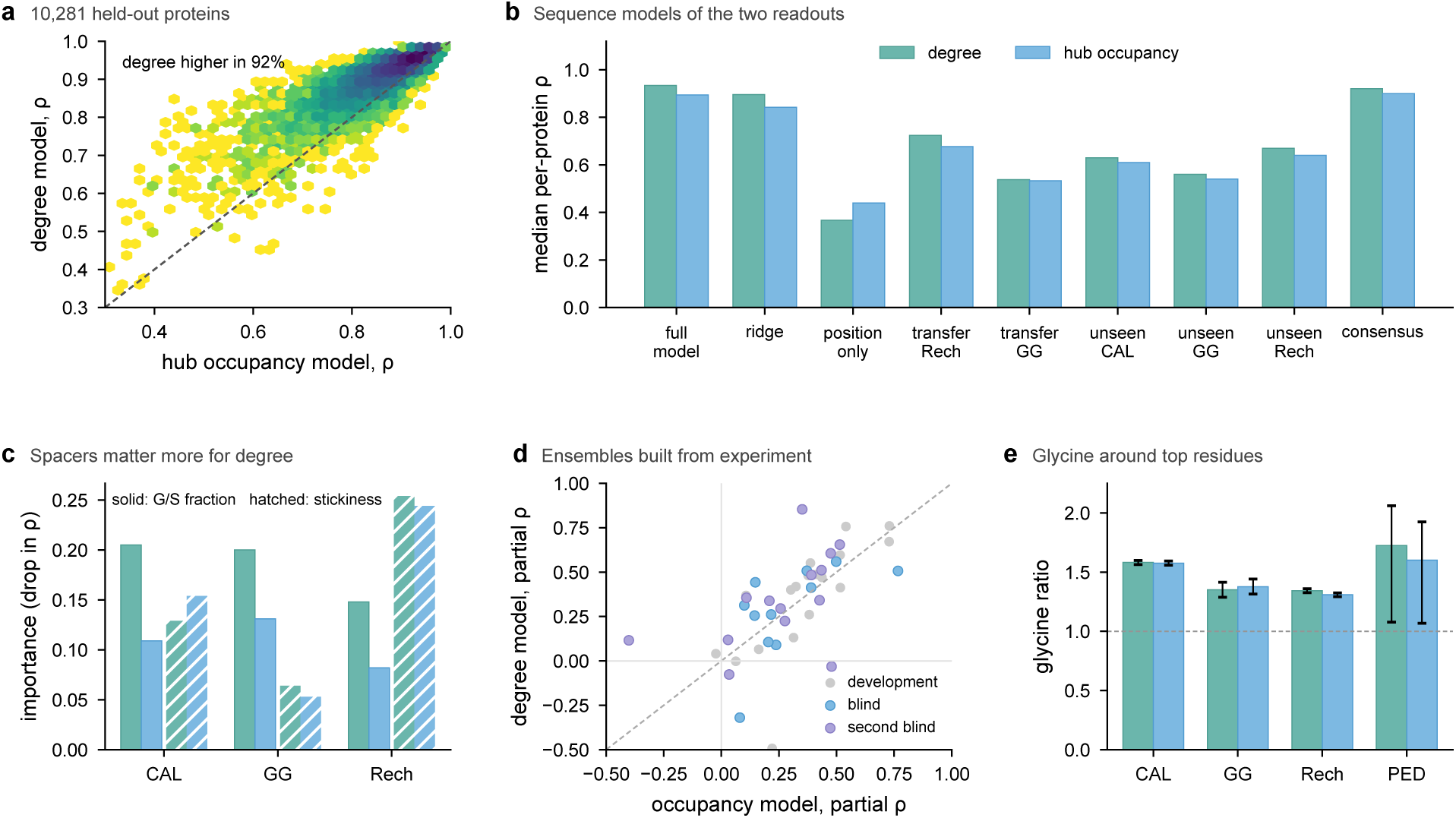
Contact degree beside hub occupancy. (a) Held-out per-protein Spearman correlation of the degree model against that of the hub occupancy model for 10,281 CALVADOS-2 proteins, with the diagonal dashed. (b) Median per-protein correlation of the sequence models of degree (teal) and hub occupancy (blue) for the full model, ridge regression and chain position on held-out CALVADOS-2 proteins, transfer to Mpipi-Recharged and Mpipi-GG, an unseen force field and the consensus model. (c) Grouped permutation importance of the sticker and spacer context model for the glycine and serine fraction (solid) and the stickiness (hatched) of the surrounding window. (d) Partial correlation given the distance from the chain ends of the frozen degree model against the frozen occupancy model on the development, blind and second blind ensembles built from experimental data. (e) Glycine among the neighbours of the top 10% of residues by position-adjusted degree (teal) or hub occupancy (blue), relative to all residues, with 95% bootstrap intervals.

#### Arrangement test with degree

An amendment to the arrangement test, hashed before any degree value of the shuffled or tyrosine-to-phenylalanine variants was read (arrangement_test_degree_plan.md), repeated the test with degree under the same decision rule. A new arrangement erases degree as completely as hub occupancy, with a median position-removed agreement of 0.11 between natural and shuffled sequences against 0.12 for hub occupancy, whereas the tyrosine-to-phenylalanine substitution and a change of force field preserve degree even more, at 0.95 against 0.87 and, between CALVADOS-2 and Mpipi-Recharged on the same proteins, 0.72 against 0.63, so the rule is met (Fig. 3a). Residue identity explains 27.3% of the within-protein variance of position-adjusted degree in natural sequences and 28.4% in the shuffled sequences (difference 1.1 points, 95% bootstrap interval 0.2 to 1.8), and natural degree profiles are more uneven, smoother and reach higher maxima than the shuffled ones, with the natural value above all three shuffles in 38%, 36% and 41% of proteins against 25% expected (*P <* 10*^−^*^8^ in each case). The frozen degree models predict the shuffles, which neither model saw, at medians of 0.93 for the CALVADOS-2 model and 0.83 for the consensus model, as well as the natural sequences of the training set (0.94 and 0.85). Without a residue-neutral null, which was computed only for betweenness, the residue-type means of degree follow the interaction scale of Mpipi-Recharged closely (*r* = 0.89) but those of CALVADOS-2 and Mpipi-GG only weakly (*r* = 0.20 and 0.18), as do the residue-type means of hub occupancy without the null (*r* = 0.30, 0.30 and 0.92), so Fig. 3b–d shows the null-corrected hub occupancy.

### S17 Phosphorylation

#### Design

The plan, the amendments of the plan, the site selection, the frozen reweighting predictions and every analysis script were hashed before the corresponding simulations. From the 83 proteins of the BENDER phosphosite set with at most 200 residues, a protein qualified if one UniProt-annotated phosphosite had an unannotated serine or threonine of the same type at least four residues from every annotated site whose log effective sample fraction (log ESS) under the charge model differed by at most 0.10, together with a second unannotated site of the same type whose log ESS exceeded that of the annotated site by at least 0.20. Of 26 qualifying proteins, 20 were drawn with a fixed seed. Each protein was simulated unmodified and with each of the three sites phosphorylated, in three independent replicates, giving 240 runs per force field with the BENDER protocol at 300 K and 150 mM. All 240 runs of each force field passed checks of sequence, phosphoresidue position, frame count and chain integrity, and the unmodified replicates match the stored BENDER runs of the same proteins about as closely as the replicates match each other.

#### Rewiring

For a site *s* with phosphorylated replicates *P*_1_ to *P*_3_ and unmodified replicates *W*_1_ to *W*_3_, and *R*(*X, Y*) the summed absolute difference of the per-residue profiles of two runs, the rewiring is 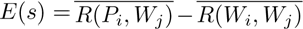, the mean over the nine mixed pairs minus the mean over the three unmodified pairs. The decisions used hub occupancy, and degree was reported alongside (Table S20). In both force fields annotated sites rewire the network no more than matched unannotated sites, so annotation adds nothing beyond the fluctuation of the potential at the site.

#### The fluctuation in Mpipi-Recharged

Since the Mpipi-Recharged phosphate changes the short-range attraction of the residue as well as the charge, a fluctuation measure native to that force field was computed before the final analysis. For every snapshot of the three unmodified replicates, the energy change of phosphorylating the site was evaluated with OpenMM on systems that serialize identically to those of the simulations, and the snapshots were weighted accordingly. Rewiring follows this fluctuation (*ρ* = −0.48 for hub occupancy and −0.62 for degree over the 60 sites, one-sided *P* = 0.005 and 0.006 when the fluctuation is permuted only among the three sites of each protein, post hoc, since the 60 sites come from 20 proteins), the low-fluctuation site is the calmest site in Mpipi-Recharged as well, in 18 of 20 proteins against both other sites, the fluctuations of the two force fields correlate at 0.68, and reweighting reproduces the direct change in radius of gyration at 0.75, with the same sign at 56 of 60 sites. Across the two force fields the rewiring of a site agrees at 0.65 for hub occupancy and 0.72 for degree (Supplementary Fig. S23).

#### Sequence controls

A linear model of the log ESS on the numbers of positive and negative residues in six bins of sequence separation, the distances to the chain ends, the length, the fraction and net charge of charged residues and the residue type was fitted to 1,735 sites of the 63 phosphosite proteins outside the direct set (protein-grouped cross-validated *R*^2^ = 0.24). On the 60 direct sites, the ensemble fluctuation predicts the rewiring scaled by the replicate noise beyond this model (permutation of the residual within proteins, *P* = 0.0007 for hub occupancy and 0.0002 for degree), and beyond the counts of positive and negative residues within five positions (*P* = 5 × 10*^−^*^5^), whereas the model in turn adds a smaller part beyond the ensemble (*P* = 0.008). The change in sequence charge decoration, examined after these tests, does not track local rewiring (within-protein *ρ* = −0.10), and neither does the change in the charge patterning parameter *κ* [11], with the site counted as −1 or −2 (*ρ* = −0.04 to −0.12, post hoc), while the fluctuation predicts rewiring beyond *κ* (*P <* 10*^−^*^4^ in a within-protein permutation test).

#### Reweighting

Supplementary Fig. S24 shows how the weights are formed and used for one protein of the direct set. In CALVADOS-2, the reweighted redistribution of hub occupancy correlates with the direct rewiring at 0.575 over the 60 sites, the reweighted and direct change profiles of individual sites agree at medians of 0.28 for hub occupancy and 0.34 for degree, and the reweighted change in radius of gyration agrees with the direct one at 0.82, with the same sign at 55 of 60 sites. For several phosphates at once, the weights concentrate on few snapshots when the phosphates are close, so the effective sample fraction falls to 1 to 2% for the three adjacent CK2 sites of HMGA1a, and direct simulation is then needed.

**Figure S23:**
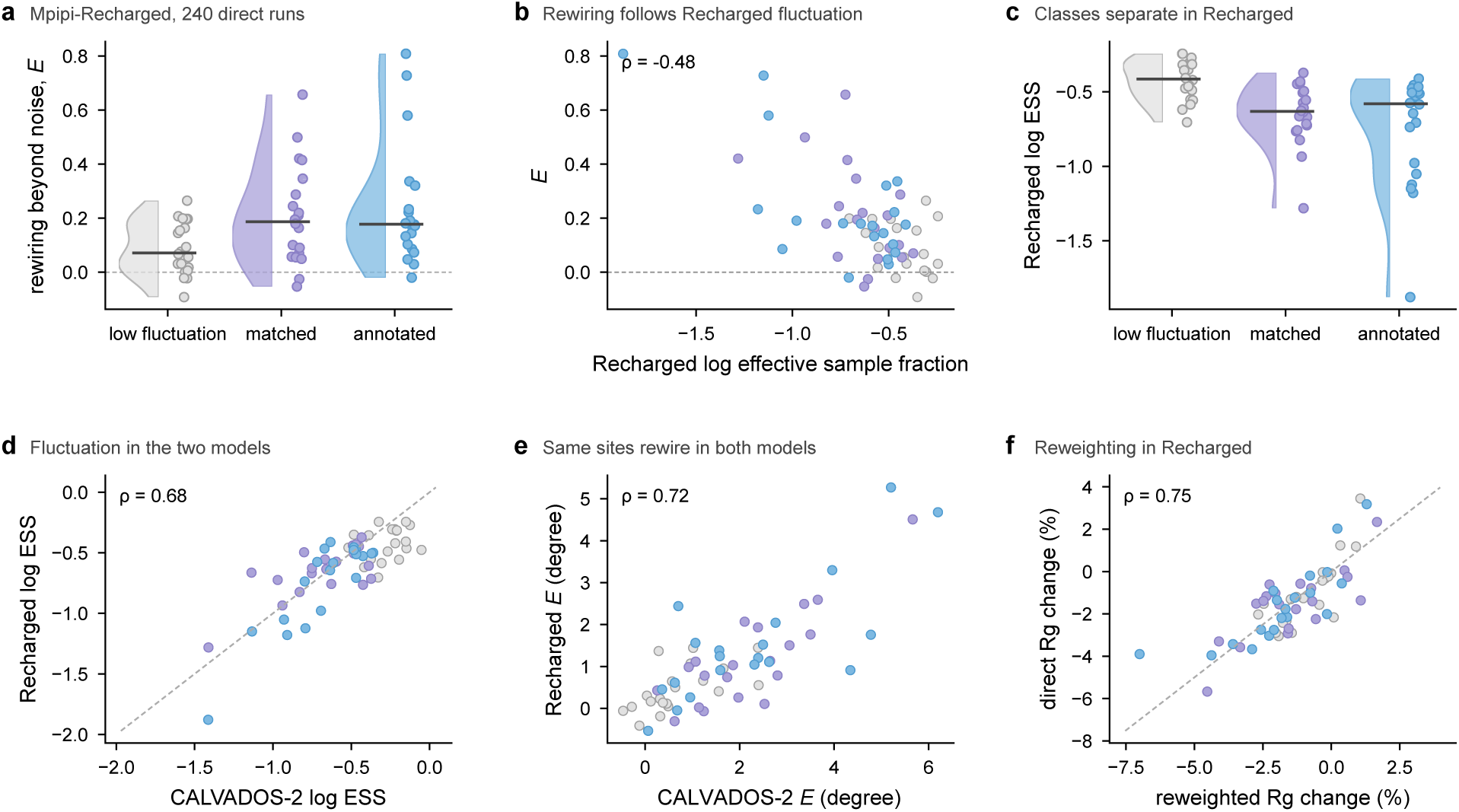
The phosphorylation result in Mpipi-Recharged. (a) Rewiring *E* of hub occupancy by site class in 240 direct Mpipi-Recharged simulations of the 20 proteins of Fig. 5, with each site as a point and the median as a line. (b) *E* against the effective sample fraction of weights from the full Mpipi-Recharged energy change of phosphorylation. (c) This effective sample fraction by site class. (d) Effective sample fractions of the two force fields. (e) Rewiring of degree in the two force fields. (f) Reweighted against direct change in radius of gyration in Mpipi-Recharged.

#### NMR case series

From the BMRB entries of disordered proteins we took every pair of an unmodified and a phosphorylated entry of the same construct measured under the same conditions whose phosphorylated serines or threonines are named (Table S21), excluding a tyrosine-phosphorylated construct and a construct whose phosphosites are not named. Both forms were simulated in three replicates under the temperature, pH and ionic strength of each sample. The amide chemical shift change is 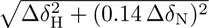 after subtracting the median change of each nucleus, and residues within three positions of a site were excluded. The pre-registered statistic is the partial Spearman correlation between the predicted contact rewiring of each residue and the measured shift change, given the logarithm of the distance to the nearest site, with a one-sided Wilcoxon test over proteins. The predicted change tracks the measured one in five of six proteins (*P* = 0.031), the change in degree in all six (*P* = 0.016), although the p53 value (+0.003) is indistinguishable from zero, the change in hub occupancy in three, and the contact change reweighted from the unmodified runs in four (*P* = 0.16). Distance from the site alone correlates with the shift change at 0.41 to 0.70 in five of the proteins, and the shifts of single phosphorylation change little, as in PAGE4, where only 8 of 83 analysed residues move by more than 0.01 ppm and where a prediction recorded beforehand that the basic N-motif would change most was not confirmed (block test *P* = 0.07).

Of the ten proteins, unmodified Sic1, a fragment of the p53 transactivation domain (residues 20 to 70) and *α*-synuclein are among the proteins whose radius of gyration was used to parameterise CALVADOS [4], and *α*-synuclein also has PRE data in that set. No phosphorylated state or chemical shift entered any force field, and Sic1 and p53 agree no better than the others.

#### Replication in Mpipi-Recharged

An amendment hashed before any Mpipi-Recharged run of these constructs (phospho_nmr_amend1_recharged.md) repeated the analysis with 36 Mpipi-Recharged runs under the conditions of each sample, without pH, which Mpipi-Recharged does not represent. The predicted change tracks the measured shifts beyond distance in all six proteins (partial *ρ* from +0.03 in H1.0 to +0.49 in the NHE1 tail, one-sided Wilcoxon *P* = 0.016), so the pre-registered reading is replicated. The change in degree is positive in five of six proteins (*P* = 0.08) and the change in hub occupancy in four (*P* = 0.34). The per-residue predictions of the two force fields agree at a median Spearman correlation of 0.57, from 0.92 in HMGA1a to −0.11 in p53.

**Figure S24:**
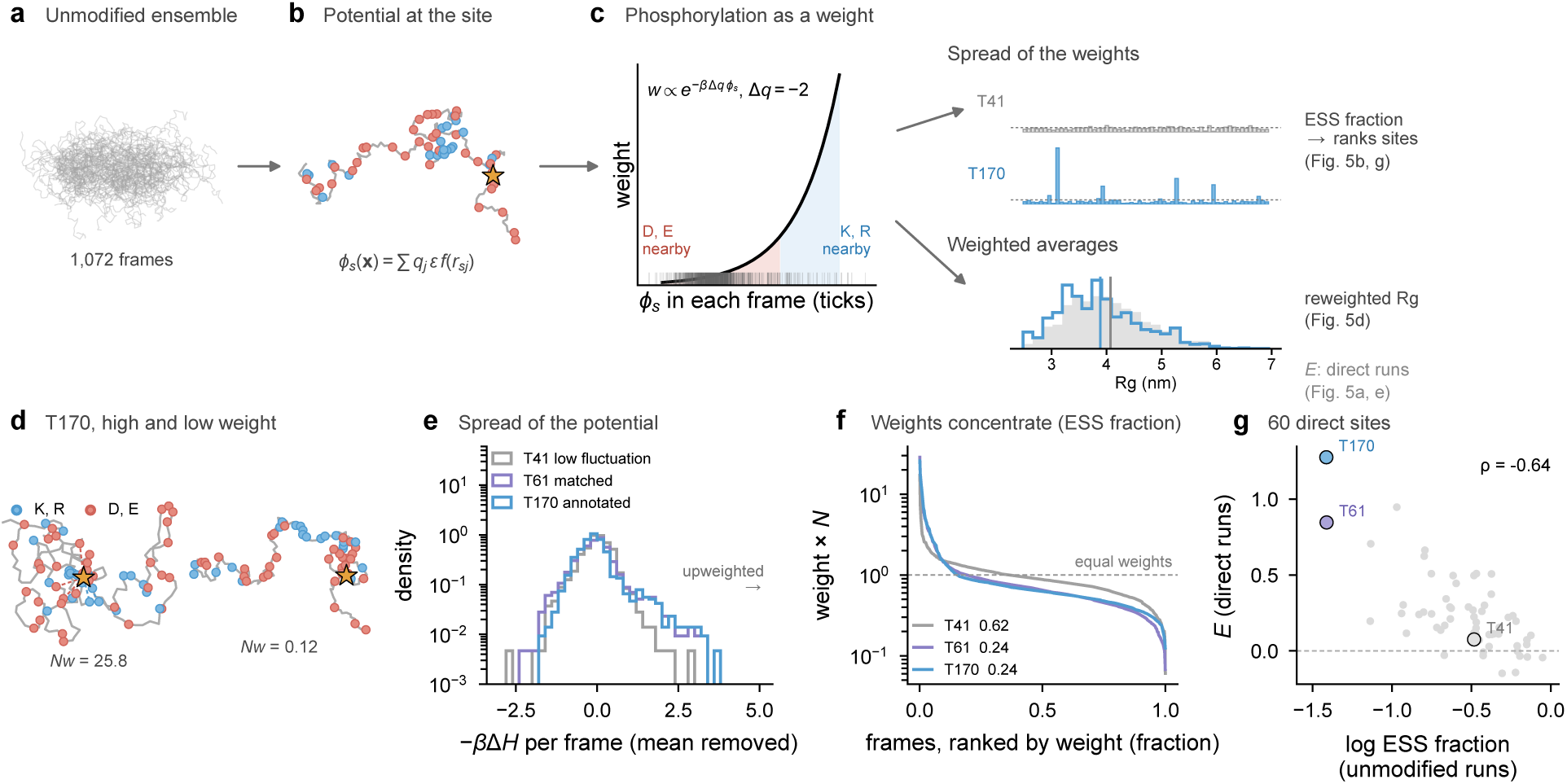
How the charge-model weights are formed and used, for the protein P28667 of the direct set. (a) Unmodified CALVADOS-2 ensemble, 30 of 1,072 frames. (b) One frame with the annotated site T170 (star), lysines and arginines (blue) and aspartates and glutamates (red), and the electrostatic potential *ϕ_s_* at the site. (c) Weight of a frame against the potential at the site, *w* ∝ *e^−β^*^Δ^*^q^ ^ϕs^* with Δ*q* = −2, and the potential in each frame of the ensemble (ticks). The spread of the weights gives the effective sample fraction, which ranks sites by rewiring, and weighted averages estimate properties of the phosphorylated form, such as the radius of gyration (grey before and blue after reweighting). Rewiring *E* is measured separately, in direct simulations of both forms. (d) Frames of highest and lowest weight for T170, with the weight relative to equal weights. (e) Distribution of −*β*Δ*H* over the frames for the three sites, with the mean removed, since a constant shift cancels on normalisation. (f) Weights ranked from largest to smallest, with the effective sample fraction of each site. (g) Effective sample fraction against directly measured rewiring for the 60 sites of Fig. 5b, with the three sites of P28667 marked.

#### Four further pairs including phosphotyrosine

A second amendment (phospho_nmr_amend2_new_ pairs.md), hashed before the shift lists of the new entries were downloaded and before any run, added every further pair found in BMRB of a disordered chain phosphorylated on that chain, with an unmodified entry of the same construct from the same study and named phosphorylated residues, *α*-synuclein phosphorylated at S129 or, in a Y133F/Y136F background, at Y125, the cytosolic domain of the B-cell receptor subunit CD79b phosphorylated at two tyrosines by Fyn, and the C-terminal domain of the bacterial effector Tir phosphorylated at four tyrosines. CD79a was excluded because the phosphorylated entry names no site, and pairs with folded domains or bound peptides were excluded. A phosphotyrosine was represented in CALVADOS-2 as the tyrosine bead with a charge of −2 and in Mpipi-Recharged as the tyrosine bead with the doubled electrostatic term of the glutamate bead, keeping the stickiness of the aromatic ring, a change that leaves systems with phosphoserine or phosphothreonine identical. Tir was recorded at pH 6.5 and 6.8 and simulated at 6.65, and the ionic strength of the CD79b samples was estimated from the buffer. With four pairs, the primary test was the mean partial correlation against independent circular shifts within each pair. The prediction tracks the shifts in all four pairs in both force fields, with a mean partial *ρ* of 0.23 in CALVADOS-2 and 0.32 in Mpipi-Recharged (*P* = 10*^−^*^4^ in each), 0.19 and 0.34 over the three tyrosine pairs (*P* = 0.005 and 10*^−^*^4^), and the result holds without Tir or without the second *α*-synuclein pair. For *α*-synuclein phosphorylated at S129, distance from the site alone does not predict the shift changes (*ρ* = −0.02), whereas the predicted rewiring does (0.37 and 0.26). The change in degree tracks the shifts in CALVADOS-2 (*P* = 0.03) but not in Mpipi-Recharged (*P* = 0.12). Over all ten pairs, the predicted change is positive in nine in CALVADOS-2 and in all ten in Mpipi-Recharged. A quality check of the Mpipi-Recharged runs first failed because the residue reader skipped the HETATM records of phosphotyrosine, and was corrected and recorded before any Mpipi-Recharged result was computed.

Supplementary Fig. S25 shows, for the other nine pairs, the predicted change in contact probability on phosphorylation next to the difference between two unmodified replicates and the measured shift changes, as Fig. 5f shows for HMGA1a. The predicted changes take three forms. In HMGA1a, the phosphorylated acidic tail gains contacts with the basic AT-hooks 20 to 45 residues away, where the largest shift changes lie. In the lysine-rich H1.0 C-terminal domain, each phosphothreonine gains contacts with neighbouring residues, whereas in the NHE1 tail the stretches around the sites lose contacts. In Sic1, PAGE4 and p53, the predicted changes are about as large as the differences between unmodified replicates (largest change 0.07, 0.05 and 0.04 against 0.06, 0.06 and 0.05), in line with the weaker agreement with the shifts in these proteins and with the single site of PAGE4 and p53 (Supplementary Table S21). The same holds in CALVADOS-2 for both *α*-synuclein pairs and for CD79b (0.05, 0.06 and 0.04 against 0.05, 0.05 and 0.06), whereas Tir changes more (0.09 against 0.06). Even where the largest changes are at the level of replicate noise, the noise-corrected per-residue prediction ranks which residues change, as the agreement with the shifts of *α*-synuclein phosphorylated at S129 shows.

#### Functional phosphosites

Whether functional scores of human phosphosites [53] track the fluctuation could not be decided, since only 21 human BENDER entries carry scored sites, and the pre-registered test on 18 proteins and 169 sites found no association (pooled within-protein *ρ* = 0.07, *P* = 0.20).

**Figure S25:**
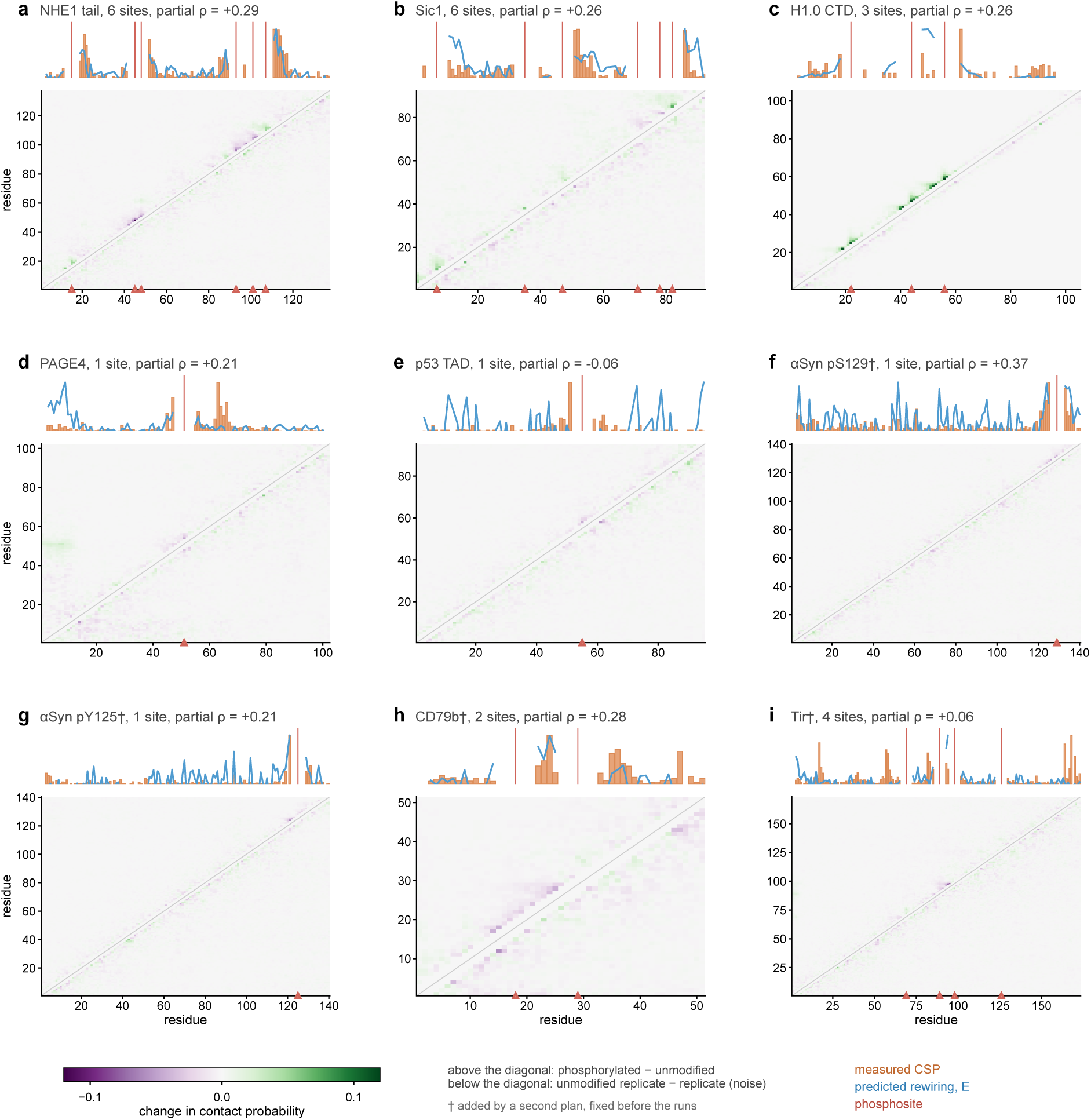
Predicted rewiring and measured shift changes for the other nine pairs of the NMR case series (HMGA1a in Fig. 5f), the five of the original plan ordered by the pre-registered partial correlation followed by the four added by the second plan (dagger). Each map shows the change in contact probability between residues on phosphorylation (mean of three phosphorylated minus mean of three unmodified CALVADOS-2 replicates) above the diagonal and the difference between the two most different unmodified replicates below the diagonal, gained in green and lost in purple, with the phosphosites marked in red. Above each map, the measured amide shift change (orange) and the predicted contact rewiring (blue) of each analysed residue, each scaled to the maximum.

**Table S5:** Enrichment of sticker and spacer sets among hubs, defined as the 10% of residues of each protein with the highest hub occupancy, relative to the share of the set among all residues, pooled over proteins, for the three force fields and the 27 ensembles built from experimental data (PED), together with the mean excess of the residues of each set over the residue-neutral null on the 907 sequences simulated with all three force fields, in units of the position-adjusted *z*-score, which averages zero over all residues. The own sets are taken from the ranking of residue types by the interaction parameter of each force field given in the text. Values in italics have 95% intervals, from 2,000 bootstrap resamples of proteins, that include 1 for the enrichment or 0 for the excess.

| Set | Residues | CALVADOS-2 | Mpipi-GG | Mpipi-Recharged | PED |
| --- | --- | --- | --- | --- | --- |
| *Enrichment among hubs* |  |  |  |  |  |
| Canonical stickers | F W Y R | 0.58 | 0.65 | 2.22 | 1.60 |
| Stickers | F W Y H | 0.58 | 1.21 | 4.20 | *1.27* |
| Aromatic residues | F W Y | 0.79 | *1.16* | 5.94 | *1.48* |
| Stickers | Y F | 0.91 | *1.04* | 5.21 | *1.32* |
| Stickers | Y R | 0.57 | 0.55 | 1.53 | 1.54 |
| Stickers | F W Y R H | 0.54 | 0.75 | 2.19 | 1.45 |
| Stickers | F W Y R K | 0.47 | 0.61 | 1.53 | 1.45 |
| Canonical spacers | G S | 2.69 | 2.10 | 1.37 | *0.84* |
| Spacers | G S N Q | 2.03 | 1.78 | 1.37 | *0.92* |
| Own four stickiest |  | 0.58 | 1.21 | 4.20 | – |
| Own five stickiest |  | 1.94 | *1.07* | 2.19 | – |
| Own six stickiest |  | 1.74 | 1.14 | 1.96 | – |
| Own stickier half |  | 1.48 | 1.19 | 1.61 | – |
| Own four least sticky |  | 0.42 | 0.81 | 0.14 | – |
| *Mean excess over the residue-neutral null* |  |  |  |  |  |
| Canonical stickers | F W Y R | *-0.00* | 0.06 | 0.51 |  |
| Stickers | F W Y H | 0.09 | 0.29 | 1.04 |  |
| Aromatic residues | F W Y | 0.19 | 0.33 | 1.45 |  |
| Stickers | Y F | 0.23 | 0.29 | 1.25 |  |
| Stickers | Y R | *-0.03* | *0.00* | 0.30 |  |
| Stickers | F W Y R H | *-0.01* | 0.09 | 0.49 |  |
| Stickers | F W Y R K | *-0.13* | *-0.01* | 0.13 |  |
| Canonical spacers | G S | 0.36 | 0.20 | 0.07 |  |
| Spacers | G S N Q | 0.24 | 0.18 | 0.13 |  |
| Own four stickiest |  | *-0.00* | 0.29 | 1.04 |  |
| Own five stickiest |  | 0.20 | 0.22 | 0.49 |  |
| Own six stickiest |  | 0.18 | 0.21 | 0.45 |  |
| Own stickier half |  | 0.12 | 0.06 | 0.27 |  |
| Own four least sticky |  | *-0.24* | *-0.06* | *-0.35* |  |

**Table S6:** Predictors trained on several force fields, as median per-protein Spearman correlation between held-out predictions and the labels of each force field. The last row gives the agreement of each simulation with the CALVADOS-2 simulation of the same sequences, which bounds a predictor that does not see the force field.

|  | CALVADOS-2 | Mpipi-GG | Mpipi-Recharged |
| --- | --- | --- | --- |
| Proteins | 10,281 | 908 | 10,266 |
| Leave one force field out | 0.61 | 0.54 | 0.64 |
| Pooled over all force fields | 0.75 | 0.77 | 0.80 |
| Consensus model | 0.79 | 0.56 | 0.85 |
| Simulation agreement with CALVADOS-2 | – | 0.53 | 0.69 |

**Table S7:**
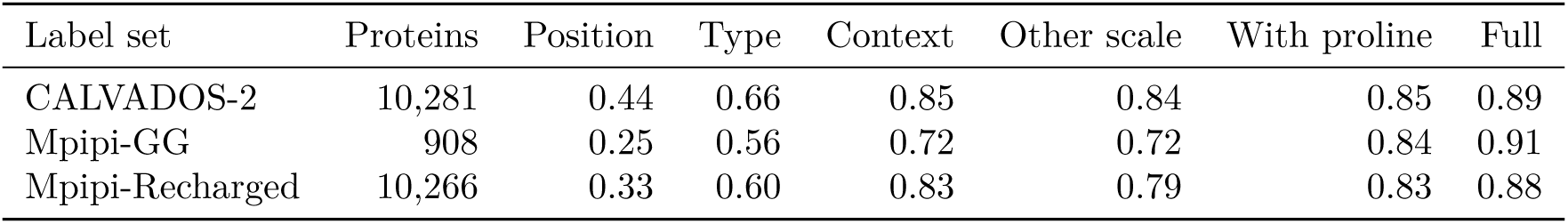
Median held-out per-protein Spearman correlation between predicted and simulated hub occupancy for models that see chain position only, together with the type of each residue, together with a description of the residue’s sticker and spacer surroundings with the stickiness scale of the same force field or of the other model family, with the proline content of the surroundings added, and for the full model with 487 sequence features.

| Label set | Proteins | Position | Type | Context | Other scale | With proline | Full |
| --- | --- | --- | --- | --- | --- | --- | --- |
| CALVADOS-2 | 10,281 | 0.44 | 0.66 | 0.85 | 0.84 | 0.85 | 0.89 |
| Mpipi-GG | 908 | 0.25 | 0.56 | 0.72 | 0.72 | 0.84 | 0.91 |
| Mpipi-Recharged | 10,266 | 0.33 | 0.60 | 0.83 | 0.79 | 0.83 | 0.88 |

**Table S8:** Composition of the ten residues around hubs, defined as the 10% of residues of each protein with the highest position-adjusted occupancy, relative to the composition around residues of the same type, for the three force fields and the 27 ensembles built from experimental data (PED). Values in italics have 95% intervals, from 1,000 bootstrap resamples of proteins, that include 1.

| Neighbours | CALVADOS-2 | Mpipi-GG | Mpipi-Recharged | PED |
| --- | --- | --- | --- | --- |
| Aromatic (F W Y) | 1.02 | *1.00* | 1.55 | *0.96* |
| R | 0.85 | *0.97* | 0.96 | 1.49 |
| K | 0.86 | 1.08 | 0.82 | *1.01* |
| Acidic (D E) | 0.93 | *0.98* | 1.02 | *0.88* |
| G | 1.30 | 1.15 | 1.18 | 1.63 |
| S | 1.15 | 1.08 | 1.02 | *0.79* |
| Polar (N Q T) | 0.94 | *1.01* | 0.95 | *0.87* |
| P | 0.95 | 0.59 | 0.93 | 0.64 |
| Aliphatic (A V I L M) | 0.96 | *0.98* | 0.89 | *1.04* |
| H | 0.90 | 1.16 | 1.09 | *0.75* |
| C | 1.06 | *1.19* | *0.99* | *1.44* |

**Table S9:**
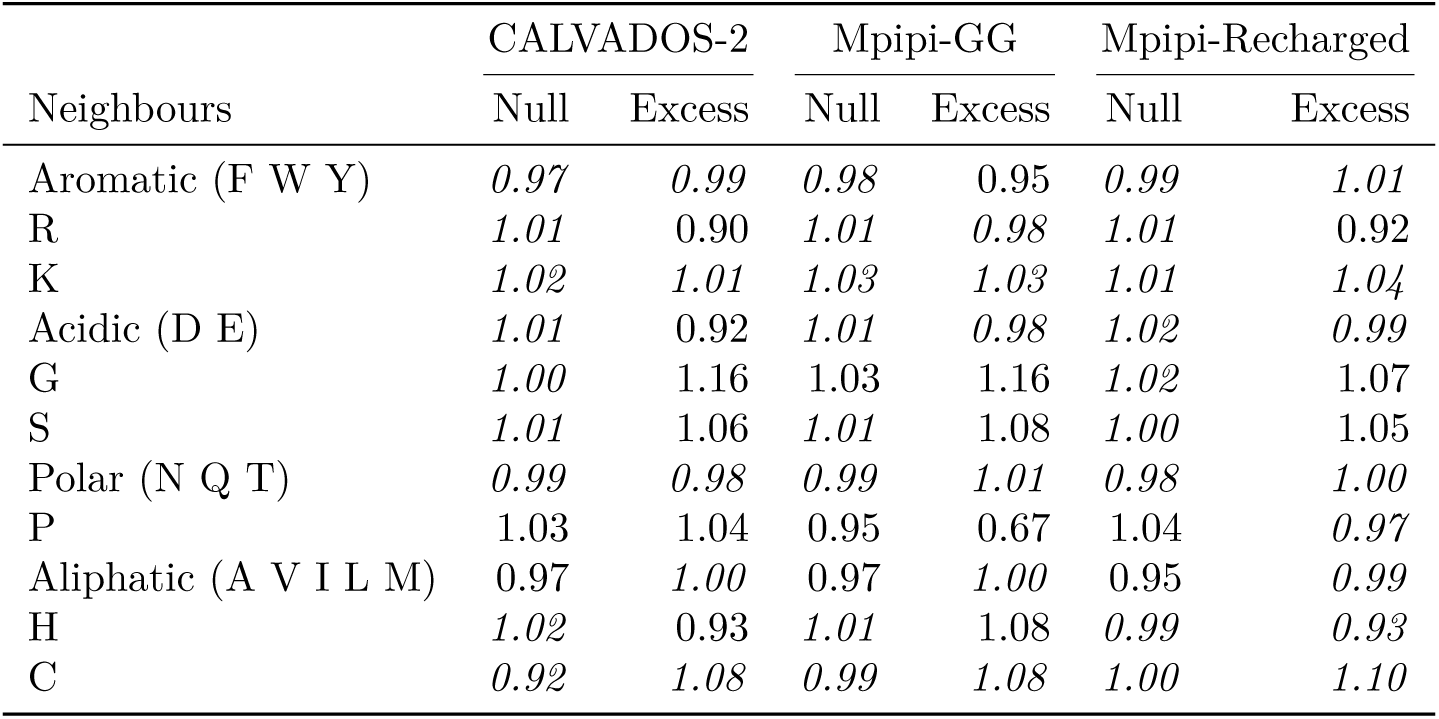
Composition of the ten residues around hubs relative to residues of the same type on the 907 sequences simulated with all three force fields, with hubs defined by the position-adjusted *z*-score of time-averaged betweenness in the residue-neutral null ensembles, which contain the bead sizes of each force field but no attraction, and by the excess of the simulations over these nulls. Values in italics have 95% bootstrap intervals over proteins that include 1.

**Table S10:** Per-ensemble Spearman correlation between predicted and ensemble hub occupancy for the consensus model, the residue-neutral null averaged over five pools and chain position alone, together with the partial correlations of the model and the null that control for distance from the chain ends. Method gives the conformer generator and the selection or refinement step that the depositors report, with data types named where given in the entry title (FM stands for flexible-meccano, MD for molecular dynamics and BME for Bayesian maximum entropy). Ensembles marked with *a* were scored after the partial correlation had been fixed as a co-primary metric.

| Entry | Protein | Method | N | Spearman |  |  | Partial |  |
| --- | --- | --- | --- | --- | --- | --- | --- | --- |
|  |  |  |  | Model | Null | Position | Model | Null |
| *Development set* |  |  |  |  |  |  |  |  |
| PED00003 | β-synuclein | MD, PRE-restrained | 134 | 0.77 | 0.32 | 0.41 | 0.73 | −0.14 |
| PED00004 | p27 KID | MD | 69 | 0.30 | 0.32 | 0.45 | −0.02 | −0.09 |
| PED00006 | $\alpha/\beta$-synuclein hybrid | MD, PRE-restrained | 145 | 0.27 | 0.27 | 0.28 | 0.11 | 0.07 |
| PED00013 | PaaA2 | ARIA, EOM | 71 | 0.38 | 0.27 | 0.32 | 0.22 | 0.10 |
| PED00022 | drkN | TraDES, ENSEMBLE | 59 | 0.45 | 0.53 | 0.78 | 0.06 | −0.24 |
| PED00023 | Sic1 | TraDES, ENSEMBLE | 92 | 0.39 | 0.15 | 0.02 | 0.39 | 0.16 |
| PED00024 | α-synuclein | MD, PRE-restrained | 140 | 0.47 | 0.39 | 0.40 | 0.30 | 0.10 |
| PED00156 | drkN | X-EISD, TraDES pool | 59 | 0.48 | 0.27 | 0.34 | 0.38 | 0.09 |
| PED00157 | drkN | X-EISD, ENSEMBLE pool | 59 | 0.51 | 0.60 | 0.80 | 0.16 | −0.09 |
| PED00159 | Sic1 | ENSEMBLE, PRE and SAXS | 92 | 0.44 | 0.25 | 0.19 | 0.44 | 0.17 |
| PED00160 | Sic1 | ENSEMBLE, PRE, SAXS and CS | 92 | 0.42 | 0.33 | 0.21 | 0.41 | 0.27 |
| PED00161 | pSic1 | ENSEMBLE, PRE and SAXS | 92 | 0.33 | 0.29 | 0.16 | 0.32 | 0.28 |
| PED00225 | eIF4G1 (1–249) | CYANA, EOM | 249 | 0.40 | 0.12 | 0.15 | 0.38 | 0.02 |
| PED00494 | FUS NTD | DEER-restrained | 270 | 0.54 | 0.30 | 0.07 | 0.54 | 0.35 |
| PED00498 | V2R C-tail | FM, RDC-refined | 33 | 0.65 | 0.59 | 0.79 | 0.31 | 0.05 |
| PED00499 | GHSR C-tail | FM, RDC-refined | 33 | 0.57 | 0.41 | 0.29 | 0.52 | 0.33 |
| PED00500 | $\beta_2$AR C-tail | FM, RDC-refined | 76 | 0.71 | 0.36 | 0.31 | 0.73 | 0.20 |
| PED00511 | PDEγ | X-PLOR | 87 | 0.53 | 0.14 | 0.16 | 0.52 | 0.06 |
| *Blind set, predictions recorded before labels* |  |  |  |  |  |  |  |  |
| PED00017 | tau fragment | FM, ASTEROIDS | 130 | 0.52 | 0.47 | 0.46 | 0.39 | 0.18 |
| PED00154 | MKK4 N-terminal region | FM, ASTEROIDS | 76 | 0.38 | 0.10 | −0.14 | 0.37 | 0.31 |
| PED00216 | CBP ID4 | FM, no selection | 207 | 0.28 | 0.12 | 0.44 | 0.08 | −0.27 |
| PED00501 | V2R C-tail, phosphomimetic | FM, RDC-refined | 33 | 0.37 | 0.32 | 0.59 | 0.50 | −0.34 |
| PED00502 | GHSR C-tail, phosphomimetic | FM, RDC-refined | 32 | 0.31 | 0.51 | 0.61 | 0.15 | 0.15 |
| PED00503 | $\beta_2$AR C-tail, phosphomimetic | FM, RDC-refined | 76 | 0.74 | 0.44 | 0.46 | 0.77 | 0.10 |
| PED00533 | Aβ40 | MD, a99SB-disp | 40 | 0.39 | 0.44 | 0.54 | 0.20 | −0.01 |
| PED00536 | ACTR | MD, a99SB-disp | 71 | 0.39 | 0.46 | 0.52 | 0.10 | 0.04 |
| PED00158<sup>a</sup> | drkN, unfolded state | X-EISD, mixed pool | 59 | 0.55 | 0.74 | 0.83 | 0.24 | 0.32 |
| PED00001<sup>a</sup> | pSic1, 11 models | TraDES, ENSEMBLE | 92 | 0.22 | 0.07 | 0.06 | 0.22 | 0.05 |
| PED00489<sup>a</sup> | 4E-BP2, 5-phosphorylated | FastFloppyTail, BME | 121 | 0.22 | 0.09 | 0.27 | 0.14 | −0.15 |

**Table S11:** Agreement in hub occupancy (Spearman correlation over residues) between independent PED ensembles of the same protein, with the method of each ensemble in parentheses and with PED00161 and PED00001 describing phosphorylated Sic1.

| Ensemble | Ensemble | Spearman |
| --- | --- | --- |
| *drkN* |  |  |
| PED00022 (TraDES, ENSEMBLE) | PED00156 (X-EISD, TraDES pool) | 0.28 |
| PED00022 (TraDES, ENSEMBLE) | PED00157 (X-EISD, ENSEMBLE pool) | 0.83 |
| PED00022 (TraDES, ENSEMBLE) | PED00158 (X-EISD, mixed pool) | 0.78 |
| PED00156 (X-EISD, TraDES pool) | PED00157 (X-EISD, ENSEMBLE pool) | 0.28 |
| PED00156 (X-EISD, TraDES pool) | PED00158 (X-EISD, mixed pool) | 0.33 |
| PED00157 (X-EISD, ENSEMBLE pool) | PED00158 (X-EISD, mixed pool) | 0.76 |
| *Sic1* |  |  |
| PED00023 (TraDES, ENSEMBLE) | PED00159 (ENSEMBLE, PRE and SAXS) | 0.16 |
| PED00023 (TraDES, ENSEMBLE) | PED00160 (ENSEMBLE, PRE, SAXS and CS) | 0.23 |
| PED00023 (TraDES, ENSEMBLE) | PED00161 (ENSEMBLE, PRE and SAXS) | 0.21 |
| PED00023 (TraDES, ENSEMBLE) | PED00001 (TraDES, ENSEMBLE) | 0.00 |
| PED00159 (ENSEMBLE, PRE and SAXS) | PED00160 (ENSEMBLE, PRE, SAXS and CS) | 0.57 |
| PED00159 (ENSEMBLE, PRE and SAXS) | PED00161 (ENSEMBLE, PRE and SAXS) | 0.61 |
| PED00159 (ENSEMBLE, PRE and SAXS) | PED00001 (TraDES, ENSEMBLE) | 0.32 |
| PED00160 (ENSEMBLE, PRE, SAXS and CS) | PED00161 (ENSEMBLE, PRE and SAXS) | 0.52 |
| PED00160 (ENSEMBLE, PRE, SAXS and CS) | PED00001 (TraDES, ENSEMBLE) | 0.30 |
| PED00161 (ENSEMBLE, PRE and SAXS) | PED00001 (TraDES, ENSEMBLE) | 0.12 |
| Median over 16 pairs |  | 0.31 |

**Table S12:**
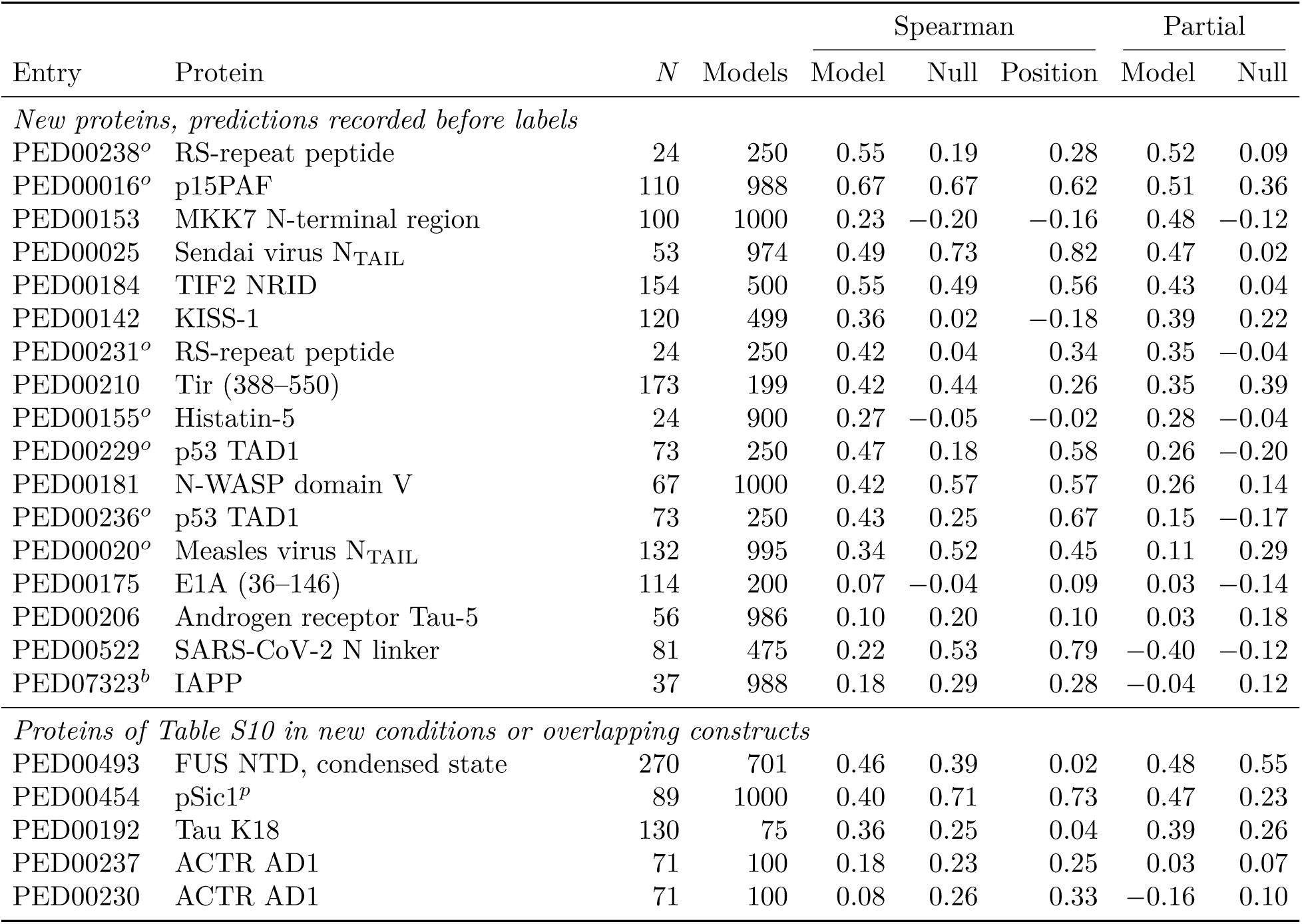
The second blind set. Per-ensemble Spearman correlation between the frozen consensus predictions and hub occupancy, for the model, the residue-neutral null averaged over five pools and chain position alone, together with the partial correlations of the model and the null that control for distance from the chain ends. Entries marked *o* have a simulated ortholog covering at least half of the construct, and IAPP, marked *b*, is excluded by the AIUPred score of the construct. The phosphorylated Sic1 ensemble, marked *p*, is scored as deposited, with 89 of 92 residues, since the deposited models omit the three phosphoserines.

**Table S13:** Agreement in hub occupancy between all-atom force fields (Spearman correlation, with the partial correlation that controls for distance from the chain ends in parentheses) and the median partial correlation of the three all-atom ensembles with the expert-built ensembles of the same protein in Table S10, with C22* for CHARMM22* and C36m for CHARMM36m.

| Protein | C22* and C36m | C22* and a99SB-disp | C36m and a99SB-disp | Expert-built |
| --- | --- | --- | --- | --- |
| Aβ40 | 0.88 (0.85) | 0.66 (0.55) | 0.68 (0.56) | – |
| ACTR | 0.74 (0.74) | 0.56 (0.51) | 0.68 (0.73) | – |
| drkN | 0.69 (0.69) | 0.41 (0.33) | 0.16 (0.12) | 0.15 |
| PaaA2 | 0.41 (0.41) | 0.43 (0.36) | 0.62 (0.65) | 0.51 |
| α-synuclein | 0.22 (0.21) | 0.23 (0.24) | 0.27 (0.18) | 0.19 |

**Table S14:** Reweighting of restraint-free pools against PRE data with two windows of excluded residues around each spin label. Agreement is the Spearman correlation between the hub occupancy of the pool and that of the expert-built restrained ensemble, before and after reweighting. Where reweighting does not improve the held-out fit, the prior weights are kept, as for *α*-synuclein with the wider window.

| Protein | Excluded residues | $\chi^2$ | | Held-out $\chi^2$<br>before / after | Effective frames | Agreement<br>before / after |
| --- | --- | --- | --- | --- | --- | --- |
|  |  | Pool | Expert |  |  |  |
| α-synuclein | $\vert i - s\vert \leq 3$ | 416 | 9.6 | 403 / 315 | 2% | 0.29 / 0.29 |
| α-synuclein | $\vert i - s\vert \leq 9$ | 7.0 | 3.2 | 7.0 / 7.0 | 100% | 0.29 / 0.29 |
| drkN | $\vert i - s\vert \leq 3$ | 1068 | 773 | 1056 / 738 | 2% | 0.69 / 0.39 |
| drkN | $\vert i - s\vert \leq 9$ | 30 | 328 | 30 / 27 | 24% | 0.69 / 0.41 |

**Table S15:**
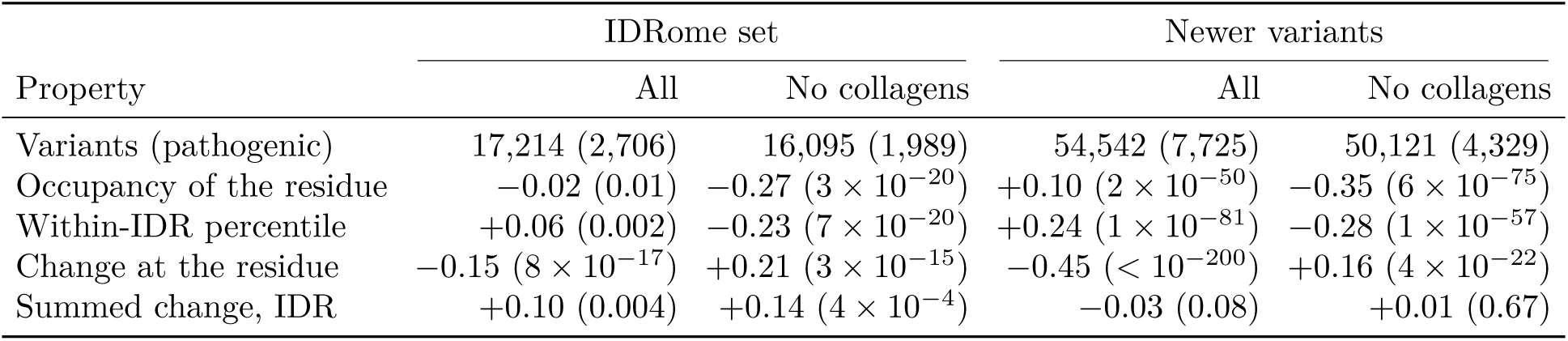
Pathogenic against benign ClinVar missense variants in human IDRs, compared as in IDRome by two-sided Brunner–Munzel tests. Values are Cohen’s *d* (pathogenic minus benign) with *P* in parentheses, for the variants mapped by IDRome and for newer variants of the ClinVar release of 5 October 2026, with and without collagens, for the hub occupancy predicted by the frozen consensus model.

**Table S16:** Simulation, prediction and experiment for the experimental constructs of the design batch. Spearman correlation with the hub occupancy of the PED ensembles of each construct (averaged over all ensembles of the same sequence) for the CALVADOS-2 and Mpipi-Recharged simulations of the construct, the prediction of the frozen consensus model and the residue-neutral null, raw and after removing the distance from the chain ends (partial). Entries marked *b* were not used during development, and the two marked *m* are unweighted molecular dynamics ensembles. Medians are computed from unrounded values and can differ by 0.01 from the median of the rounded entries.

| Construct | Entry | *N* | CALVADOS-2 |  | Mpipi-Recharged |  | Model |  | Null |
| --- | --- | --- | --- | --- | --- | --- | --- | --- | --- |
|  |  |  | Raw | Partial | Raw | Partial | Raw | Partial | Partial |
| α-synuclein | PED00024 | 140 | 0.52 | 0.37 | 0.29 | 0.14 | 0.47 | 0.30 | 0.10 |
| β-synuclein | PED00003 | 134 | 0.81 | 0.79 | 0.65 | 0.56 | 0.77 | 0.73 | -0.14 |
| $\alpha/\beta$-synuclein hybrid | PED00006 | 145 | 0.36 | 0.23 | 0.32 | 0.17 | 0.27 | 0.11 | 0.07 |
| drkN | PED00022 | 59 | 0.54 | 0.18 | 0.36 | 0.18 | 0.50 | 0.21 | -0.08 |
| Sic1 | PED00023 | 92 | 0.33 | 0.31 | 0.29 | 0.32 | 0.36 | 0.36 | 0.22 |
| pSic1 (phosphomimetic) | PED00161 | 92 | 0.40 | 0.38 | 0.31 | 0.34 | 0.38 | 0.37 | 0.23 |
| eIF4G1 (1-249) | PED00225 | 249 | 0.38 | 0.36 | 0.36 | 0.34 | 0.40 | 0.38 | 0.02 |
| V2R C-tail | PED00498 | 33 | 0.61 | -0.12 | 0.32 | -0.29 | 0.65 | 0.31 | 0.05 |
| GHSR C-tail | PED00499 | 33 | 0.67 | 0.64 | 0.53 | 0.56 | 0.57 | 0.51 | 0.33 |
| $\beta_2$AR C-tail | PED00500 | 76 | 0.63 | 0.71 | 0.68 | 0.73 | 0.71 | 0.73 | 0.20 |
| PDEγ | PED00511 | 87 | 0.47 | 0.48 | 0.61 | 0.60 | 0.53 | 0.52 | 0.06 |
| PaaA2 | PED00013 | 71 | 0.09 | -0.28 | 0.53 | 0.45 | 0.38 | 0.22 | 0.10 |
| p27 KID | PED00004 | 69 | 0.03 | -0.09 | 0.22 | 0.10 | 0.30 | -0.02 | -0.09 |
| FUS NTD | PED00494 | 270 | 0.76 | 0.77 | 0.45 | 0.45 | 0.54 | 0.54 | 0.35 |
| V2R C-tail, phosphomimetic<sup>b</sup> | PED00501 | 33 | 0.41 | 0.48 | 0.35 | 0.43 | 0.37 | 0.50 | -0.34 |
| GHSR C-tail, phosphomimetic<sup>b</sup> | PED00502 | 32 | 0.37 | 0.40 | 0.20 | 0.27 | 0.31 | 0.15 | 0.15 |
| $\beta_2$AR C-tail, phosphomimetic<sup>b</sup> | PED00503 | 76 | 0.50 | 0.64 | 0.61 | 0.75 | 0.74 | 0.77 | 0.10 |
| MKK4 N-terminal region<sup>b</sup> | PED00154 | 76 | 0.46 | 0.46 | 0.23 | 0.25 | 0.38 | 0.37 | 0.31 |
| tau fragment<sup>b</sup> | PED00017 | 130 | 0.50 | 0.39 | 0.30 | 0.27 | 0.52 | 0.39 | 0.18 |
| CBP ID4<sup>b</sup> | PED00216 | 207 | -0.02 | -0.24 | 0.13 | -0.06 | 0.28 | 0.08 | -0.27 |
| Aβ40<sup>b,m</sup> | PED00533 | 40 | 0.14 | -0.03 | 0.37 | 0.20 | 0.39 | 0.20 | -0.01 |
| ACTR<sup>b,m</sup> | PED00536 | 71 | 0.45 | 0.41 | 0.08 | -0.01 | 0.39 | 0.10 | 0.04 |
| Median of the 20 without *m* |  |  | 0.47 | 0.38 | 0.34 | 0.33 | 0.43 | 0.37 | 0.10 |

**Table S17:**
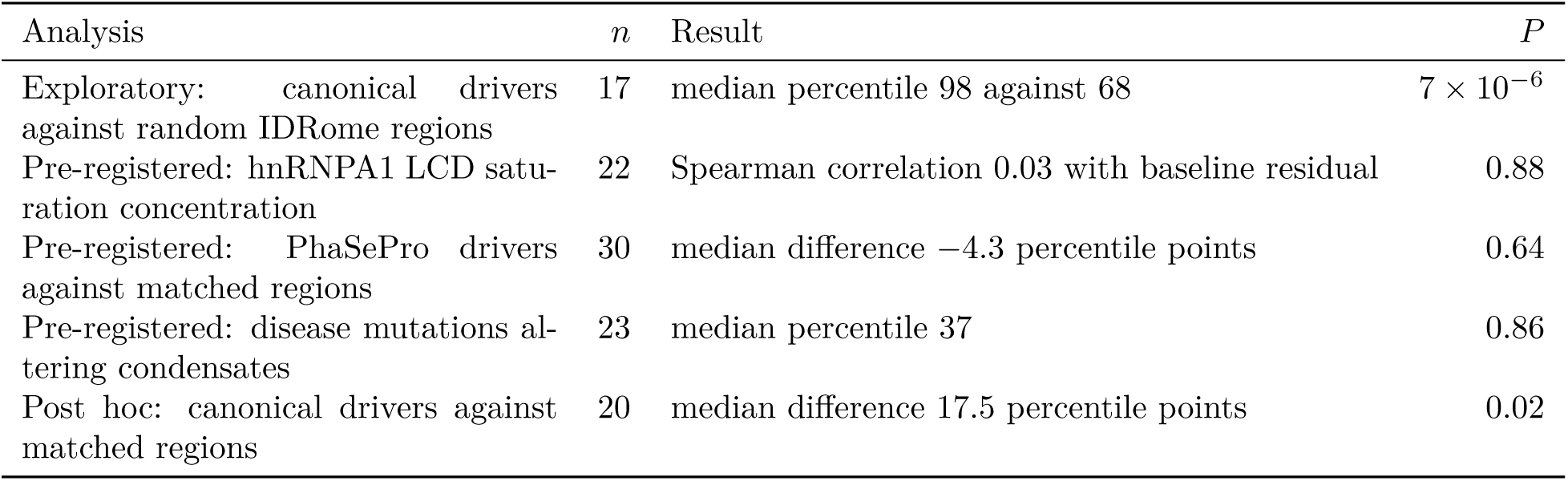
Predicted hubs and phase separation. Percentiles compare the wild type with local shuffles of the same sequence (drivers) or a disease mutation with the same substitution at positions with the same wild-type residue (mutations). The exploratory and post hoc analyses were not declared in advance.

**Table S18:** Sequence models of degree and of hub occupancy, median per-protein Spearman correlation on held-out proteins (five folds by protein).

|  | degree | hub occupancy |
| --- | --- | --- |
| CALVADOS-2, full gradient-boosted model | 0.934 | 0.894 |
| CALVADOS-2, top 10% recovered | 75% | 67% |
| CALVADOS-2, ridge on the same features | 0.896 | 0.842 |
| CALVADOS-2, chain position only | 0.367 | 0.440 |
| transfer to Mpipi-Recharged (simulations agree at) | 0.724 (0.751) | 0.677 (0.69) |
| transfer to Mpipi-GG (simulations agree at) | 0.538 (0.555) | 0.533 (0.53) |
| unseen force field, CALVADOS-2 / Mpipi-GG / Mpipi-Recharged | 0.63 / 0.56 / 0.67 | 0.61 / 0.54 / 0.64 |
| consensus model against the consensus label | 0.92 | 0.90 |
| consensus model against CALVADOS-2 / Mpipi-GG / Mpipi-Recharged | 0.83 / 0.57 / 0.88 | 0.79 / 0.56 / 0.85 |
| sticker and spacer context, CALVADOS-2 | 0.867 | 0.849 |
| simulated shuffles, context model | 0.858 | 0.845 |

**Table S19:** The frozen degree model against the degree of ensembles built from experimental data. Partial correlation given the distance from the chain ends, medians over ensembles (second blind set, over proteins).

| set | $n$ | degree model | above null | occupancy model | decision with degree |
| --- | --- | --- | --- | --- | --- |
| development | 18 | 0.42 | 17/18 | 0.38 | not blind |
| blind | 11 | 0.31 | 10/11 | 0.22 | met |
| second blind | 14 | 0.34 | 11/14 | 0.31 | met ($P = 0.03$) |

**Table S20:** Direct rewiring by site class, one-sided Wilcoxon signed-rank tests over 20 proteins (proteins with a positive difference, *P*).

| contrast | CALVADOS-2 |  | Mpipi-Recharged |  |
| --- | --- | --- | --- | --- |
|  | occupancy | degree | occupancy | degree |
| matched minus low fluctuation | 19, $3 \times 10^{-6}$ | 20, $1 \times 10^{-6}$ | 15, 0.006 | 16, 0.0007 |
| annotated minus low fluctuation | 18, $5 \times 10^{-5}$ | 19, $2 \times 10^{-6}$ | 15, 0.002 | 18, 0.0001 |
| annotated minus matched | 9, 0.43 | 10, 0.43 | 14, 0.17 | 12, 0.15 |

**Table S21:**
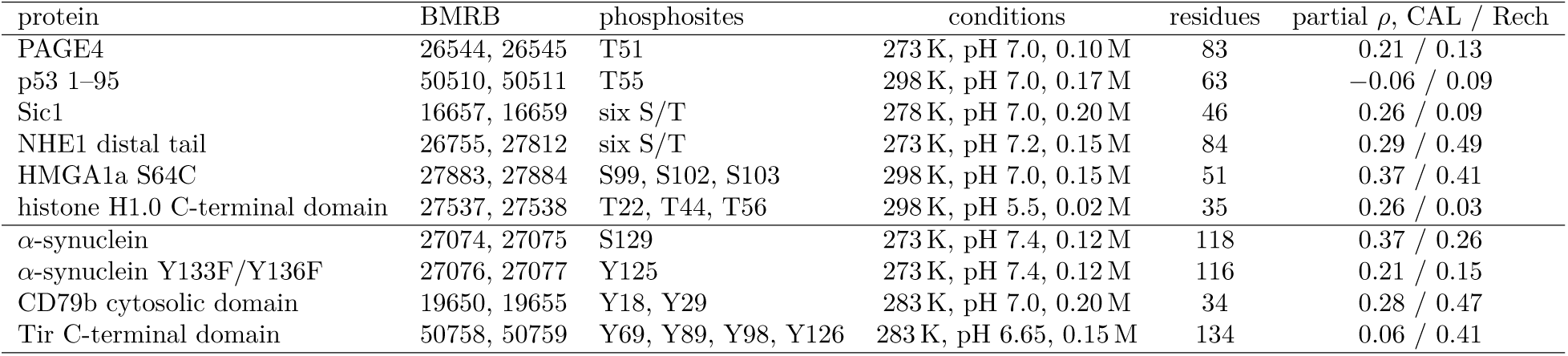
NMR case series. Partial Spearman correlation between predicted contact rewiring and measured amide shift change given the distance to the nearest phosphosite, in CALVADOS-2 and Mpipi-Recharged. The last four pairs were added by a second amendment, fixed before the runs.

